# Photoclickable HaloTag Ligands for Spatiotemporal Multiplexed Protein Labeling on Living Cells

**DOI:** 10.1101/2025.11.13.686492

**Authors:** Franziska Walterspiel, Begoña Ugarte-Uribe, Alex Cabrera, Arif Ul Maula Khan, Stefan Terjung, Claire Deo

## Abstract

Precise spatiotemporal control over fluorescence labeling is a powerful approach for selective marking and tracking of proteins of interest within living systems. Here, we report a photoclickable labeling platform based on the 2,3-diaryl-indanone epoxide (DIO) photoswitch scaffold and the self-labeling protein HaloTag. Upon illumination, the protein-bound DIO undergoes reversible photoisomerization to form a metastable oxidopyrylium ylide (PY) that reacts with ring-strained dipolarophiles via [5+2] cycloaddition, enabling covalent spatiotemporal labeling. We synthesize and characterize a library of DIO-HaloTag and DIO-SNAP-tag ligands, systematically examining the effects of linker architecture and scaffold substitution on the photoswitching and photoclick reactivity *in vitro* and on living cells. We identify a naphthyl-substituted DIO ligand exhibiting superior photoswitching and photoclick efficiency, allowing robust and selective labeling of HaloTag on the surface of living cells using visible light activation. Using this system, we achieve two- and three-color labeling of defined cell surface regions with excellent spatial and temporal precision, additionally allowing combinatorial labeling. Together, this work establishes a versatile framework for multiplexed, light-directed protein labeling compatible with living systems, with promising future applications in long-term tracking and cellular barcoding.

## INTRODUCTION

Precise spatiotemporal control of fluorescence is essential for dissecting complex biological processes in space and time, enabling selective marking and long-term tracking of proteins of interest within a biological sample.^1-3^ This can be achieved using photo-responsive fluorophores, including light-controllable fluorescent proteins or small-molecule dyes,^4-7^ targeted to proteins of interest. However, these unimolecular systems typically operate within a single spectral channel, restricting their utility to single color imaging. Extending photoactivation strategies to selectively label distinct protein subsets in different colors can offer a powerful method for long-term tracking and dynamic barcoding in complex biological environments. A particularly attractive strategy toward this goal is to control fluorophore conjugation photochemically, such that the labeling event itself is light-triggered.^8^ In principle, this can enable repeated labeling cycles with different fluorophores, yielding highly multiplexed and programmable fluorescent tagging of cellular protein targets. Several approaches have been explored using photoresponsive systems, including host–guest pairs,^9^ photocaged self-labeling tags or ligands,^10, 11^ and photoclick reactions.^12-16^ Despite these advances, achieving multicolor and spatiotemporally resolved protein labeling at the subcellular level remains challenging. Indeed, the activated species must remain localized to prevent off-target labeling by diffusion, and must either fully react or deactivate rapidly after reaction to allow sequential re-labeling. To date, only a few systems have demonstrated true sequential multicolor labeling,^9, 16^ and broadly applicable, subcellular-scale approaches compatible with live-cell imaging are still lacking.

Here, we present a photoclickable labeling system based on the self-labeling protein HaloTag,^17, 18^ enabling sequential multicolor labeling of proteins on the surface of living cells with excellent spatiotemporal control using visible light (Figure 1a). Our design is based on the 2,3-diaryl-indanone epoxide (DIO) scaffold, which undergoes reversible photoisomerization to form a metastable oxidopyrylium ylide (PY) intermediate.^19, 20^ The PY species can spontaneously react via [5+2] cycloaddition with ring-strained dipolarophiles such as bicyclononyne (BCN) and *trans*-cyclooctene (TCO) (Figure 1b).^21^ This reactivity was recently exploited for single-color fluorescence labeling on live-cell surfaces via antibody conjugation.^20^ We reasoned that integrating this photoswitchable–photoclickable platform with a broadly applicable self-labeling protein tag could yield a robust and generalizable strategy for photo-controlled protein labeling compatible with living systems. For this purpose, we rationally design and synthesize a small library of DIO-ligands of the self-labeling HaloTag^17, 18^ and SNAP-tag,^22^ and systematically investigate their reactivity *in vitro* and on the surface of living cells. We develop a naphthyl-substituted DIO ligand which undergoes efficient photoclick reaction with fluorophores on HaloTag, using visible light. With this novel system, we achieve multicolor labeling of specifically targeted regions on living cell surface, both at the cellular and subcellular level, with promising applications for dynamic imaging of protein targets.

**Figure 1.**
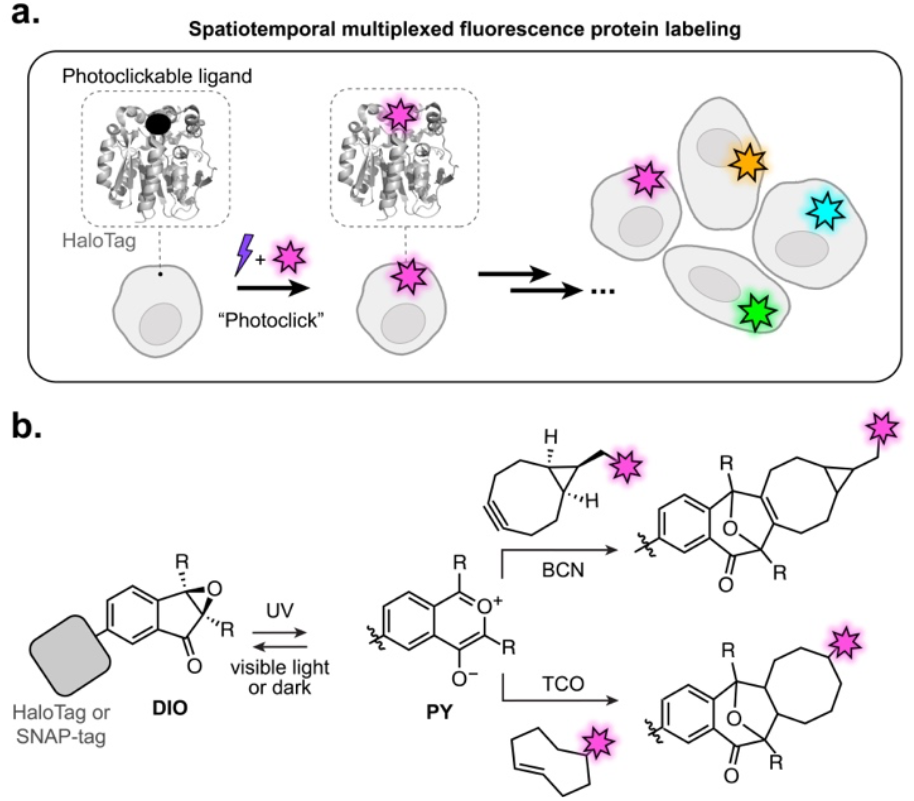
**a**. General approach for spatiotemporal multiplexed fluorescence protein labeling using photoclickable HaloTag ligands. **b**. Photoswitching and subsequent click reaction of protein-bound DIO with bicyclononyne (BCN) and *trans*-cyclooctene (TCO) attached to fluorophores, to yield fluorescently labeled proteins.

## RESULTS AND DISCUSSION

### Design and synthesis

To engineer a system for generalizable photo-controlled protein labeling, we set out to adapt the DIO-PY photoclickable scaffold to the well-established self-labeling protein tags HaloTag^17, 18^ and SNAP-tag.^22^ In particular, we reasoned that the mode of attachment to the protein would be critical for function. Indeed, the tight interaction of the protein tag with its ligand is known to substantially alter the local environment around the substrate,^23^ which in this instance could potentially impair photoswitching and/or subsequent click reaction. We therefore examined both HaloTag and SNAP-tag systems, and designed a small library of DIO ligands (Figure 2a, Figure S1). We started with the simplest, unsubstituted, DIO scaffold, functionalized at the 6-position of the indanone core with the respective protein tag ligand. For HaloTag, we assessed the influence of linker length (compounds **1**−**3**) and rigidity (compound **10**). For SNAP-tag, we compared the conventional benzyl-guanine (compound **4**) and benzyl-chloropyrimidine (compound **5**) ligands.^24^ In addition, substitution on the phenyl rings at the 2,3-position of the DIO scaffold has been shown to substantially affect their properties,^20^ and we therefore also included sterically hindered derivatives (naphthyl analogs **6** and **7**), and compounds bearing electron-withdrawing substituents (CF_3_ derivatives **8** and **9**), to fine-tune reactivity. The 6-carboxy DIO derivatives were synthesized via a palladium-catalyzed annulation from the corresponding 1,2-diphenylethynes, followed by epoxidation (Figure S2-S6). The resulting functionalized DIO intermediates were then subjected to amide coupling with the NH_2_-substituted ligands to afford compounds **1**–**10**.

**Figure 2.**
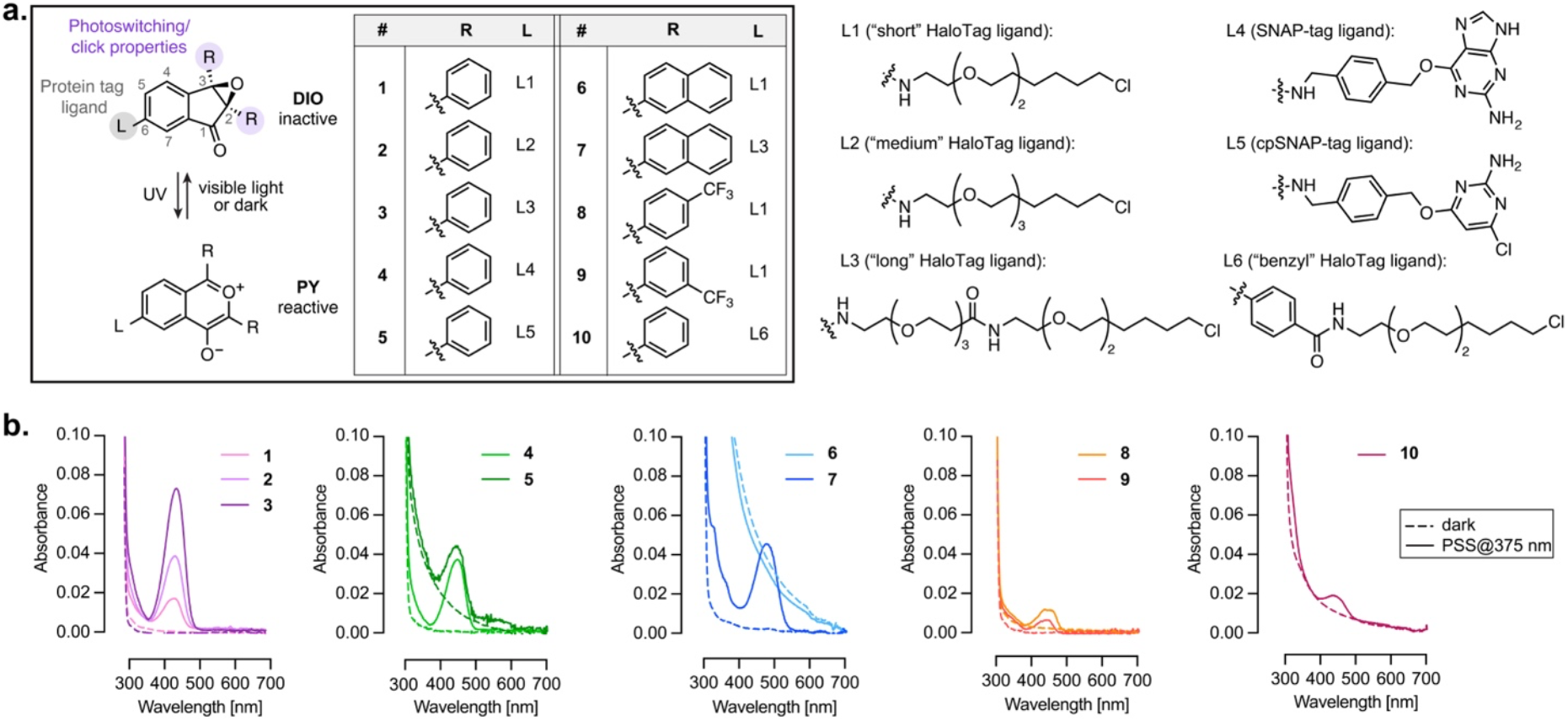
**a**. Chemical structures and photoswitching of DIO ligands **1**−**10** synthesized and studied in this work. **b**. Absorption spectra of ligands **1**−**10** bound to HaloTag or SNAP-tag in the dark (dashed lines), and at the photostationary state (PSS) reached after illumination at 375 nm (solid lines).

### Photophysical properties in vitro

This new library of DIO-based ligands was characterized *in vitro*. The DIO→PY photoswitching was induced by illumination at 375 nm (700 μW·cm^-2^), using a custom-built LED device (Figure S7).^17^ In the dark, the free ligands **1**−**10** showed no absorption in the visible range, consistent with the chemical structure of the DIO scaffold (Figure S8). Illumination at 375 nm in a PBS:MeCN (1:1) mixture led to formation of the reactive PY species, evidenced by an increase in absorption in both the UV (350–400 nm, π → π* transition) and visible regions (450–650 nm, n → π* transition). Importantly, formation of the PY isomer was fully reversible upon illumination at 545 nm (700 μW·cm^-2^), indicating efficient back-isomerization to the initial state, and allowing multiple 375 nm/545 nm switching cycles with minimal loss in performance, except for compounds **6, 8** and **9** which showed lower fatigue resistance (Figure S9). The DIO→PY photoconversion was also thermally reversible in the dark, with half-lives (t_1/2,relax_) ranging from 15 to 200 min for the free ligands at room temperature, in the same range as previously reported (Figure S9, Table S1).^20^ Compounds **1**−**5** displayed comparable photoswitching amplitude and kinetics (ε between 600 and 2200 M^-1^·cm^-1^ at the photostationary state (PSS), t_1/2,relax_ between 60 and 87 min^-1^, Table S1), indicating that the ligand itself does not significantly affect photoswitching. Compounds **6** and **7** were red-shifted by 20 nm, and exhibited substantially higher absorption at the PSS (ε = 10000 and 5400 M^-1^·cm^-1^, respectively), while relaxation kinetics were similar to compounds **1**−**5**. Introduction of CF_3_ substituents on the phenyl rings (**8**,**9**) accelerated thermal relaxation,^20^ while, in contrast, compound **10** showed markedly slower relaxation. In aqueous buffer alone, all ligands exhibited substantial aggregation (Figure S10); however, light-induced formation and disappearance of the ∼520 nm absorption band confirmed that photoswitching still occurred under these conditions.

Next, we evaluated the properties of ligands **1**−**10** covalently bound to their respective self-labeling protein tags in solution. Binding efficiency to the purified proteins was assessed via pulse-chase assay with JF_635_-HaloTag or -SNAP-tag ligands (Figure S11).^25^ Most ligands achieved complete labeling within two hours, except compounds **5, 6**, and **9**, possibly due to steric hindrance and/or aggregation of the free ligands. As expected, increasing linker length (e.g. compound **7** versus **6**) accelerated conjugation, and in all cases complete labeling was obtained after five hours at room temperature. Importantly, all protein-bound ligands retained photoswitching, as evidenced by the increase in visible absorption upon illumination at 375 nm (Figure 2b, Figure S12). Surprisingly however, all bound ligands displayed an ∼80 nm blue shift of the visible absorption (λ_max_ = 437−474 nm) compared to the free compounds, suggesting that the PY isomer interacts closely with the protein surface. This similar absorption shift observed regardless of linker length, and for both HaloTag and SNAP-tag, supports a non-residue-specific effect, potentially arising from local polarity, electrostatics or pH around the protein surface. The protein-bound PY isomers exhibited weak fluorescence (λ_em_ = 474−537 nm, Φ_F_ ≤ 0.24, Figure S13). Photoswitching amplitude varied markedly across protein-ligands conjugates (Figure 2b, Figure S12): more efficient switching was observed with increasing linker length (e.g. **3** > **2** > **1**), evidenced by larger absorption at the photostationary state. In contrast, the rigid spacer in **10** resulted in a lower turn-on. SNAP-tag ligands **4** and **5** displayed photoswitching amplitude comparable to compound **2**, likely reflecting the greater conformational flexibility of the SNAP-tag surface. Within the naphtyl series, compound **6** exhibited negligible switching, likely due to its poor binding to HaloTag, whereas **7** showed large absorption increase. CF_3_-substituted compounds **8**,**9** displayed smaller photo-responses. Overall, the protein-bound ligands exhibited slower photoswitching kinetics compared to the free ligands (Table S2), and did not display thermal relaxation over two hours (Figure S14), further evidencing the strong impact of the protein scaffold on the photoswitching properties. Nevertheless, the photoisomerization was reversible upon illumination at 450 nm (Figure S12). As increasing linker length did not abolish the protein–switch interaction, we next examined whether altering the protein structure itself could eliminate this interaction, hence restoring photoswitching behavior similar to that of the free ligand. Our previous study showed that HaloTag circularly permuted at position 178 (cp178HaloTag) retained binding to JF_635_–HaloTag ligand without fluorescence turn-on, evidencing minimal interaction with the bound fluorophore.^26^ We therefore investigated the photoswitching of compound **7** with cp178HaloTag (Figure S15). Compound **7** efficiently bound to the modified protein (Figure S15b). Upon illumination at 375 nm, photoswitching occurred, yielding two absorption bands at 460 and 561 nm, therefore partially recovering the red-shifted absorption characteristic of the free ligand. This isomerization was somewhat reversible upon illumination at 545 nm, which suggests that the reduced interaction with cp178HaloTag partially restores free-ligand-like photoswitching behavior. Overall, all compounds except compound **6** exhibited reversible photoswitching upon 375 nm/450 nm illumination when bound to HaloTag or SNAP-tag, and we next examined their potential as photoclickable protein-labeling partners.

### Photoclick reaction in vitro

The metastable PY species formed upon illumination has been shown to undergo spontaneous [5+2] cycloaddition with bicyclononyne (BCN) and *trans*-cyclooctene (TCO) derivatives, yielding a covalent adduct under physiological conditions (Figure 1b).^20^ To assess whether our protein-bound system retained similar reactivity, we illuminated the DIO–protein tag conjugates in the absence or presence of *endo*-BCN, in aqueous buffer (Figure S16). In the absence of BCN, photoswitching proceeded as described above, evidenced by the gradual increase in visible absorption around 440 nm. In contrast, no absorption increase was observed in the presence of BCN. This indicates that the species formed upon illumination rapidly reacts with BCN to form the non-absorbing covalent adduct, supporting that all the DIO–protein conjugates undergo efficient photoclick reaction under physiological conditions. Interestingly, when excess *endo*-BCN was added only after illumination (once the photostationary state had been reached), the visible absorption showed only minor decrease, suggesting very slow click reaction under these conditions (Figure S17a,b). The free ligand did not show this behavior, undergoing rapid reaction with *endo*-BCN regardless of illumination/addition order (Figure S17c,d). These observations suggest that the species reactive toward the click reaction on protein is not the long-lived form absorbing around 440 nm, but rather a transient intermediate generated during the photoswitching process.

To further evaluate the possibility to use the photoclick reaction for fluorescence protein labeling, we synthesized clickable fluorophore **11** (JF_549_-BCN, Figure 3a, Figure S18-S20), and examined its [5+2] cycloaddition with protein-bound ligands **1**−**10** by SDS–PAGE gel electrophoresis (Figure 3b-e). The DIO– protein conjugates were incubated with excess JF_549_–BCN and either kept in the dark or illuminated (375 nm, 10 min). After removal of unreacted dye and denaturation, gel electrophoresis clearly revealed light-dependent fluorescence labeling. For all HaloTag ligands (**1**−**3, 6**−**10)**, the click reaction was highly efficient, fully saturating the HaloTag as evidenced by the positive control for maximal intensity JF_549_-HaloTag ligand (Halo_549_, Figure 3b).^25^ Ligands **6** and **7** exhibited minor residual labeling in the dark, likely due to their red-shifted absorption spectra which render them more sensitive to residual ambient light during handling. In contrast, SNAP-tag ligands **4** and **5** displayed reduced photoclick efficiency, reaching only 40–50% labeling under identical conditions. As protein-bound **4** and **5** exhibited similar photoswitching as **2** (Figure 2b), this significant difference can be attributed solely to the protein scaffold, with the SNAP-tag environment partly impairing the photoclick reaction. Kinetic analysis of **3**-HaloTag revealed that the reaction reached >80% completion within 5 min under our illumination conditions, and that equimolar amount of JF_549_-BCN was sufficient for complete reaction (Figure S21a-d). Light-intensity titration indicated that photoswitching, rather than cycloaddition, was the rate-limiting step (Figure S21e,f). Overall, these results demonstrate that the photoclick reaction proceeds efficiently for protein-bound ligands *in vitro*, leading to controlled fluorescence protein labeling, with the HaloTag conjugates showing the highest reactivity.

**Figure 3.**
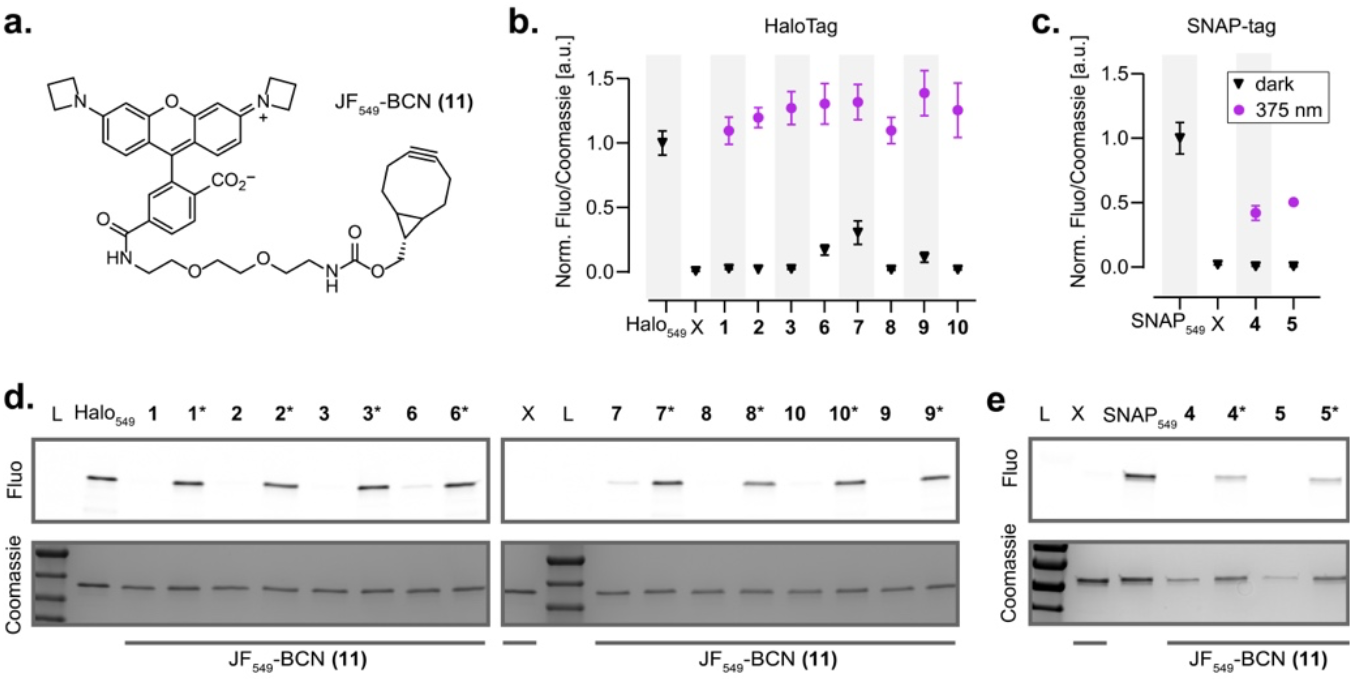
Photoclick reaction with protein-bound **1**−**10** *in vitro*. **a**. Structure of JF_549_-BCN **11. b**,**c**. Normalized JF_549_ fluorescence/Coomassie signal ratio in the dark (black triangles) and after photoclick reaction (purple dots) for protein-bound ligand **1**−**10**, quantified on the corresponding SDS-PAGE gels in **d**,**e**. Mean±SEM for three replicates. HaloTag and SNAP-tag labeled with the corresponding JF_549_ ligand (Halo_549_ and SNAP_549_, respectively) were used as positive control for maximal labeling. X: no ligand; L: protein ladder. * in **d**,**e** indicates the illuminated samples, the other samples were kept in the dark.

### Photo-controlled protein labeling on the surface of living cells

We next evaluated the use of ligands **1**−**10** in living cells, assessing protein labeling both for tags expressed extracellularly on the cell surface, or localized intracellularly in the cytosol. Similarly as *in vitro*, labeling efficiency was determined by pulse– chase assay, using JF_635_–HaloTag ligand or JF_635_-SNAP-tag ligand, to quantify remaining unbound protein after incubation with ligands **1**−**10** (Figure S22, see Methods). On the cell surface, all HaloTag ligands achieved complete labeling within two hours, evidenced by the absence of JF_635_ signal. In cells expressing cytosolic HaloTag, labeling outcomes were more variable. Ligands **1, 3, 7, 9, 10** showed >75% labeling after two hours, whereas **2, 6** and **8** produced only weak labeling, likely due to lower membrane permeability and/or solubility. SNAP-tag ligands **4** and **5** showed negligible labeling both outside and inside cells in these conditions.

Given the efficient labeling outside of cells, we next investigated the photoclick reaction on the surface of living cells expressing the self-labelling protein tag extracellularly (IgK-chain leader sequence). Living U2OS cells co-expressing a SNAP-tag−HaloTag fusion on the cell surface and EGFP in the cytosol were first labeled with DIO ligands **1**−**10**. For the clickable fluorophore, we used the commercially available, cell-impermeant CF®647-TCO, which was then added to the medium before the plates were either illuminated at 375 nm for 5 min, or kept in the dark (Figure 4a). Excess fluorophore was washed away, and click efficiency was quantified by widefield fluorescence microscopy (Figure 4b,c, Figure S23-S25). While most ligands led to only a modest increase in far-red fluorescence following illumination, ligand **7** exhibited the highest performance, yielding ∼4-fold increase in far-red fluorescence intensity after photoclick reaction. The fluorescence signal was localized on the cell surface, consistent with protein-targeted photoclick labeling (Figure 4c, Figure S23, S24). Flow cytometry confirmed these findings, showing ∼3-fold higher fluorescence signal for **7** compared to the other ligands (Figure S26, S27). In contrast, no detectable photo-induced labeling was observed when HaloTag was expressed intracellularly and reacted with the cell-permeable fluorophore JF_549_–BCN. Although the precise cause of this loss of activity inside cells is unclear, possible contributing factors include reduced stability of the PY form in the intracellular environment or heterogeneous distribution of the clickable fluorophore partner inside cells, which could be directly linked with the photoswitching behavior on protein (see above). Importantly, no cytotoxicity or phototoxicity were observed with this system (Figure S28). Overall, ligand **7** clearly outperformed all other compounds, enabling efficient, selective, and non-toxic photoclick labeling of HaloTag on the surface of living cells.

**Figure 4.**
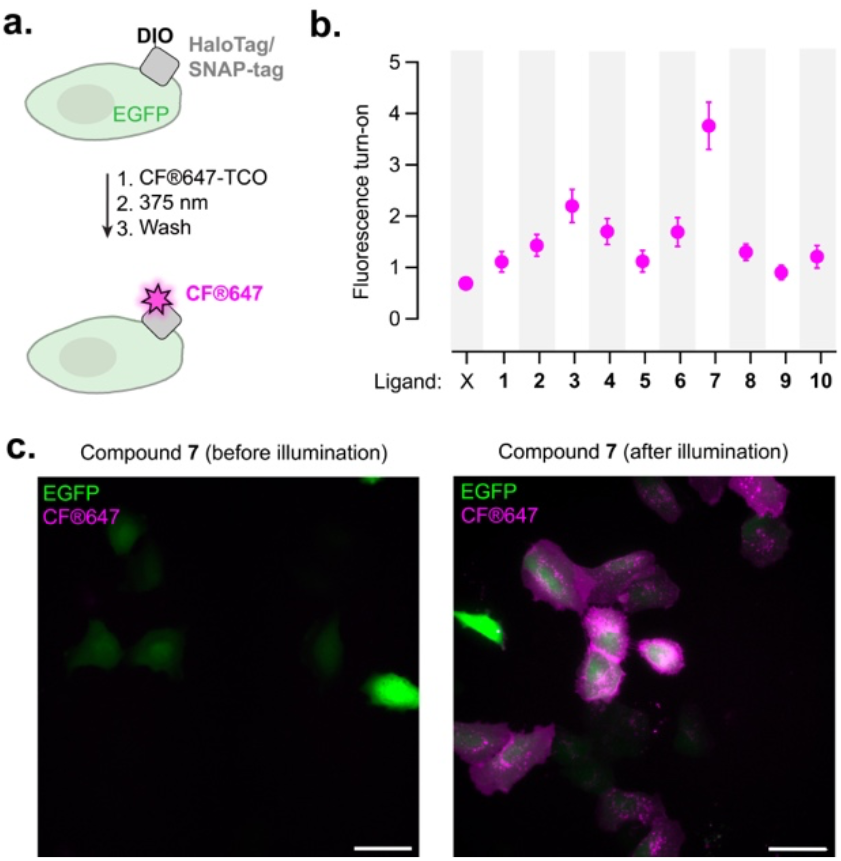
**a**. Photoclick labeling with CF®647-TCO on living U2OS cells co-expressing cell surface SNAP-tag−HaloTag fusion and cytosolic EGFP. **b**. Fluorescence turn-on of cells labeled with ligands **1**−**10** or no ligand (X) after photoclick reaction with CF®647-TCO (2 μM) induced by illumination at 375 nm for 5 minutes (700 μW·cm^-2^). Mean±SEM for N > 200 cells in each condition, two independent replicates. **c**. Representative widefield fluorescence images for compound **7** incubated with CF®647-TCO and kept in the dark (left panel) and after photoclick reaction (right panel). Scale bars: 50 μm.

The efficient photoclick reaction on HaloTag can enable selective, sequential labeling of HaloTag-expressing cells or cell-surface regions with distinct colors defined by targeted illuminated areas, thereby allowing fluorescence barcoding of spatially and temporally defined targets. This approach, however, requires that all reactive species formed during labeling with the first color are fully inactivated before introducing the second color, to prevent signal leakage into undesired regions. The DIO–PY system was only previously used in single color labeling.^20^ However, its thermal and photochemical reversibility makes it particularly well suited for sequential multicolor photo-labeling, with promising application for highly multiplexed fluorescence barcoding. We therefore evaluated the best-performing ligand **7** for two-color surface labeling of living cells (Figure 5a, Figure S29). Living U2OS cells co-expressing cell surface HaloTag and intracellular EGFP were labeled with ligand **7**. CF®647–TCO was added to the medium, and selected cell regions were illuminated for 20 seconds using the confocal laser scanning microscope at 405 nm (scanning speed, 4μs/pixel, 41 kW·cm^-2^, see Methods). After washing away the excess of dye, the photoclick process was repeated with the blue-shifted Cy3–TCO, illuminating different cellular regions. Gratifyingly, spatially defined and highly selective two-color labeling was clearly visible, with no detectable signal in non-illuminated regions. This indicates that external deactivation of potentially remaining reactive species is not necessary between cycles, simplifying the workflow, which could be due to the stabilization of a non-reactive species on HaloTag after photoswitching (see above and Figure S17) In addition, only a few seconds of illumination are required to achieve clearly visible labeling. The resulting fluorescence labeling was stable, and imaging over time showed diffusion of labeled surface proteins (Figure S30, Movie S1). By using cells not expressing EGFP, this strategy could be readily extended to three-color labeling, even within a single cell, demonstrating the versatility and precision of fluorescence labeling in defined subcellular regions (Figure 5b,c, Figure S31). Ultimately, the number of accessible unique fluorescence identifiers is limited by the multiplexing capability of the microscope. However, multiple fluorescence barcodes can still be generated using only a small number of fluorophores by performing combinatorial labeling. In practice, the labeling can be precisely controlled simply by adjusting the degree of photoactivation, defined by the microscope illumination time/power. Using this approach, we demonstrate that four cells can be distinctively labeled using only two fluorophores in two photoclick cycles (Figure S32). This combinatorial labeling could be quantified, with the relative signal intensity in the two channels showing excellent correlation with illumination time regardless of variability in expression levels across cells (Figure S32c). Although this optical barcoding strategy using a limited set of fluorophores will require system-specific calibration, these results demonstrate the proof-of-principle for highly multiplexed fluorescence barcoding of HaloTag protein targets on the surface of living cells.

**Figure 5.**
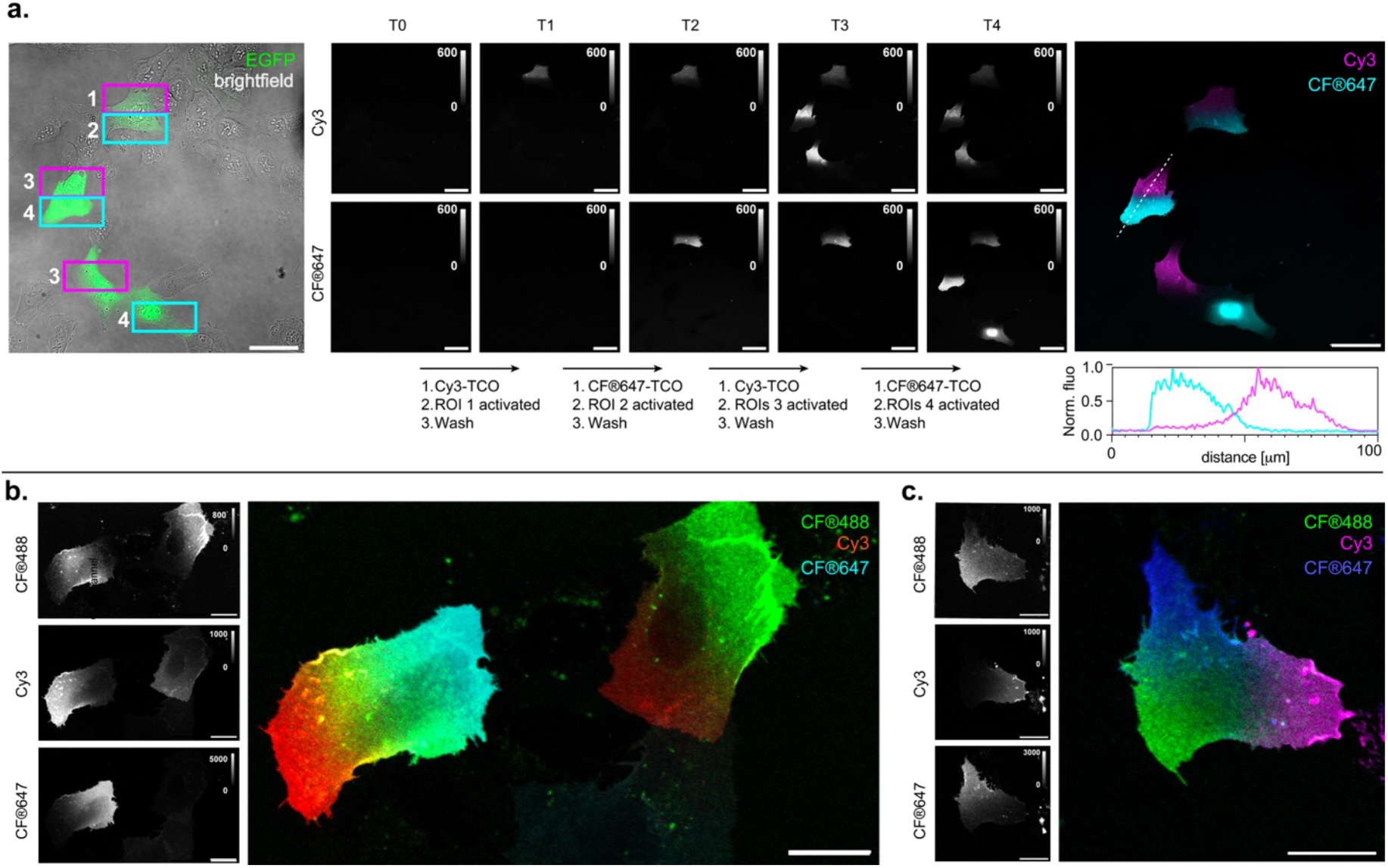
Spatiotemporal multicolor labeling of living U2OS cells expressing extracellular SNAP-tag−HaloTag fusion and labeled with ligand **7. a**. Confocal images of two-color sequential labeling. Illumination of selected ROIs was performed at 405 nm for 20 s (T1–T4) in the presence of CF®647-TCO or Cy3-TCO (2 μM) (see Methods). From left to right: fluorescence in EGFP channel with selected ROIs for the four successive cycles; fluorescence images for the Cy3 (middle top) and CF®647 (middle bottom) channels, after each photoclick cycle; overlay of the Cy3 and CF®647 channels after final cycle, with line profile quantification underneath. Representative images of five independent experiments. Scale bars: 50 μm. **b, c**. Confocal images of three-color sequential labeling with CF®488A-TCO, Cy3-TCO and CF®647-TCO. Representative images of three independent experiments. Scale bars: 20 μm.

## CONCLUSION

In conclusion, we synthesized and systematically characterized a family of DIO-based photoclickable ligands for self-labeling protein tags, enabling light-controlled protein labeling under physiological conditions. Through systematic variation of linker length, rigidity, and protein tag partner, we established structure–property relationships governing photoswitching efficiency and photoclick reactivity, and evidence the strong influence of the protein environment on these properties. Among this family of ligands, the novel naphthyl-DIO ligand **7** emerged as the most efficient, showing robust light-induced labeling of HaloTag on the surface of living cell. We demonstrate that this system enables highly specific, multicolor labeling of spatially and temporally defined cells and cell surface regions. The short illumination time required and the spontaneous deactivation of the reactive intermediate enable a rapid and streamlined labeling workflow, establishing this system as a practical tool for fluorescence barcoding. While achieving intracellular protein labeling will require further optimization of the system, the current iteration already enables precise labeling on the cell surface, with promising applications for multiplexed cell or protein tracking within a complex biological sample. Beyond fluorescence labeling, this platform could be readily applied to other clickable fluorescent partners including functional reporters such as biosensors, and extended to non-optical applications such as affinity capture handles or oligonucleotides barcodes.^27, 28^ Together, these results demonstrate that tailoring the DIO–PY system to the self-labeling HaloTag enables robust spatiotemporally controlled protein labeling on living cells, hereby expanding the toolbox for light-directed multiplexed protein tagging.

## SUPPLEMENTARY MATERIAL AND DATA AVAILABILITY

Supplementary figures and tables, methods, synthesis and characterization for all new compounds (pdf). Movie S1: Time-lapse of two-color sequential fluorescence labeling on targeted regions of living cell surface.

The image analysis code used for segmentation and intensity quantification of widefield images was adapted from previous work.^26^ It can be found along with test data at https://git.embl.org/grp-cba/htlov-mitochondria-intensity-quantification.

## ACKOWLEDGMENTS

This work was supported by the European Molecular Biology Laboratory (EMBL) and the Chan Zuckerberg Initiative (Deep Tissue Imaging grant no. 2024-337799).

The authors acknowledge: the Lavis group (Janelia Research Campus, HHMI) for sharing fluorophores; the Johnsson group (MPI for Medical Research) for sharing plasmids; Franziska Kundel and Marina Makharova (EMBL) for help with the microfluidics system; The Core Facilities at EMBL: the Electrical and Mechanical Workshops (LED illumination device), the Protein Expression and Purification Core Facility (protein purification and characterization), the Advanced Light Microscopy Facility (live cell imaging), the Bioimage Analysis Service team of EMBL Data Science Centre (image analysis), the Flow Cytometry Core Facility (flow cytometry data acquisition and analysis), the Metabolomics Core Facility (liquid chromatography-mass spectrometry data acquisition and analysis).

## COMPETING INTERESTS

The authors declare no competing interests.

### ABBREVIATIONS

BCN: bicyclononyne
DIO: 2,3-diaryl-indanone epoxide
HTL: HaloTag ligand
PSS: photostationary state
PY: oxidopyrylium ylide
STL: SNAP-tag ligand
TCO: trans-cyclooctene.

## AUTHOR CONTRIBUTION

FW: conceptualization, investigation, methodology, formal analysis, validation, visualization, writing (original draft, review & editing); BUU: investigation, methodology, writing (review & editing); AC: investigation, methodology, writing (review & editing); AUMK: software, data curation, writing (review & editing); ST: investigation, methodology, writing (review & editing); CD: funding acquisition, project administration, resources, supervision, formal analysis, visualization, writing (original draft, review & editing).

