## Supporting Information for "Photoclickable HaloTag Ligands for Spatiotemporal Multiplexed Protein Labeling on Living Cells"

### Supplementary Figures

**Figure S1**: Structures of photoclickable DIO ligands **1−10** synthesized and investigated in this work.

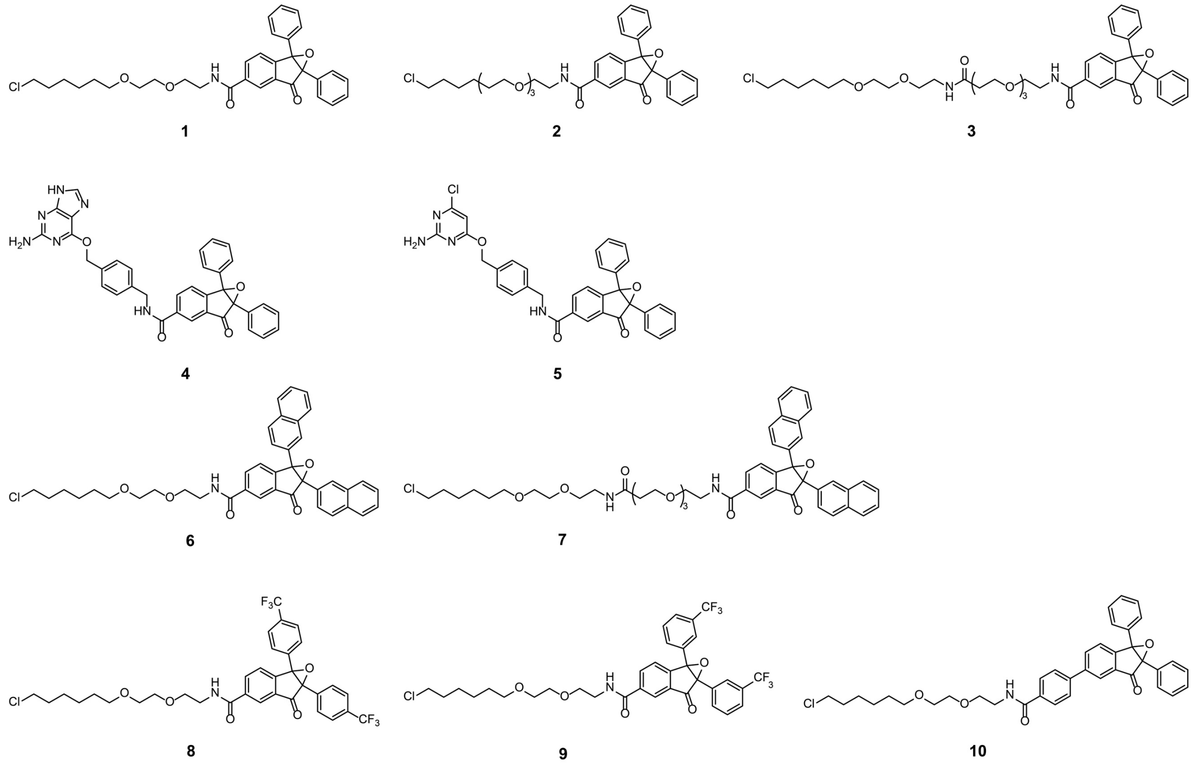

**Figure S2**: Synthesis of compounds **1−5**. **a**. Deprotection of methyl 1-oxo-2,3-diphenyl-1H-indene-6-carboxylate. **b.** Epoxidation. **c**. Amide coupling with HaloTag and SNAP-tag ligands.

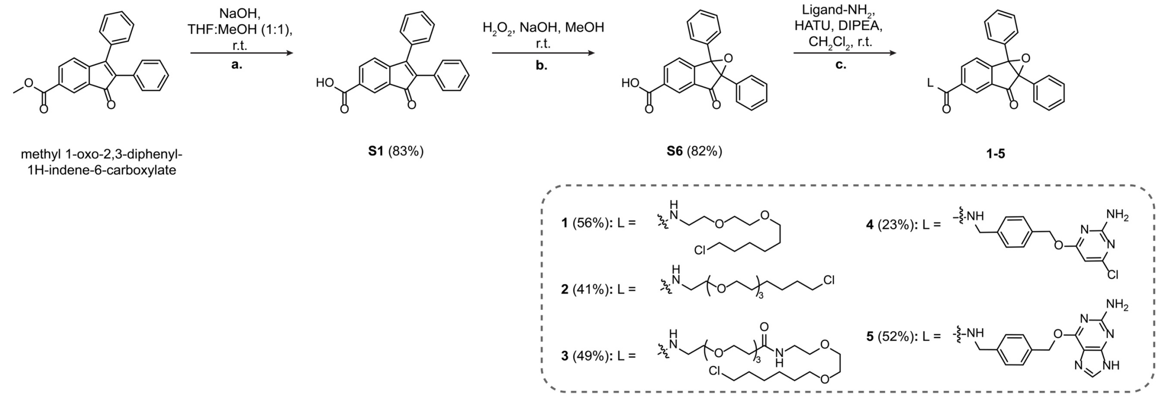

*Note:* Xie *et al.* [1] described an alternative synthetic route to the methyl ester-protected epoxides. However, we were unable to isolate the product in good yield using this method, and obtained higher yields when the carboxylic acid was deprotected prior to epoxidation.

**Figure S3**: Synthesis of compounds **6**, **7**. **a**. Palladium-catalyzed annulation of methyl 4-bromo-3-formylbenzoate and 1,2-di(naphthalen-2-yl)ethyne. **b.** Epoxidation. **c**. Amide coupling with HaloTag ligands.

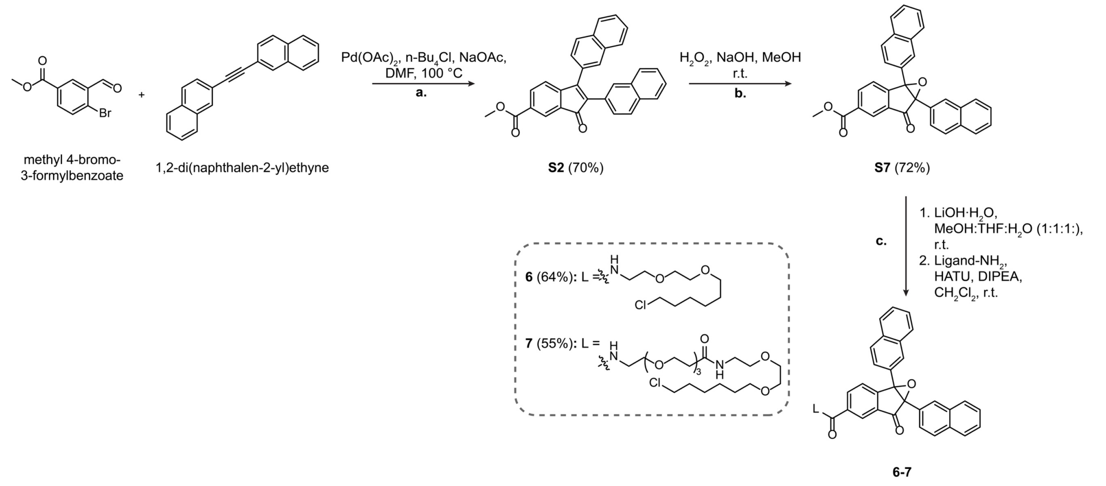

**Figure S4**: Synthesis of compounds **8, 9**. **a**. Palladium-catalyzed annulation of methyl 4-bromo-3-formylbenzoate and 1,2-bis(4/3-trifluoromethyl)phenyl)ethyne. **b.** Epoxidation. **c**. Amide coupling with HaloTag ligand.

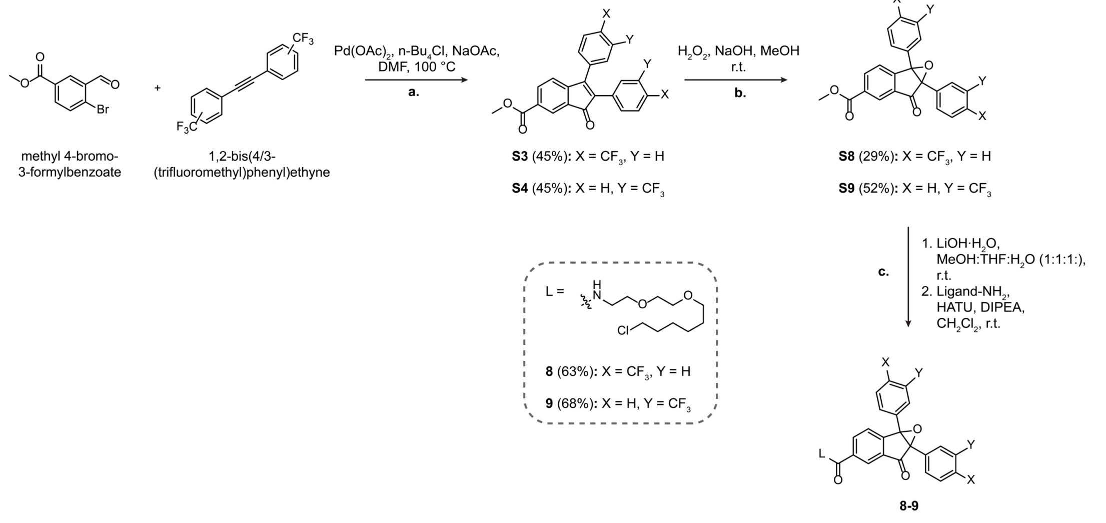

**Figure S5**: Synthesis of compound **10**. **a**. Palladium-catalyzed Suzuki coupling of 6-bromo-2,3-diphenyl-1H-inden-1-one and (4-(methoxycarbonyl)phenyl)-boronic acid. **b.** Epoxidation. **c**. Amide coupling with HaloTag ligand.

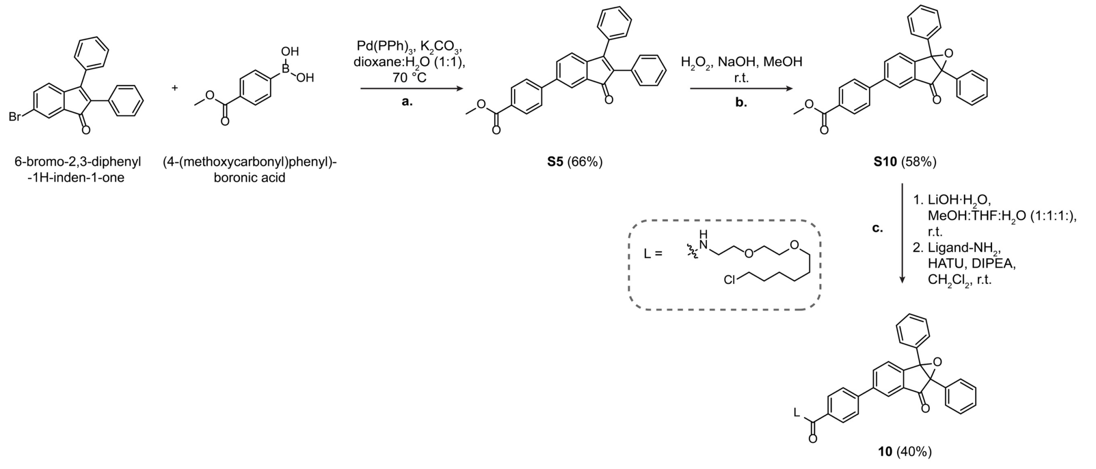

**Figure S6**: Synthesis of compound **S9**. **a**. Amide coupling followed by deprotection of the amino group.

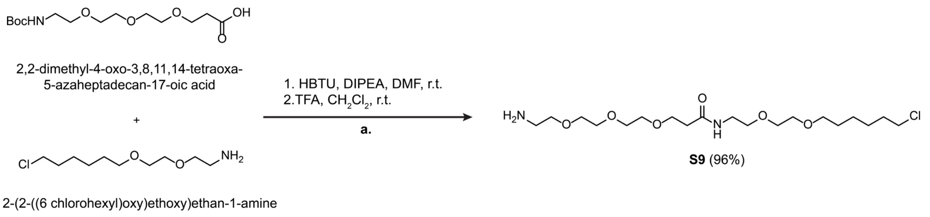

**Figure S7**: Picture of the custom-built illumination device used for inducing photoswitching *in vitro* and in live cell widefield imaging experiment. The illumination box was re-adapted from our previous work.[2] It was designed to fit a 96-well plate, and is composed of 24 LEDs and a controller to trigger illumination. Interchangeable LED plates at different wavelengths (375 nm, 450 nm and 545 nm) were assembled for this project.

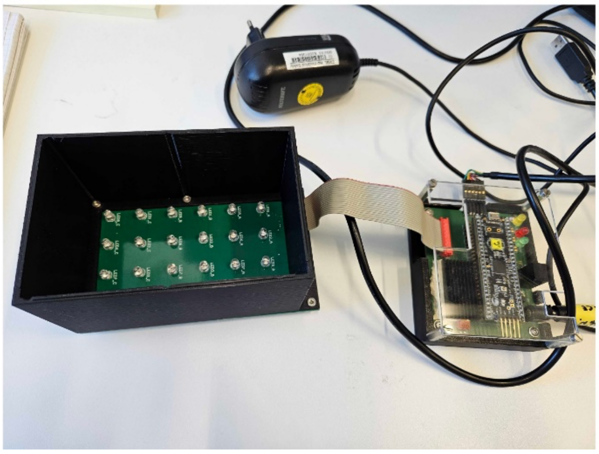

**Figure S8**: Absorption spectra of free ligands **1−10** in the dark (dashed black lines), at the photostationary state (PSS) after illumination at 375 nm (solid purple lines), and at the PSS after subsequent illumination at 545 nm (solid green lines) in MeCN/PBS (1:1). Absorption measurements were performed at 10 μM DIO-ligand. Mean of three replicates.

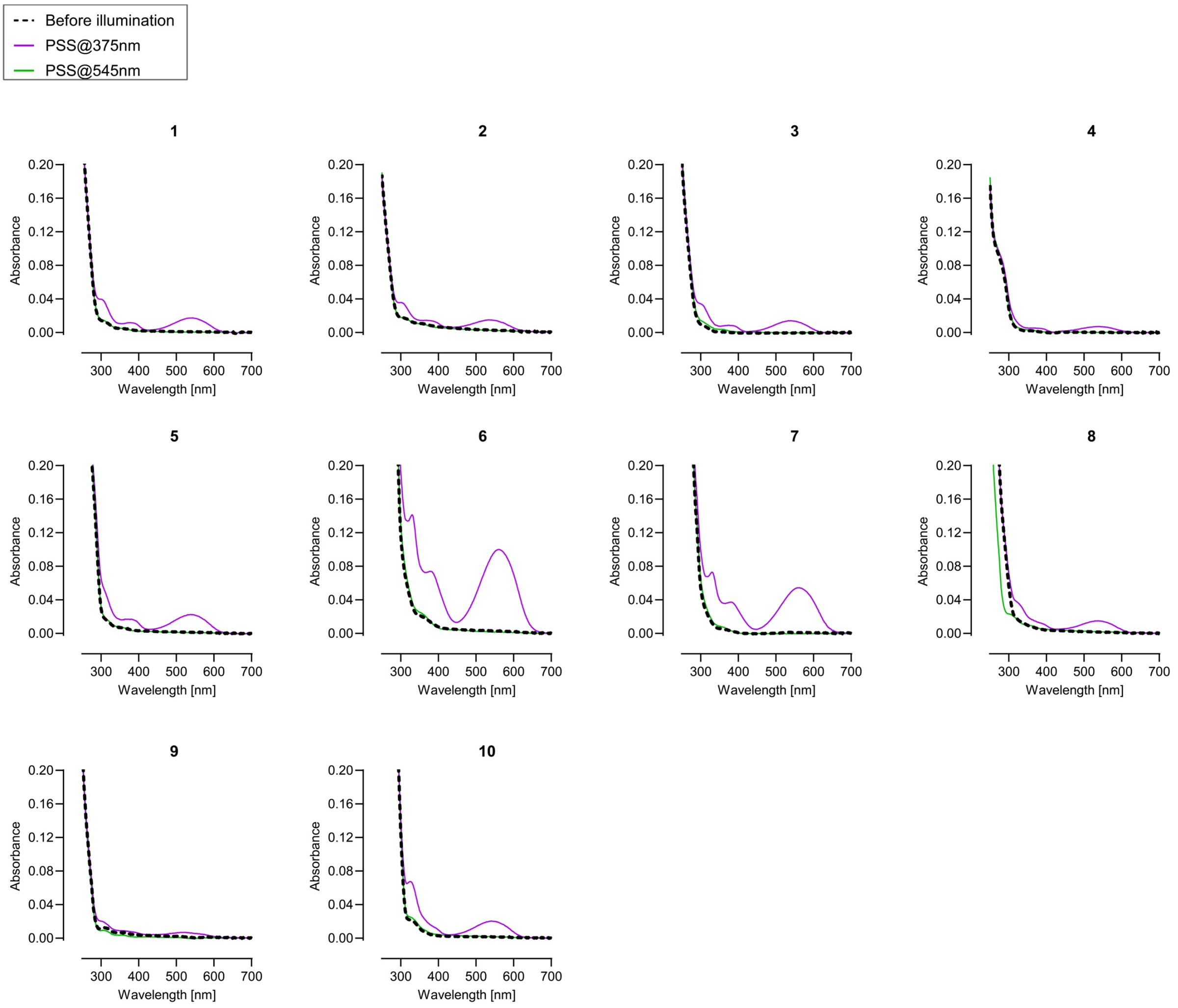

**Figure S9**: Absorption spectra during thermal recovery of free ligands **1−10** measured after illumination at 375 nm (left panel); corresponding linear fit of ln($A_0$/$A_t$) as a function of time, at $\lambda_{max}$ (middle panel); Absorption at $\lambda_{max}$ during successive cycles of illumination at 375 nm and 545 nm (right panel). Mean±SEM of three replicates.

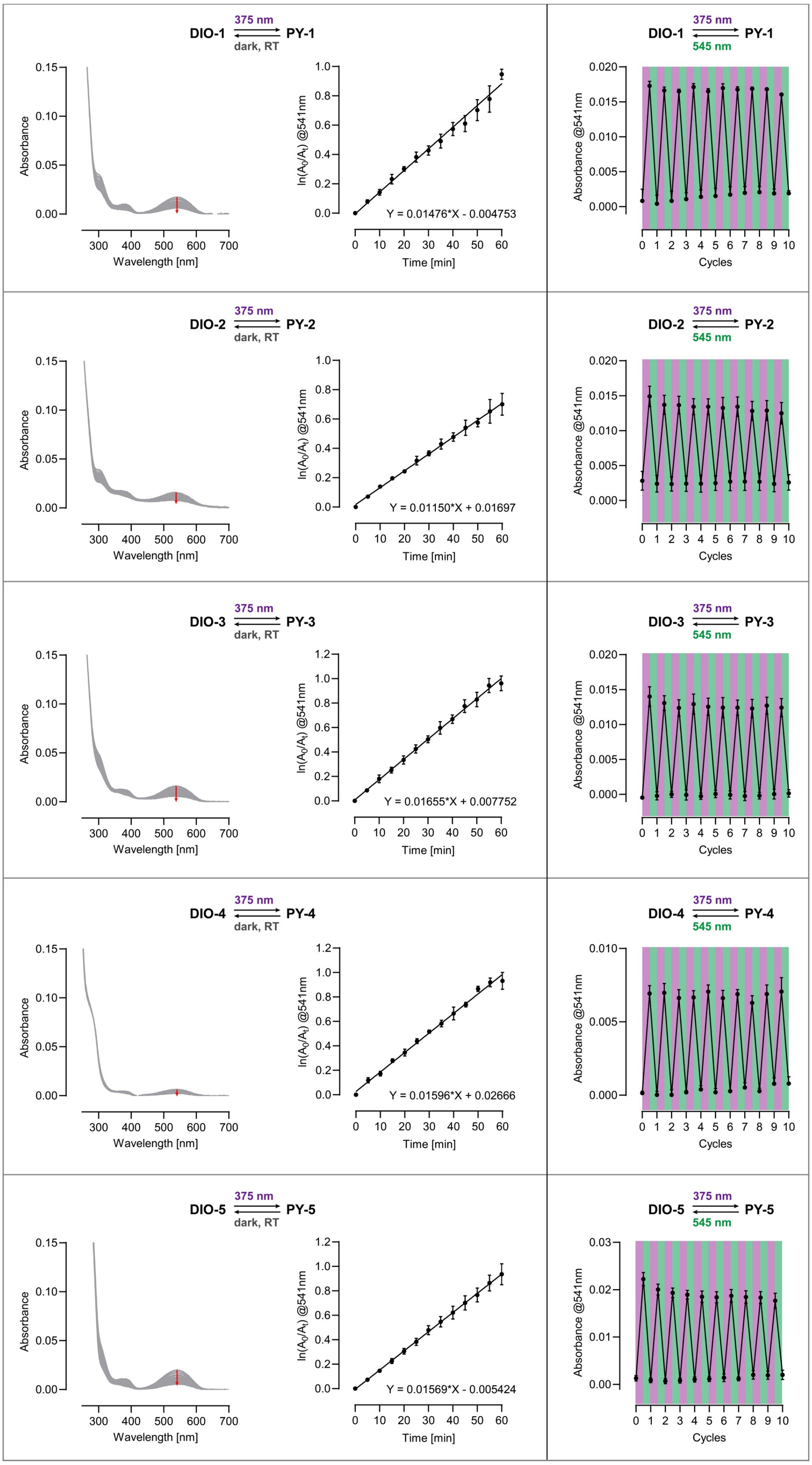

Continuation of **Figure S9**:

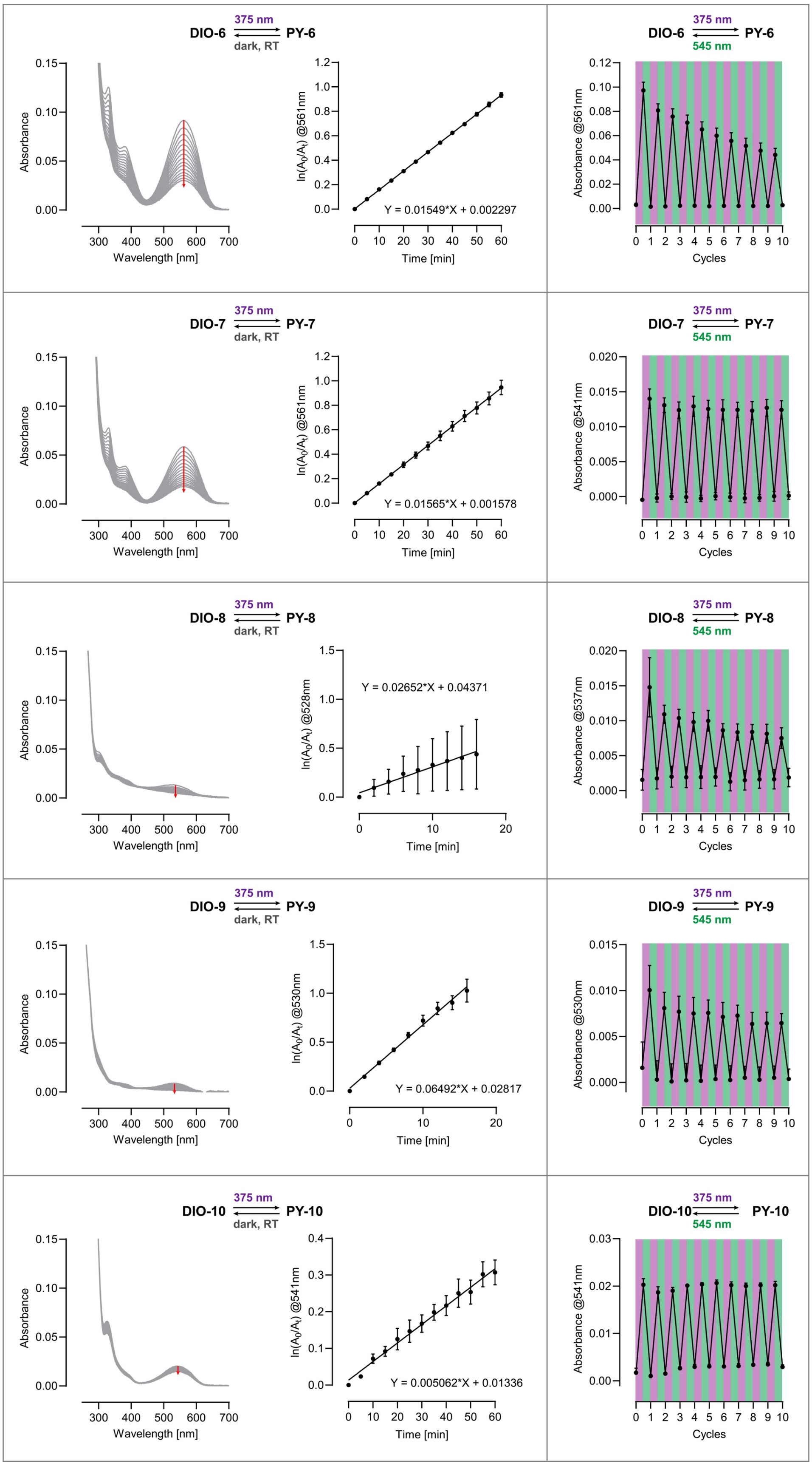

**Figure S10**: Absorption spectra of free ligands **1−10** in the dark (dashed black lines), at the PSS after illumination at 375 nm (solid purple lines), and at the PSS after subsequent illumination at 545 nm (solid green lines) in spectroscopy buffer (pH 7.4). Absorption measurements were performed at 10 µM DIO-ligand. Mean of three replicates.

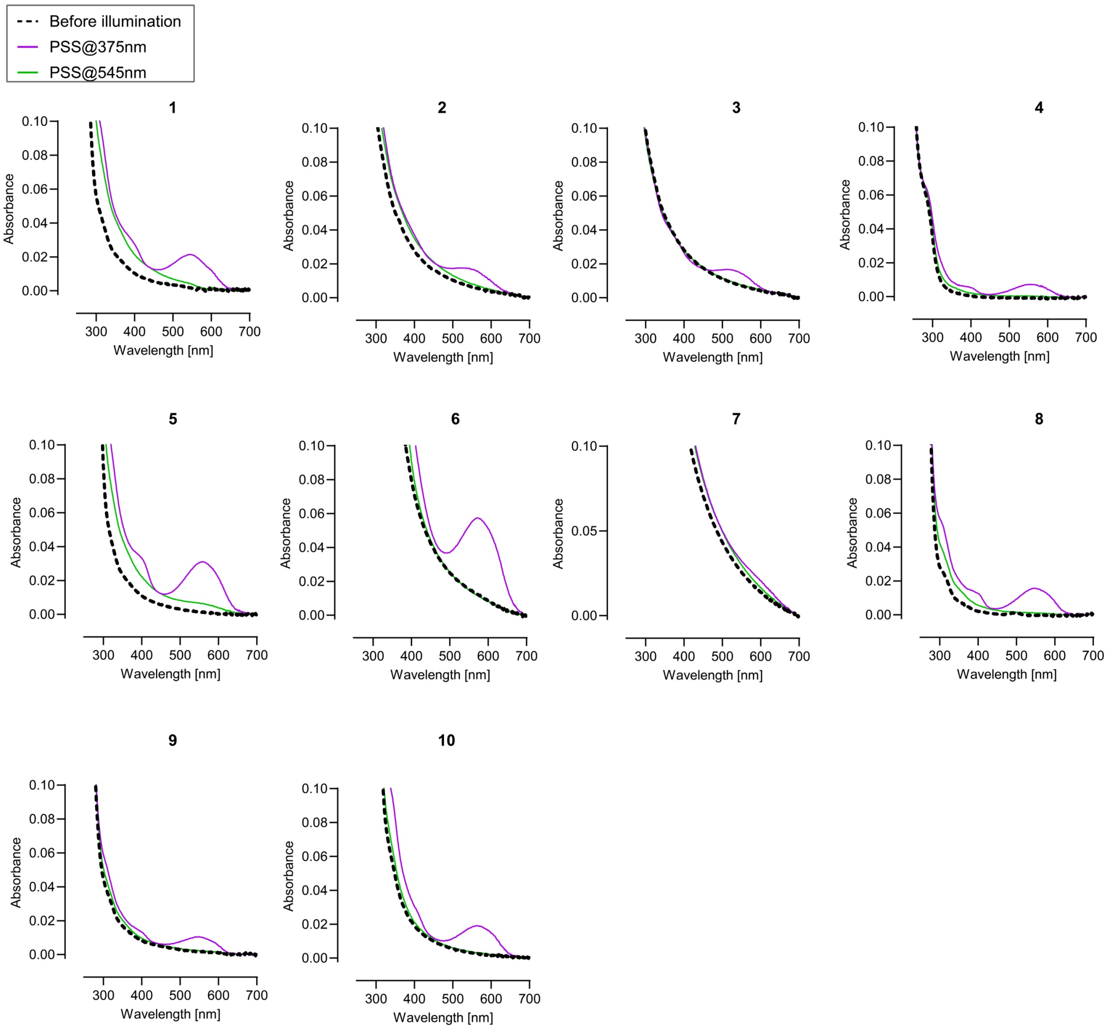

**Figure S11**: Measurement of the protein labeling efficiency of compounds **1−10** (2 μM) with the corresponding self-labeling protein (1 μM) *in vitro*, by pulse chase experiment with either $JF_{635}$-HaloTag ligand ($JF_{635}$-HTL, 1 μM) or $JF_{635}$-SNAP-tag ligand ($JF_{635}$-BG, 1 μM). **a, b.** Fluorescence intensity before (grey), and after addition of $JF_{635}$ ligand after 2 h (blue) and 5 h (purple), respectively, for HaloTag ligands (**a**.) and SNAP-tag ligands (**b.**). The controls were incubated for two hours for $JF_{635}$-HTL and for 5 h for $JF_{635}$-BG, respectively. Mean±SEM of two independent experiments, measured in triplicate.

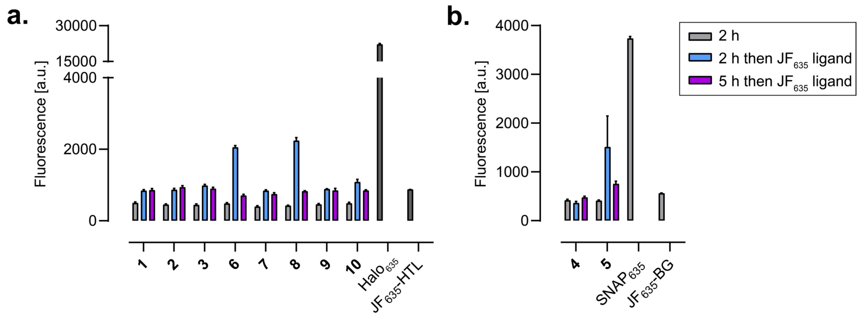

**Figure S12**: Absorption of ligands **1−10** bound to their respective protein tag (HaloTag for compounds **1−3** and **6−10**, SNAP-tag for compounds **4**,**5**) in the dark (dashed black lines), at the PSS after illumination at 375 nm (solid purple lines) and at the PSS after subsequent illumination at 450 nm (solid blue lines) in spectroscopy buffer (pH 7.4). Absorption measurements were performed at 5 μM DIO-ligand and 7.5 μM protein. Mean of two independent replicates. Compound **6** shows important aggregation which appears to impair photoswitching, likely due to poor binding to the protein.

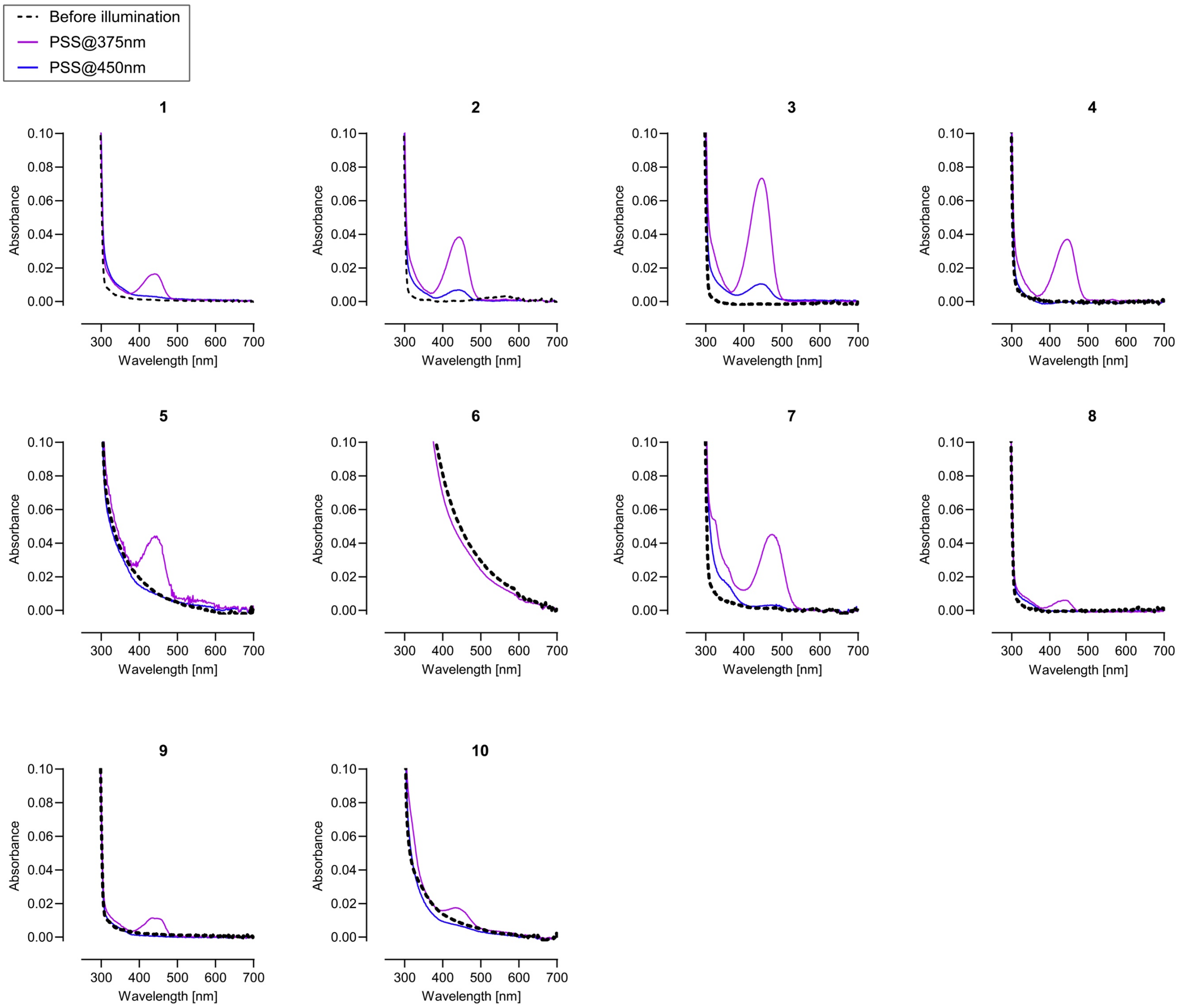

**Figure S13**: Normalized emission spectra of ligands **1−10** bound to their respective protein tag (HaloTag for compounds **1−3** and **6−10**, SNAP-tag for compounds **4** and **5**), at the photostationary state (PSS) after illumination at 375 nm in spectroscopy buffer (pH 7.4). Measurements were performed at 5 µM DIO-ligand and 7.5 µM protein. Mean of two replicates. Compound **6** was not measured due to aggregation.

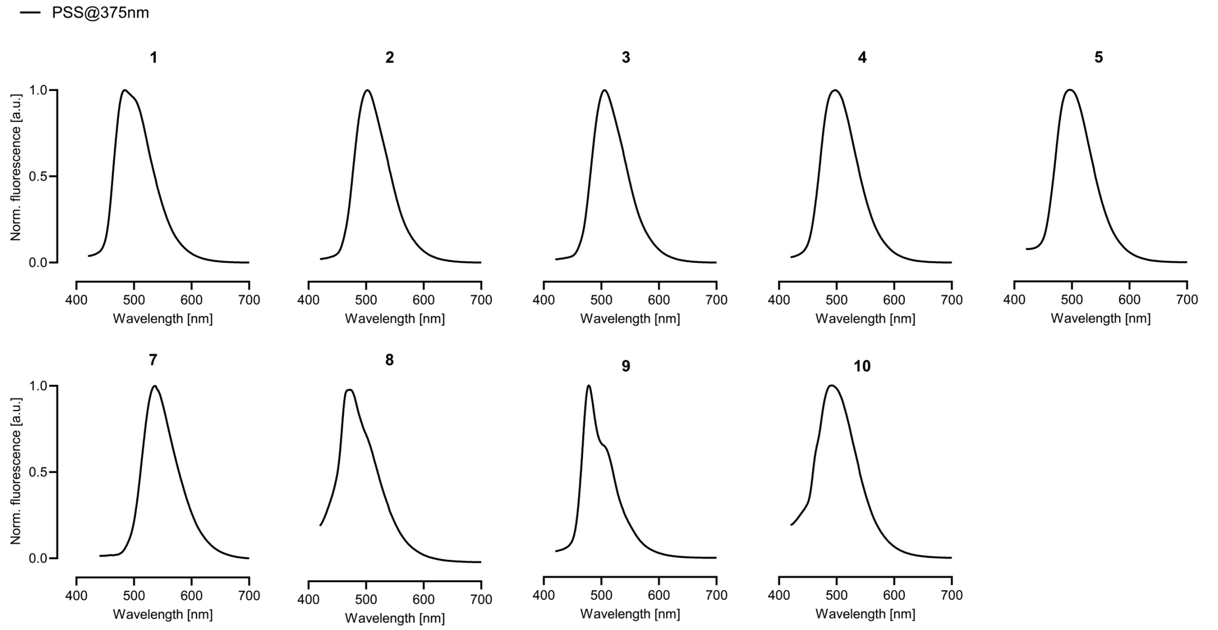

**Figure S14:** Thermal relaxation of selected HaloTag-bound ligand. Ligands **2**, **3**, **7** and **10** were bound to HaloTag and illuminated at 375 nm until the photostationary state was reached. Absorption at $\lambda_{max}$ was then monitored in the dark, at room temperature, over two hours. The HaloTag-bound ligands show no thermal relaxation over that time period. Mean of two replicates.

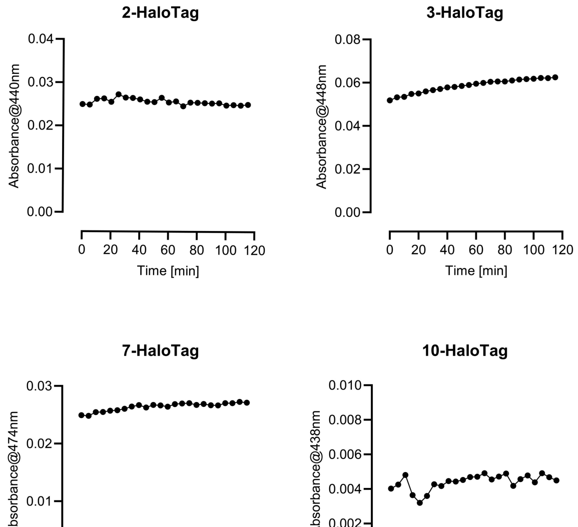

**Figure S15: a.** Electrostatic surface potential of HaloTag-TMR (PDB: 6U32)[3] and cp178HaloTag (Alphafold predicted structure), respectively. Electrostatic potentials are represented as protein surfaces from −2.0 (red) to 2.0 (blue) kJ·mol$^{-1}$ and were obtained using the APBS plugin in Pymol with standard parameters. The surface evidences differences in the orientation of residues near the ligand binding surface. **b.** Measurement of labeling efficiency of compound **7** (2 µM) with cp178HaloTag (1 µM) *in vitro*, by pulse chase assay with $JF_{635}$-HTL (1 µM). Fluorescence intensity before (grey) and after addition of $JF_{635}$-HTL after 2 h (blue). The control was incubated for two hours for $JF_{635}$-HTL. Mean±SEM of two independent experiment, measured in triplicate. The results show complete binding after two hours. **c.** Absorption spectrum of ligand **7** (5 µM) bound to cp178HaloTag (7.5 µM) in the dark (dashed black line), at the PSS after illumination at 375 nm (700 µW·cm$^{-2}$, solid purple line) and at the photostationary state (PSS) after subsequent illumination at 545 nm (700 µW·cm$^{-2}$, solid green line) in spectroscopy buffer (pH 7.4). Mean of two independent replicates.

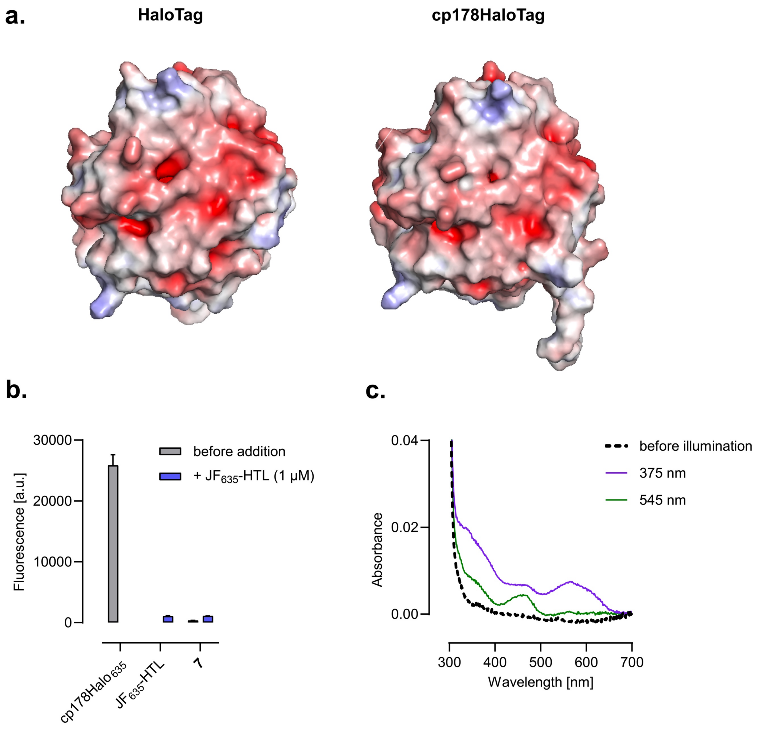

**Figure S16**: Absorption at $\lambda_{max}$ of protein-bound **1–10** under illumination at 375 nm over time in the absence (purple lines) and presence (black lines) of *endo*-BCN. Two samples of ligands **1–10** (5 µM) were incubated with the respective protein tag (HaloTag for compounds **1–3** and **6–10**, SNAP-tag for compounds **4**,**5**, 7.5 µM), for two hours in the dark. To one sample, *endo*-BCN (10 µM) was added before illumination, and both samples were illuminated at 375 nm (700 µW·cm$^{-2}$) while absorbance was monitored at $\lambda_{max}$. Mean of two independent replicates.

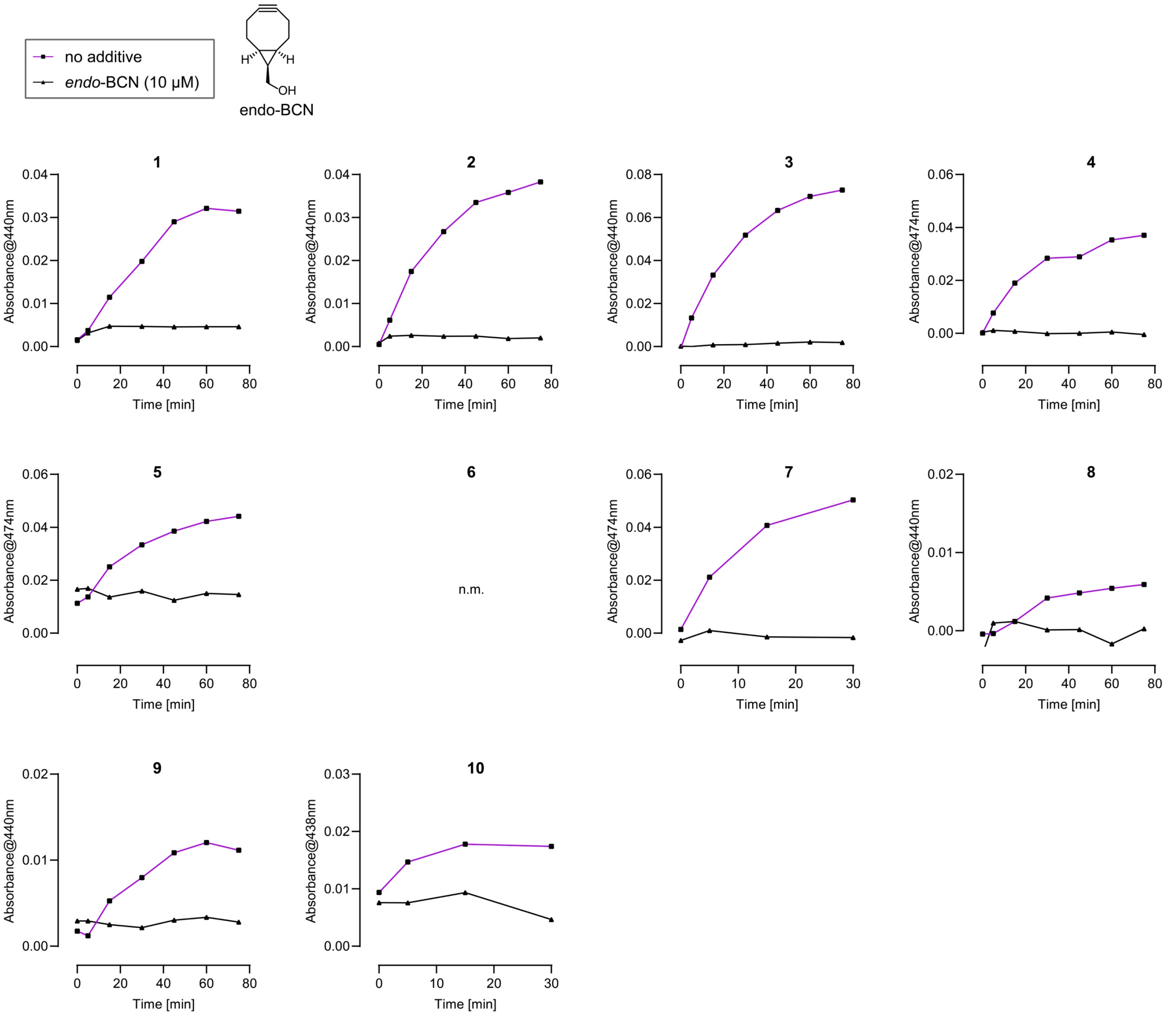

**Figure S17**: Investigation of the photoclick reaction between *endo*-BCN and ligand **7**. **a,b**. Ligand **7** (5 µM) was incubated with HaloTag (7.5 µM) for 2 h in the dark. *endo*-BCN was added either before illumination (**a**.) or after illumination, once the PSS had been reached (**b**.). Illumination time is indicted by the purple area. Mean of two independent replicates. **c,d.** Same experiment as in a,b. for the free ligand **7**.

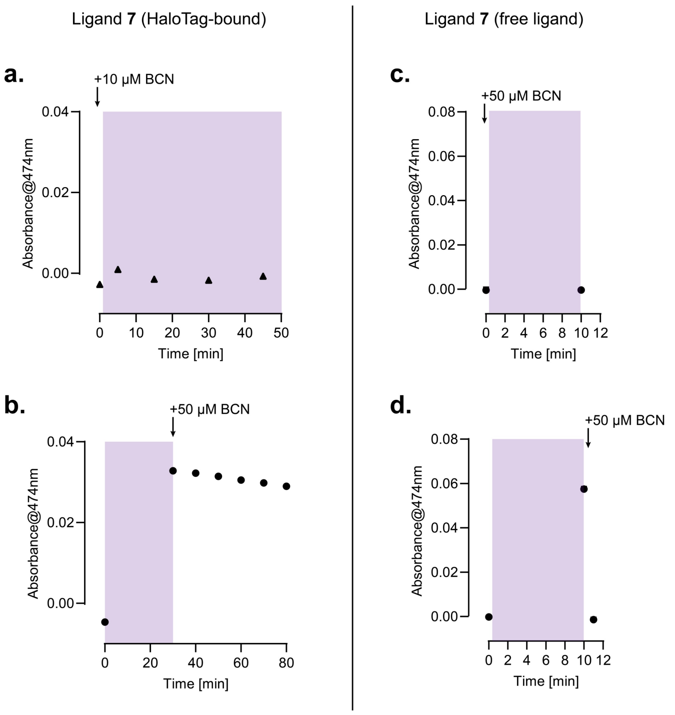

**Figure S18**: Synthesis of compound **11** by amide coupling.

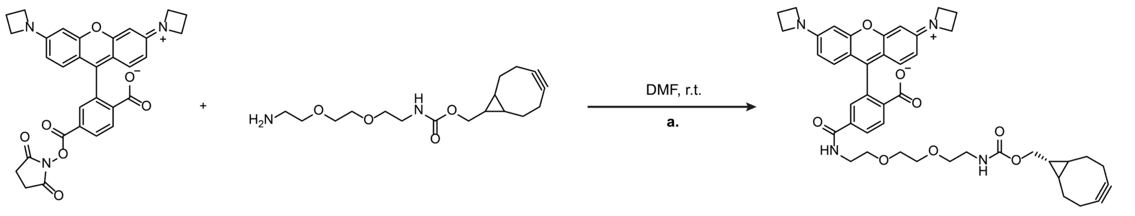

JF549-NHS ester BCN-amine **11** (63%)

**Figure S19**: Normalized absorption and emission spectra of **11** in spectroscopy buffer (pH 7.4). Mean of two replicates.

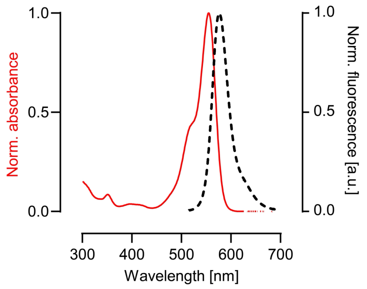

**Figure S20**: Verification of the click reactivity of JF$_{549}$-BCN (**11**). **a**. Reaction scheme of the click reaction between JF$_{549}$-BCN (**11**) and 3,6-di-2-pyridyl-1,2,4,5-tetrazine. **b**. LCMS traces and corresponding detected mass of JF$_{549}$-BCN (**11**) (top panel), 3,6-di-2-pyridyl-1,2,4,5-tetrazine (middle panel) and the cycloadduct (bottom panel).

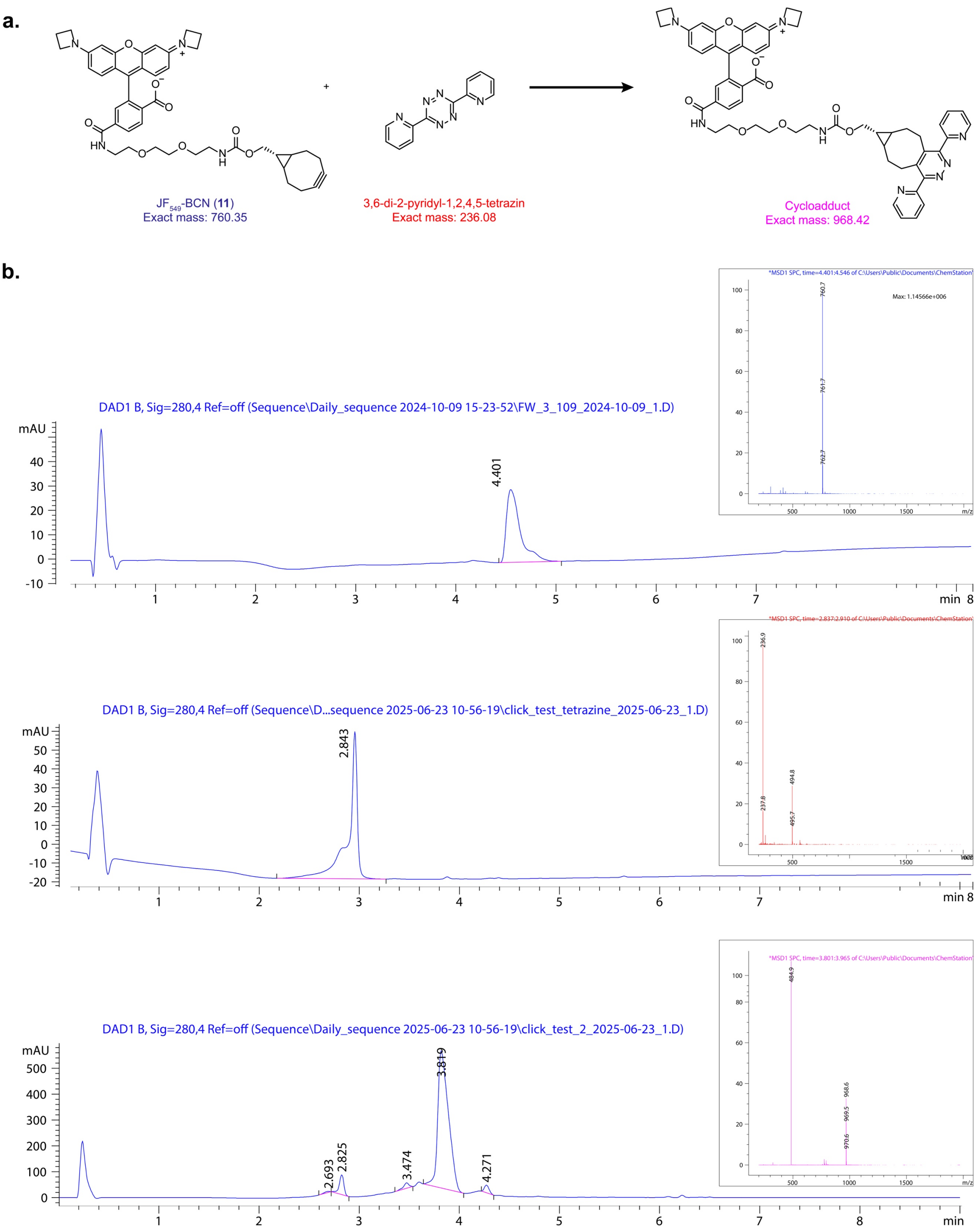

**Figure S21**: Characterization of the click reaction between ligand **3** bound to HaloTag and $JF_{549}$-BCN (**11**) by SDS-PAGE gels. As a control, HaloTag labeled with $JF_{549}$-HaloTag ligand ($Halo_{549}$) was used to determine the maximum fluorescence labeling in each experiment. **a.** Fluorescence (top panel) and Coomassie staining (bottom panel) of an SDS-PAGE gel, with varying concentration of **11**. **b.** Dose-response curves from fluorescence intensity analysis of gels corresponding to panel a. **c.** Fluorescence (top panel) and Coomassie staining (bottom panel) of an SDS-PAGE gel, with varying illumination time at 375 nm. **d.** Dose-response curves from fluorescence intensity analysis of gels corresponding to panel c. **e.** Fluorescence (top panel) and Coomassie staining (bottom panel) of an SDS-PAGE gel, with varying illumination intensity. **f.** Dose-response curves from fluorescence intensity analysis of gels from the click reaction corresponding to panel e. Mean±SEM of three replicates. L: protein ladder, X: no ligand.

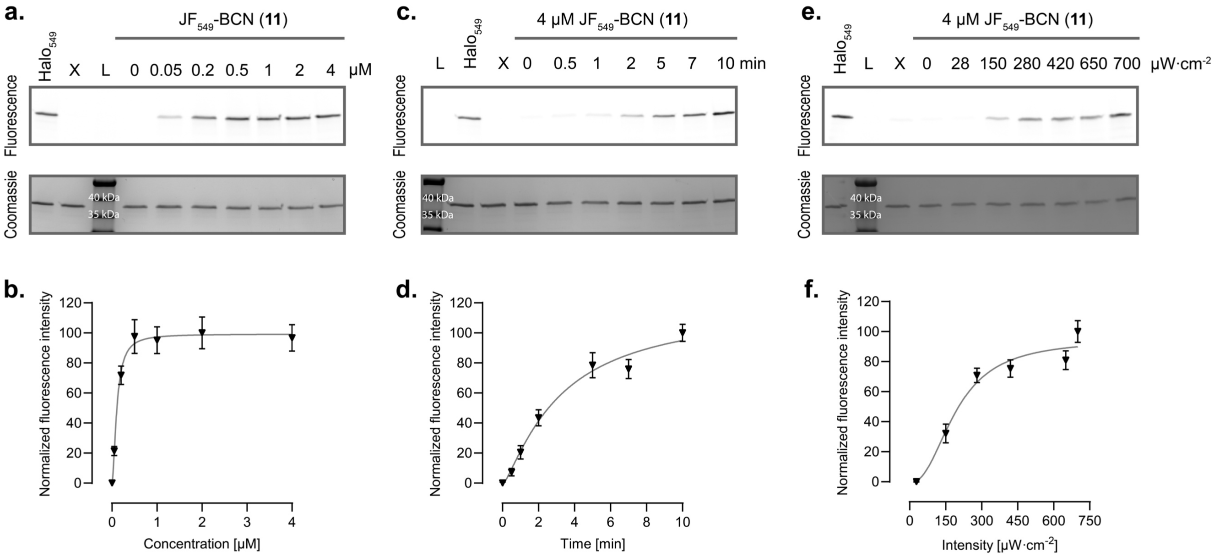

**Figure S22:** Measurement of labeling efficiency of compounds **1−10** in living U2OS cells, by pulse chase experiment with $JF_{635}$-HaloTag ligand (1 µM) or $JF_{635}$-SNAP-tag ligand (1 µM): extracellularly on the cell surface (**a**, **b**.) intracellularly in the cytosol (**c**, **d**.). U2OS cells co-expressing EGFP in the cytosol and SNAP-tag−HaloTag either on the cell surface or in the cytosol, were incubated with ligands **1−10** (2 µM), for 2 h at 37°C and washed 3 times with prewarmed imaging buffer. $JF_{635}$-HaloTag ligand (1 µM) or $JF_{635}$-SNAP-tag ligand (1 µM) was then added and the cells were incubated for another 30 min at 37°C. After washing with prewarmed imaging buffer the cells were imaged. The EGFP signal was cytosolic in both experiments. For quantification, a code based on Cellpose was used creating a mask based on the EGFP signal (see Methods). The normalized fluorescence ratio of the $JF_{635}$ and EGFP fluorescence signal was plotted for individual cells. Mean±CI 95% for N > 200 cells in two independent replicates for each condition. **a.** HaloTag ligands **1−3** and **6−10** on the cell surface. Statistical test: unpaired t-test vs the control ($Halo_{635}$) value $p < 0.0001$ (****). **b.** SNAP-tag ligands **4** and **5** on the cell surface. Statistical test: unpaired t-test vs the control ($SNAP_{635}$) value ns ($p > 0.05$), indicating no binding. **c.** HaloTag ligands **1−3** and **6−10** inside living cells. Statistical test: unpaired t-test vs the control ($Halo_{635}$) value **** $p<0.0001$. **b.** SNAP-tag ligands **4** and **5** inside cells. Statistical test: unpaired t-test vs the control ($SNAP_{635}$) value ns ($p > 0.05$), indicating that ligands **4** and **5** do not bind efficiently. X: no ligand.

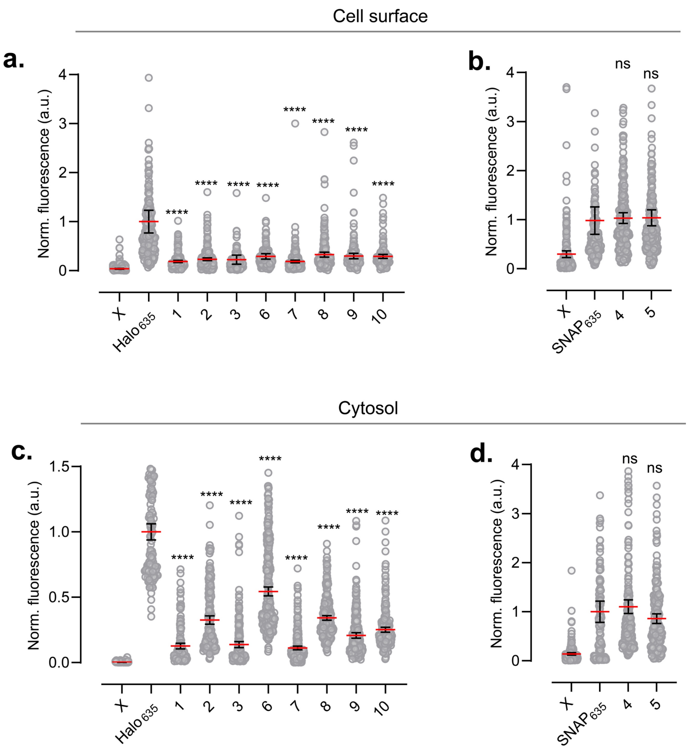

**Figure S23:** Representative widefield fluorescence images of U2OS cells co-expressing a SNAP-tag−HaloTag fusion on the cell surface and EGFP in the cytosol, incubated with ligands **1−10** (1 µM) for 2 h at 37°C in the dark**,** and after photoclick reaction with CF®647-TCO (2 µM) induced by illumination at 375 nm for 5 minutes (700 µW·cm$^{-2}$). From left to right: fluorescence in the EGFP channel; fluorescence of CF®647 in the far-red channel; overlay of the EGFP and far-red channels. Representative images from two independent experiments, two wells for each condition, nine fields of view per well. Scale bars: 50 µm.

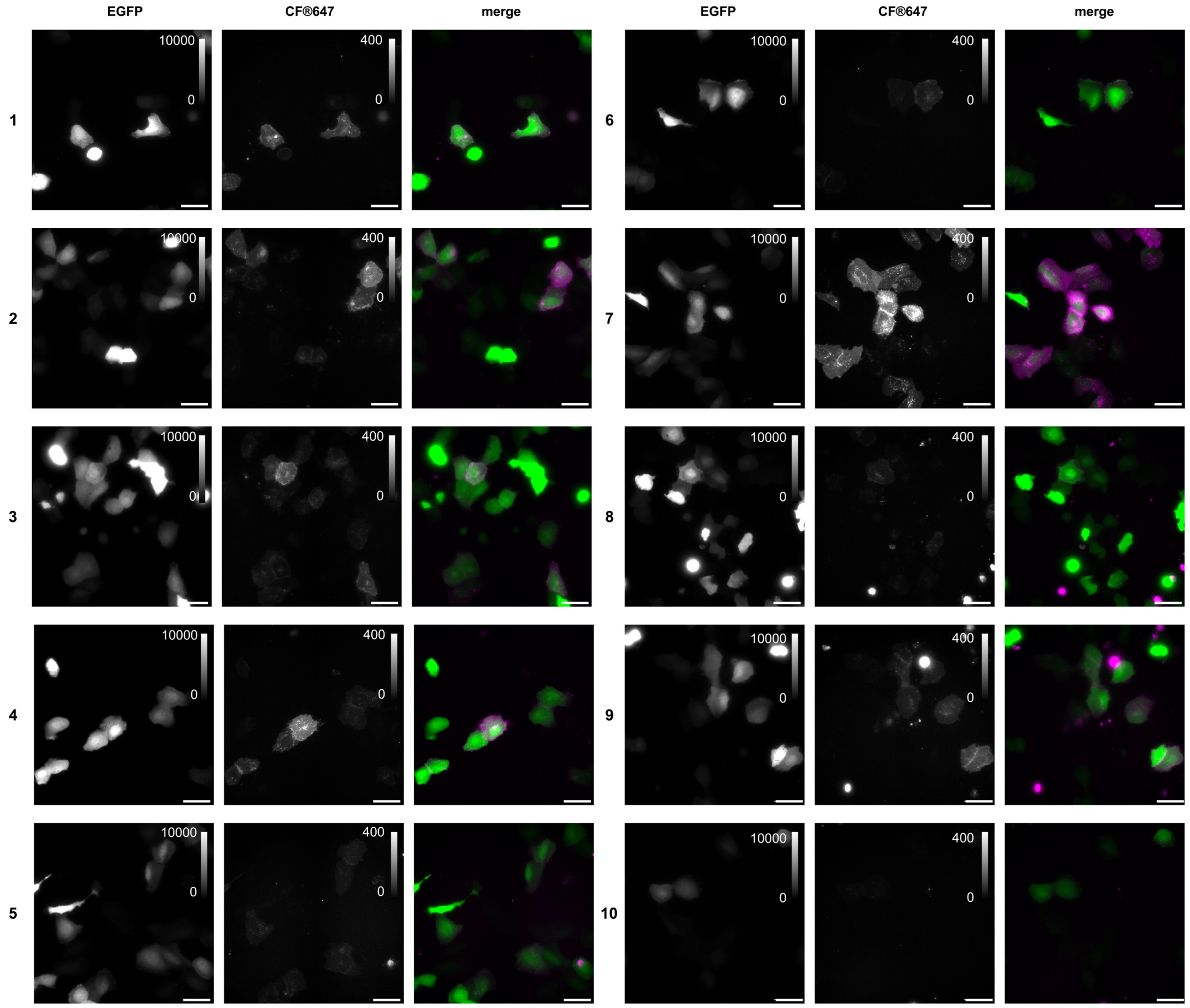

**Figure S24**: Representative widefield fluorescence images of U2OS cells co-expressing a SNAP-tag−HaloTag fusion on the cell surface and EGFP in the cytosol, either without ligand or labeled with ligand **7** (1 μM) for 2 h at 37°C in the dark, after incubation with CF®647-TCO (2 μM) and for the illuminated negative sample (middle) illumination at 375 nm for 5 minutes (700 μW·cm$^{-2}$). From left to right: fluorescence in the EGFP channel; fluorescence of CF®647 in the far-red channel; overlay of the EGFP and far-red channels. Representative images from two independent experiments, two wells for each condition, nine fields of view per well. Scale bars: 50 μm.

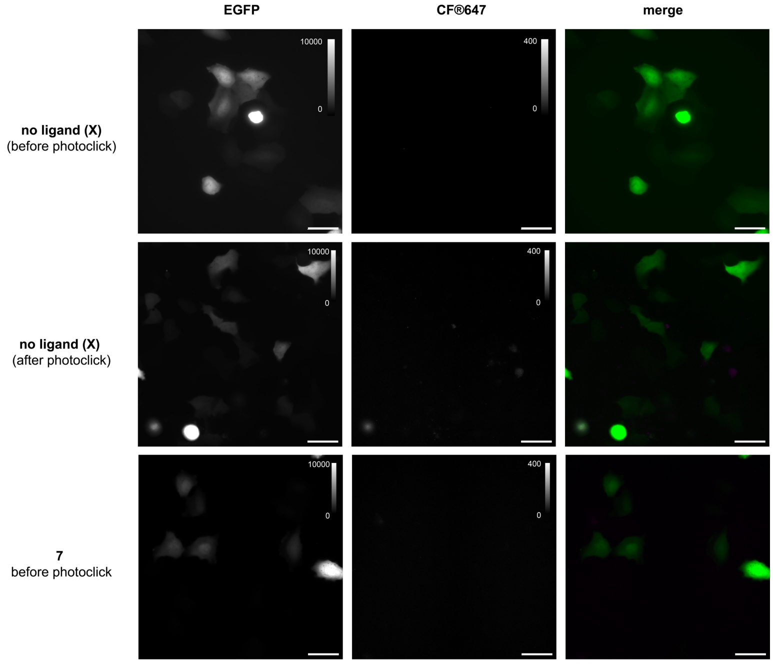

**Figure S25**: Normalized fluorescence ratio of the CF®647/EGFP channels for U2OS cells co-expressing a SNAP-tag−HaloTag fusion on the cell surface and EGFP in the cytosol, and labelled with either ligands **1−10** or no ligand (X) in the dark (**a**.) and after photoclick reaction (**b**.). Mean±CI 95% for N > 200-300 cells in two independent experiments for each condition. Statistical test: unpaired t-test vs the control (no ligand: X) values: ****p < 0.0001, *** p = 0.0006 and ** p = 0.0031. In **a.** all values are statistically insignificant vs the control (p > 0.05).

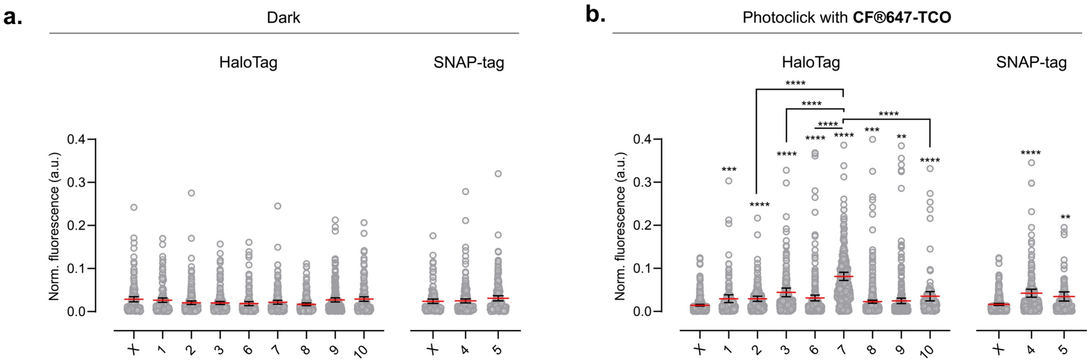

**Figure S26**: Characterization of photoclick reaction on the surface of living cells by flow cytometry. U2OS cells co-expressing a SNAP-tag−HaloTag fusion on the cell surface and EGFP in the cytosol were incubated with ligands **1−10** (1 µM), for 2 h at 37°C in the dark, and washed four times with prewarmed imaging buffer. CF®647-TCO (2 µM) was added to the well and incubated for 5 min. The plates were illuminated at 375 nm (700 µW·cm$^{-2}$, 5 min) followed by washing with prewarmed imaging buffer five times. Ice-cold PBS and cell scrapers were used to detach the cells. As controls to define the gates, the following samples were used: to check for transfection efficiency, transfected cells were labeled with $JF_{635}$-HaloTag ligand (1 µM), unlabeled transfected cells served as negative control for the far-red channel and non-transfected cells served as negative control for the EGFP gating. 10000 EGFP-positive cells were detected and their CF®647 fluorescence signal was measured. **a**. Overlaid histograms of the CF®647 fluorescence for EGFP-positive cells from each ligand **1−10**. It shows which cells expressed correctly, and can be detected in the CF®647 channel. **b**. Overlaid histograms of the CF®647 fluorescence for CF®647-positive cells from each ligand **1−10**. **c**. Plot of the measured mean fluorescence intensity after photoclick reaction with CF®647-TCO for the different ligands. a.b. correspond to one representative example of two independent experiments displayed in c.

**a.**

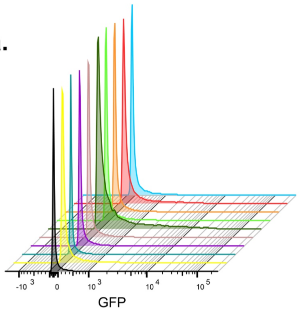

**b.**

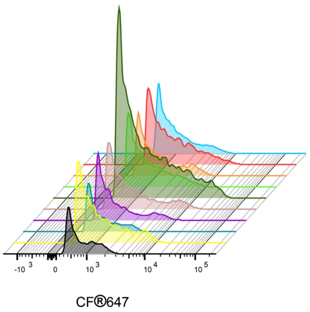

| Sample Name | Subset Name | Count | Mean : 640-670_30-A |
|---|---|---|---|
| Ligand **10** | eGFP + | 29283 | 80,1 |
| Ligand **9** | eGFP + | 26073 | 312 |
| Ligand **8** | eGFP + | 26004 | 94,3 |
| Ligand **5** | eGFP + | 26260 | 176 |
| Ligand **4** | eGFP + | 27364 | 283 |
| Ligand **7** | eGFP + | 30491 | 1137 |
| Ligand **6** | eGFP + | 28405 | 216 |
| Ligand **3** | eGFP + | 24502 | 203 |
| Ligand **2** | eGFP + | 20954 | 416 |
| Ligand **1** | eGFP + | 23572 | 195 |

| Sample Name | Subset Name | Count | Mean : 640-670_30-A |
|---|---|---|---|
| Ligand **10** | TCO-647 | 1352 | 919 |
| Ligand **9** | TCO-647 | 3707 | 1773 |
| Ligand **8** | TCO-647 | 1445 | 1000 |
| Ligand **5** | TCO-647 | 2234 | 1483 |
| Ligand **4** | TCO-647 | 3036 | 2056 |
| Ligand **7** | TCO-647 | 9329 | 3424 |
| Ligand **6** | TCO-647 | 3120 | 1454 |
| Ligand **3** | TCO-647 | 2472 | 1458 |
| Ligand **2** | TCO-647 | 3814 | 1890 |
| Ligand **1** | TCO-647 | 2517 | 1289 |

**c.**

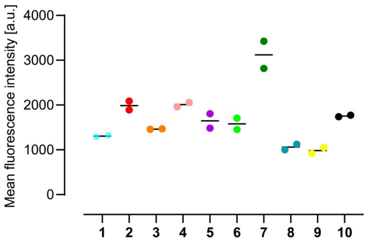

**Figure S27**: Flow cytometry gating strategy for analysis of the photoclick reaction of ligand **7** bound to the HaloTag-protein on the surface of EGFP-positive cells. Cells were first gated based on forward and side scatter to select the main population (top left panel), followed by exclusion of doublets using FSC-H versus FSC-A properties (top right panel). Within the singlet population, 10000 EGFP-positive cells were identified and selected for analysis (middle left panel). The percentage of EGFP-positive cells is reported. CF®647 fluorescence was then analyzed within this EGFP-positive population by both histogram (middle right panel) and two-dimensional gating (middle left panel), allowing quantification of the fraction of EGFP-positive cells exhibiting CF®647-TCO signal. Percentages at each gate reflect the proportion of events passing to the next step, ensuring robust analysis of CF®647 signal specifically within single, EGFP-positive cells.

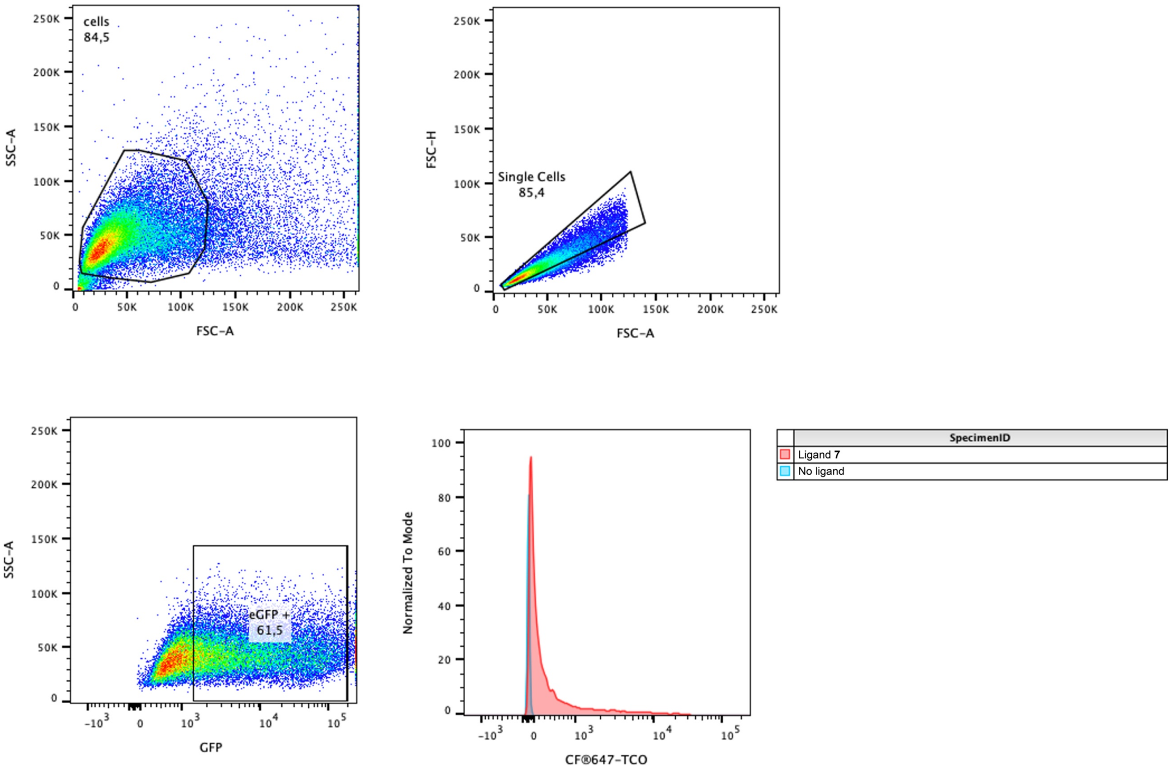

**Figure S28**: MTS assay to quantify cell viability of transfected U2OS cells expressing a SNAP-tag−HaloTag fusion on the cell surface, labeled with compound **1**, **7**, **10** (1 µM) or no ligand (X), in the dark and after illumination (375 nm, 700 µW·cm$^{-2}$, 5 min). Mean±SEM of two independent experiments performed in triplicates, where unpaired t-test showed no significant p values ($p > 0.005$ in all cases). Values are normalized to the dark, transfected and unlabeled sample (X).

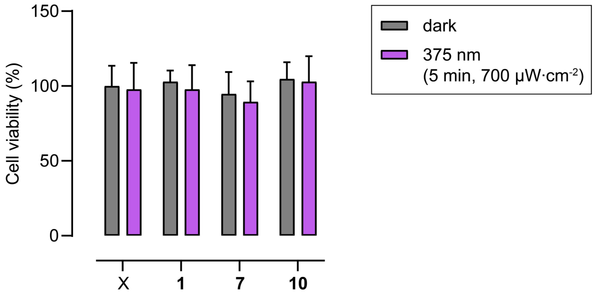

**Figure S29**: Additional representative example of spatiotemporal two-color labeling of cell surface regions of living U2OS cells. U2OS cells expressing a SNAP-tag−HaloTag fusion on the cell surface and EGFP intracellularly were incubated with ligand **7** (1 μM) for 2 h at 37 °C in the dark (T0). Sequential ROI labeling (T1–T3) was performed using either CF®647-TCO (2 μM) or Cy3-TCO (2 μM) (see Methods). Photolabeling was induced by direct illumination from the microscope at 405 nm (4μs/pixel, 0.5% power level (41 kW·cm$^{-2}$)) for 20 s, during which each ROI was scanned several times. Fluorescence of Cy3 (top row) and CF®647 (middle row) channels after activation of different ROIs; Bottom row: fluorescence in EGFP channel with selected ROIs for the 3 sequential rounds (left panel); overlay of the Cy3 and far-red channels after the final photoclick round (T3) (right panel). Representative images from five independent experiments. Scale bars: 50 μm.

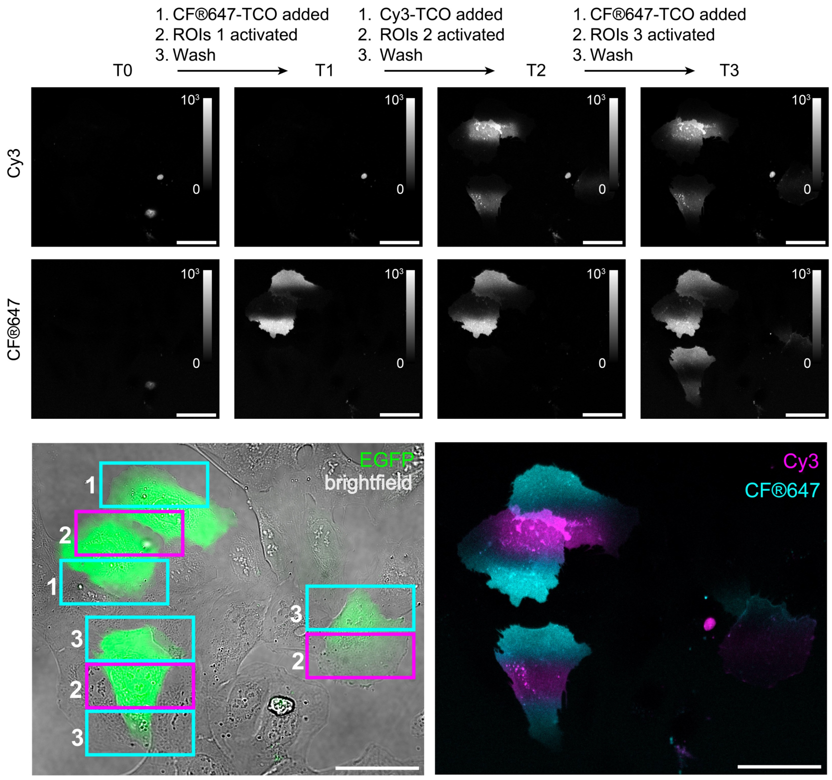

**Figure S30:** Representative confocal images of a U2OS cell expressing a SNAP-tag−HaloTag fusion on the cell surface and EGFP intracellularly, labelled with ligand **7**, after two-color photoclick on the cell surface with CF®647-TCO and Cy3-TCO. Image immediately after photoclick, and image after 30 minutes of imaging. The full timelapse can be found in Movie S1. Scale bars: 20 µm.

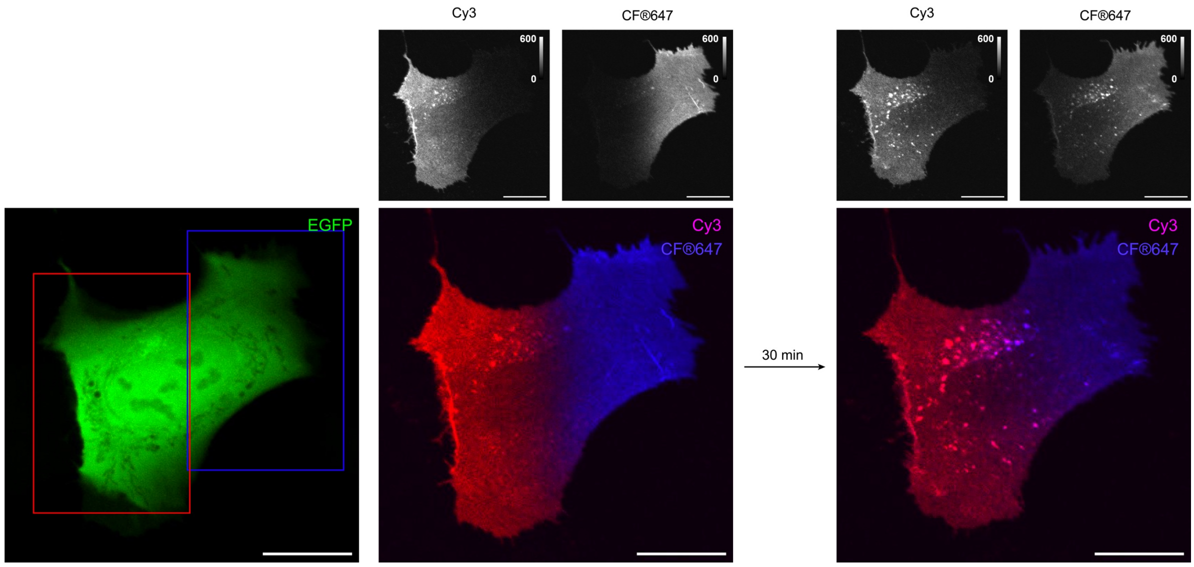

**Figure S31**: Additional representative example of spatiotemporal three-color labeling of cell surface regions of living U2OS cells. U2OS cells expressing a SNAP-tag−HaloTag fusion on the cell surface were incubated with a mixture of $JF_{722}$-HaloTag ligand (100 nM) and ligand **7** (1 µM) for 2 h at 37 °C, in the dark. $JF_{722}$ was used as marker to identify transfected cells. Sequential ROI photolabeling was performed using CF®488A-TCO (2 µM), Cy3-TCO (2 µM) and CF®647-TCO (2 µM). Photolabeling was induced by direct illumination from the microscope at 405 nm (4µs/pixel, 0.5% power level (41 $kW \cdot cm^{-2}$)) for 20 s, during which each ROI was scanned several times. **a.** Representative fluorescence image before photobarcording. $JF_{722}$-HaloTag ligand signal was very faint and did not interfere with the signal from the photoclicked dyes, while still allowing for identification of HaloTag-expressing cell. **b.** Representative fluorescence image of the same cell after sequential spatially defined labeling with CF®488A-TCO (green), Cy3-TCO (magenta) and CF®647-TCO (blue) in different regions. Scale bars: 10 µm.

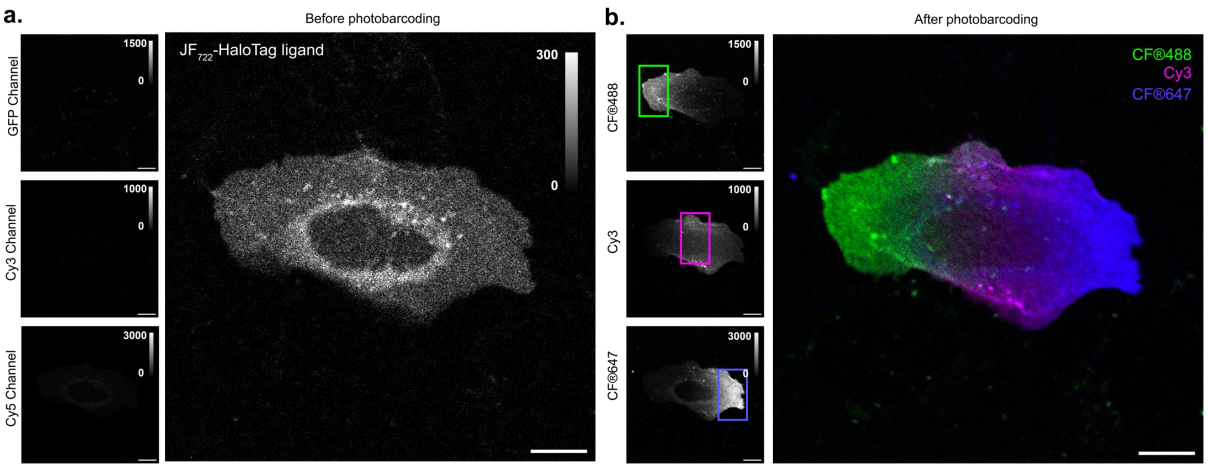

**Figure S32**: Proof-of-principle of fluorescence barcoding of living cells using controlled combinatorial labeling with two fluorophores. U2OS cells expressing a SNAP-tag−HaloTag fusion on the cell surface and EGFP in the cytosol were incubated with ligand **7** (1 µM) for 2 h at 37 °C in the dark (T0). Sequential photolabeling of single cells with varying illumination times was performed using CF®647-TCO (2 µM) and then Cy3-TCO (2 µM). Photolabeling was induced by direct illumination from the microscope at 405 nm (4µs/pixel, 0.5% power level (41 kW·cm$^{-2}$)) for different durations. **a.** From left to right: EGFP fluorescence indicating selected cells and corresponding illumination times (CF®647-TCO/Cy3-TCO in seconds, annotated next to the corresponding cell); Cy3 (top) and CF®647 (bottom) fluorescence following activation of distinct cells with different illumination durations; merged Cy3 and far-red channels after the second photoclick step (T2). Representative images from three independent experiments. Scale bars: 50 µm. **b.** Illumination times and signal intensity in each channel after the two-step photoclick labeling. **c.** Correlation between the ratio of the illumination time and the ratio of signal intensity in the CF®647 and Cy3 channels after the two-step photoclick labeling.

**a.**

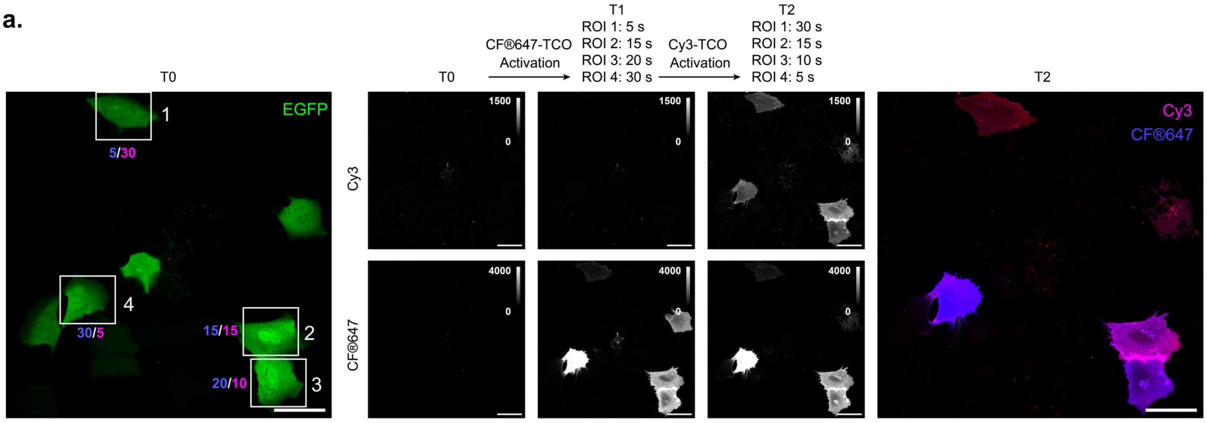

**b.**

| Cell | Illumination time CF®647/Cy3 [s] | Signal intensity (mean) EGFP | CF®647 | Cy3 | Ratio CF®647/Cy3 |
|---|---|---|---|---|---|
| 1 | **5/30** | 1537 | 351 | 411 | **0.9** |
| 2 | **15/15** | 2326 | 2403 | 786 | **3.1** |
| 3 | **20/10** | 2426 | 2501 | 526 | **4.8** |
| 4 | **30/5** | 1836 | 4081 | 519 | **7.9** |

**c.**

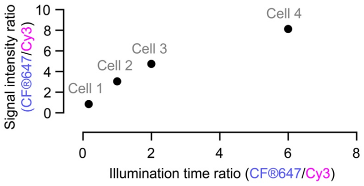

### Supplementary Tables

**Table S1**: Spectroscopic properties of ligands **1−10** as free ligands in solution (PBS:MeCN 1:1), and protein-bound. Wavelengths ($\lambda_{max}$, $\lambda_{em}$) and apparent extinction coefficients ($\varepsilon_{PSS}$) correspond to the photostationary state at 375 nm. $t_{1/2, relax}$ corresponds to the half-life of the thermal relaxation (measured in the dark at room temperature). n.m.: not measured.

| Free ligands (PBS:MeCN 1:1) | | | | Protein bound ligands | | |
|---|---|---|---|---|---|---|
| **Compound** | **$\lambda_{max}$ [nm]** | **$\varepsilon_{PSS}$ [$M^{-1}\cdot cm^{-1}$]** | **$t_{1/2, relax}$ [min]** | **$\lambda_{max}/\lambda_{em}$ [nm]** | **$\varepsilon_{PSS}$ [$M^{-1}\cdot cm^{-1}$]** | **$\Phi_F$** |
| **1** | 541 | 1700 | 68 | 440/483 | 3300 | n.m. |
| **2** | 541 | 600 | 87 | 440/502 | 5000 | 0.11 |
| **3** | 541 | 1400 | 60 | 448/505 | 1400 | 0.10 |
| **4** | 541 | 700 | 63 | 445/497 | 4900 | 0.12 |
| **5** | 541 | 2200 | 64 | 440/496 | 5900 | n.m. |
| **6** | 561 | 10000 | 65 | n.m. | n.m. | n.m. |
| **7** | 561 | 5400 | 64 | 474/537 | 6000 | 0.24 |
| **8** | 537 | 1500 | 38 | 437/474 | 800 | n.m. |
| **9** | 530 | 600 | 15 | 448/478 | 1500 | n.m. |
| **10** | 541 | 2000 | 198 | 438/491 | 2300 | <1% |

**Table S2**: Illumination times required to reach respective photostationary states (PSS) for ligands **1−10** under our illumination conditions (700 μW·$cm^{-2}$). Measurements were performed at 10 μM for the free ligands and 5 μM for the protein-bound ligands.

| | Free ligands (PBS:MeCN 1:1) | | Protein bound ligands | |
|---|---|---|---|---|
| **Compound** | **Time to reach PSS @375 nm [min]** | **Time to reach PSS @545 nm [min]** | **Time to reach PSS @375 nm [min]** | **Time to reach PSS @450 nm [min]** |
| **1** | 6 | 2 | 75 | 10 |
| **2** | 6 | 2 | 75 | 30 |
| **3** | 6 | 2 | 75 | 25 |
| **4** | 6 | 2 | 75 | 20 |
| **5** | 6 | 2 | 75 | 20 |
| **6** | 10 | 4 | n.m. | n.m. |
| **7** | 10 | 4 | 30 | 10 |
| **8** | 6 | 2 | 75 | 10 |
| **9** | 6 | 2 | 75 | 10 |
| **10** | 6 | 2 | 30 | 10 |

**Table S3**: List and composition of buffers used in this work.

| **Buffer** | **Composition** |
|---|---|
| Spectroscopy buffer | 20 mM Tris-HCl pH 7.4, 100 mM NaCl, 0.1 mg·$ml^{-1}$ CHAPS |
| Ni-NTA chromatography wash buffer | 20 mM Tris HCl pH 7.4, 300 mM NaCl, 10 mM imidazole |
| Ni-NTA chromatography elution buffer | 20 mM Tris-HCl pH 7.4, 300 mM NaCl, 500 mM imidazole (150 mM imidazole for protein crystallization) |
| Size exclusion chromatography (SEC) buffer/Storage buffer | 20 mM Tris-HCl pH 7.4, 100 mM NaCl |
| DMEM | DMEM high glucose (4.5 g·$L^{-1}$), 10% fetal bovine serum, 2 mM L-glutamine, 1 mM sodium pyruvate, 100 U·$mL^{-1}$ penicillin, 100 μg·$mL^{-1}$ streptomycin |
| Imaging buffer (microscopy) | DMEM without phenol red high glucose (4.5 g·$L^{-1}$), 10% fetal bovine serum, 1 mM sodium pyruvate, 1 mM L-glutamine |

**Table S4**: Intensity values of the LED boxes used in this work.

| Wavelength [nm] | Light intensity |
|---|---|
| 375 | 700 μW·cm$^{-2}$ |
| 450 | 2 mW·cm$^{-2}$ |
| 545 | 700 μW·cm$^{-2}$ |

**Table S5**: List of fluorophores and their properties used in this work. Values were obtained from suppliers' website or original publication unless underwise noted.

| Name | $\lambda_{max}/\lambda_{em}$ [nm] | ε [cm$^{-1}$·M$^{-1}$] | $\Phi_F$ | Supplier |
|---|---|---|---|---|
| $JF_{549}$-BCN | 555/581* | 120000* | 0.79* | Home-made |
| $JF_{549}$-HTL[4] | 549/571 | 101000 | 0.88 | Gift from Lavis lab |
| CF®488-TCO | 490/515 | 70000 | - | Biotium |
| Cy3 TCO | 555/565 | - | - | Biomol |
| CF®647-TCO | 650/665 | 240000 | 0.43* | Biotium |
| $JF_{635}$-HTL (HaloTag bound)[2] | 635/652 | 81000 | 0.75 | Home-made |
| $JF_{635}$-HTL (free ligand)[5] | 635/652 | 400 | 0.56 | Home-made |
| $JF_{722}$-HTL[4] | 722/743 | 87200 | 0.77 | Gift from Lavis lab |

*measured at EMBL Heidelberg

**Table S6:** Plasmids used for cell culture and microscopy to express SNAP-tag−HaloTag fusion, with or without EGFP cytosolic expression.

| ***Cellular localization*** | ***Construct*** |
|---|---|
| Cell surface (extracellular) | pCDNA5_FRT_TO_IgKchL_HA_SNAP-tag_HaloTag_myc_PDGFRtmb |
| | pCDNA5_FRT_TO_IgKchL_HA_SNAP-tag_HaloTag_myc_PDGFRtmb_T2A_EGFP |
| Cytosol (intracellular) | pCDNA5_FRT_TO_SNAP-tag_HaloTag_T2A_EGFP |

**Table S7**: Characterization of purified proteins by NanoDSF.

| Purified protein | $T_{onset}$ [ºC] | $T_m$ [ºC] | $T_{agg}$ [ºC] |
|---|---|---|---|
| HaloTag | 52.0 | 61.6 | 56.7 |
| SNAP-tag-HaloTag | 50.1 | 59.5 | 47.6 |
| cp178HaloTag | 24.4 | 37.5 | 51.7 |

### Materials and Methods

#### Synthesis

Commercial reagents were obtained from reputable suppliers (e.g. Sigma-Aldrich, BLD Pharmatech, Tokyo Chemical Industry, abcr GmbH) and used as received. All solvents used for chemical reactions were of anhydrous grade, purchased in septum-sealed bottles stored under an inert atmosphere (Sigma-Aldrich). Synthetic-grade solvents (VWR) were used without further purification. Reactions were conducted in round-bottomed flasks, Schlenk flasks, Anton Paar G10 and G30 PTFE-coated-silicon-septum-capped microwave vials or septum-capped Biotage® reaction vials containing Teflon-coated magnetic stir bars. Reactions under an inert atmosphere were purged under vacuum/argon on a Schlenk line. Heating of reactions was accomplished using oil baths on top of a stirring hotplate equipped with an electronic contact thermometer to maintain the indicated temperatures.

Reactions were monitored by thin layer chromatography (TLC) on precoated aluminium plates (silica gel 60 F254, 200 µm thickness) or by LCMS (Agilent 1260 Infinity II; ZORBAX SB-C18 18 µm 80 Å, 2.1x50 mm column, 5 to 20 µL injection, 5–95% $CH_3CN/H_2O$ gradient with constant 0.1% v/v $HCO_2H$ additive, 8 to 10 min run, 0.6 mL/min flow, ESI positive ion mode, detection at 254 nm). TLC plates were visualized by UV illumination (254 nm, Konrad Benda - Herolab 220 V 2x6 Watt UV lamp) or developed with stains (e.g.phosphomolybdic acid stain). Compounds were purified by flash chromatography on an automated purification system (Biotage Isolera One) using pre-packed silica cartridges (Biotage Sfär Duo, 60 Å pores, 60 µm particles size) or by preparative HPLC (Agilent 1260 Infinity II, Phenomenex Gemini NX 21.2x150 mm, 5 µm C18 column). High-resolution mass spectrometry was performed by Dr. Mariana Natali Barcenas Rodriguez (Metabolomics Core Facility, EMBL Heidelberg).

The identity and purity of synthesized compounds were determined by NMR spectroscopy. $^{1}H$, $^{13}C$, and $^{19}F$ NMR spectra were recorded on a Brunker 400 UltraShield NMR and the frequencies used were 400, 101, and 376 MHz, respectively. Deuterated solvents were used as purchased (Deutero GmbH). $^{1}H$ and $^{13}C$ NMR chemical shifts (δ) were referenced to the residual solvent peak. Data for $^{1}H$ NMR spectra are reported as chemical shift (δ in ppm), multiplicity (s = singlet, d = doublet, t = triplet, q = quartet, p = pentuplet, dd = doublet of doublets, ddd= doublet of doublets of doublets, dt = doublet of triplets, tt = triplet of triplets, dtt = doublet of triplets of triplets, m = multiplet, br s = broad signal), coupling constant (*J* in Hz), integration. Data for $^{13}C$ NMR spectra are reported by chemical shift (δ in ppm) with hydrogen substitution information obtained from DEPT spectra (C, CH, $CH_2$, $CH_3$). Data for $^{19}F$ NMR are reported by chemical shift (δ in ppm) and coupling constant (*J* in Hz). Data was processed using Mnova from Mestrelab. The $^{13}C$ NMR spectra are not reported for compounds containing HaloTag or SNAP-tag ligands due to the limited amount of compound available, and the complexity of the spectra. Determination of sample concentration for UV-Vis and fluorescence spectroscopy was performed by $^{1}H$ NMR of samples in $(CD_3)_2SO$ containing 5 mM DMF for relative integration.

$JF_{635}$-HaloTag ligand and $JF_{549}$-HaloTag ligand were synthesized as previously described, and $^{1}H$ NMR and HPLC confirmed identity and purity of the compounds.[5,6] $JF_{722}$-HaloTag ligand, $JF_{635}$-SNAP-tag ligand and $JF_{549}$-cpSNAP-tag ligand were kindly provided by the Lavis group (Janelia Research Campus). The clickable fluorophores were obtained from reputable suppliers and used as received: CF®488A-TCO (Biotium) Cy3-TCO (Biomol) and CF®647-TCO (Biotium) The full list of fluorophores used can be found in Table S5. The composition of buffers used in this work can be found in Table S3.

#### UV-Vis and Fluorescence Spectroscopy

All solvents used for photophysical measurements were of spectroscopic grade. All measurements were performed at room temperature (23±2°C). DIO-ligands **1−10** were initially prepared as 2 mM stock solutions in DMSO, and subsequently diluted in appropriate solvents and buffers to ensure that the final DMSO concentration did not exceed 1% v/v. Unless otherwise specified, spectroscopic measurements with purified protein were performed in aqueous buffer containing 20 mM Tris-HCl and 100 mM NaCl at pH 7.4, to which 0.1 $mg{\cdot}mL^{-1}$ 3-((3-cholamidopropyl)dimethylammonio)-1-propanesulfonate (CHAPS) was added. Spectroscopic measurements were made in 1-cm path-length quartz cuvettes (Hellma Suprasil Quartz), or in 96-well plates (Nunc™ MicroWell™ 96-Well, Nunclon Delta-Treated, Flat-Bottom Microplate).

Absorption spectra were recorded using a Cary Model 60 spectrophotometer (Agilent). Absorption spectra, including maximum absorption wavelength ($\lambda_{max}$), extinction coefficient ($\varepsilon$), and maximum emission wavelength ($\lambda_{em}$), were measured in triplicates for the free DIO-ligands, and in duplicate for the protein-bound ligands. Spectra were recorded at 10 μM dye concentration for the free DIO-ligands and at 5 μM with the protein, unless otherwise stated. For measurements in the presence of proteins, the DIO ligand was incubated with 1.5 equivalents (eq) of purified protein for at least 2 h in the dark at room temperature while shaking.

Fluorescence spectra were recorded using a JASCO spectrofluorometer (FP-8500), at 5 μM dye concentration of and 7.5 μM protein concentration, unless otherwise stated.

Data analysis and graph plotting were performed with GraphPad Prism.

The following notations are used:

| | |
|---|---|
| $\lambda_{max}$ [nm] | Wavelength at maximal absorption |
| $\lambda_{em}$ [nm] | Wavelength at maximal fluorescence emission |
| $\varepsilon$ [$M^{-1}\cdot cm^{-1}$] | Molar extinction coefficient |
| $\Phi_F$ | Fluorescence quantum yield |
| $t_{!/2, relax}$ [min] | Half-life of the thermal relaxation in the dark, at room temperature |

#### Fluorescence Quantum Yield Measurement

Fluorescence quantum yields ($\Phi_F$) were measured using an absolute quantum yield measurement system (Quantaurus, C11347, Hamamatsu). Measurements were carried out using dilute samples so that absorbance did not exceed 0.1. Self-absorption corrections were performed using the Quantaurus software.

#### Photoswitching

Photoswitching was performed using a LED illumination device, designed and built by the Electronic and Mechanical Workshops at EMBL Heidelberg (Figure S7), which was re-adapted from our previous work.[2] The device consists of two main parts: the controller board, and the interchangeable LED boards which comprise 24 LEDs at a specific wavelength (LED450-03, LED375-04 or LED545-01, Roithner LaserTechnik). The illumination device is controlled using the RealTerm Serial terminal program, with the following adjustable parameters: light intensity, ON and OFF time (in ms), and number of ON/OFF cycles. For all experiments, the maximum light intensity of the system was used. Maximum values for the different LED boxes can be found in Table S4.

#### Thermal Relaxation Kinetics

Thermal relaxation kinetics were measured using a Cary Model 60 spectrophotometer (Agilent) at room temperature, in the dark, in 1-cm path-length quartz cuvettes (Hellma Suprasil Quartz), in aqueous buffer (20 mM Tris-HCl, 100 mM NaCl, 0.1 $mg\cdot mL^{-1}$ CHAPS, pH 7.4). The cuvette was illuminated at 375 nm (700 $\mu W\cdot cm^{-2}$) until the photostationary state was reached, measured by absorbance. Absorption spectra were then recorded every five minutes. The resulting relaxation half-lifes ($t_{1/2, relax}$) of the species formed upon illumination were determined by linear fitting of $\ln(A_0/A_t)$ as a function of time.

#### Photoswitching Cycles

Photoswitching cycles were measured using a Cary Model 60 spectrophotometer (Agilent) at room temperature, in the dark, in 1-cm path-length quartz cuvettes (Hellma Suprasil Quartz), in aqueous buffer (20 mM Tris-HCl, 100 mM NaCl, 0.1 $mg\cdot mL^{-1}$ CHAPS, pH 7.4). The cuvette was illuminated alternatively at 375 nm (700 µW·cm$^{-2}$) and 545 nm (700 µW·cm$^{-2}$) for 2 min. Absorption spectra were recorded after every illumination step, and the absorbance at $\lambda_{max}$ was plotted.

#### Labeling Efficiency Measurement by Pulse-Chase Assay *in vitro*

The labeling efficiency assay was performed using a microplate reader (TECAN Spark microplate reader) at room temperature, in the dark, in 96-well plates (Nunc™ MicroWell™ 96-Well, Nunclon Delta-Treated, Flat-Bottom Microplate), in aqueous buffer (20 mM Tris-HCl, 100 mM NaCl, 0.1 mg·mL$^{-1}$ CHAPS, pH 7.4). Ligands **1−10** (2 µM) were incubated with the purified protein tag (1 µM) for 2 h or 5 h in the dark, after which either $JF_{635}$-HaloTag ligand (1 µM) or $JF_{635}$-SNAP-tag ligand was added and the fluorescence of $JF_{635}$ ($\lambda_{max}/\lambda_{em}$ = 640/660 nm) was measured. If binding of the switch ligand to the protein was complete during the incubation period, no increase in far-red fluorescence is observed following addition of $JF_{635}$ ligand. The controls ($Halo_{635}$ and $SNAP_{635}$) were made by incubating the purified protein with $JF_{635}$-HaloTag ligand for 2 hours, or with $JF_{635}$-SNAP-tag ligand for 5 hours.

#### Click Reaction with *endo*-BCN Characterized by UV-Vis Spectroscopy

Two samples of ligand **1−10** (5 µM) were incubated with their respective protein tag (HaloTag for compounds **1−3** and **7−10**, SNAP-tag for compounds **4,5**, 7.5 µM) for two hours in the dark. To one sample, *endo*-BCN (10 µM) was added before illumination. Both samples were illuminated at 375 nm to reach the maximum (700 µW·cm$^{-2}$) and absorbance was monitored at $\lambda_{max}$. For both samples $\lambda_{max}$ was plotted against time. The experiment was performed in duplicate.

### Click Reaction with $JF_{549}$-BCN Characterized by SDS-PAGE Gel

Purified proteins (1 µM, HaloTag or HaloTag-SNAP-tag) were incubated with their respective ligands **1−10** (2 µM) for 5 hours in the dark. $JF_{549}$-BCN (**11,** 4 µM) was added and the samples were then divided into light and dark groups. The “light” samples were illuminated for 10 min at 375 nm (700 µW·cm$^{-2}$), while the “dark” samples were kept in the dark at room temperature for 10 min. To remove unreacted fluorophores, the samples were filtered using spin columns (Zeba Spin Desalting Plates, 7K MWCO, Thermo Scientific), mixed with Laemmli sample buffer (Biorad) supplemented with 50 mM DTT, denatured (98°C, 5 min), and loaded onto 4–20% Mini-PROTEAN TGX Stain-Free Protein Gels (Bio-Rad). Afterward, the gel was imaged using a fluorescence gel reader (Typhoon FLA 7000), followed by Coomassie staining and imaged again in a ChemiDoc imaging system (ChemiDoc™ Touch, version 2.3.0.07, Bio-Rad). As a positive control, the purified protein was labeled with $JF_{549}$-HaloTag ligand or $JF_{549}$-cpSNAP-tag ligand (2 µM), and incubated for the same duration as in the photoclick experiment. Analysis of the results was performed using ImageJ. Background fluorescence was subtracted, and the fluorescence signal was measured from manually drawn ROIs corresponding to the visible band on the Coomassie staining image. Normalization to account for protein concentration was performed by dividing the fluorescence signal by the Coomassie signal.

The impact of the concentration of clickable fluorophore **11**, illumination time and illumination intensity was evaluated *in vitro*. The samples were prepared as described above. As representative, ligand **3** (2 µM) was used and incubated with HaloTag (1 µM) for 2 h in the dark. The resulting normalized fluorescence intensity values were plotted and fitted by dose-response equation in GraphPad Prism ([Agonist] vs. response -- Variable slope (four parameters)).

$$\mathrm{Y = Bottom + (X^\wedge Hillslope) * (Top - Bottom)/(X^\wedge HillSlope + EC50^\wedge HillSlope)}$$

### Vectors and Cloning

For cloning, the nucleic acid sequence was amplified by polymerase chain reaction (PCR) and inserted into the desired vector backbone via Gibson assembly (New England Biolabs) and purified by DNA Clean & Concentrator-5 (Zymo Research). Plasmid DNA was subsequently electroporated in *E. coli* Turbo cells (New England Biolabs) plated on LB agar plates with appropriate antibiotics and incubated at 37°C overnight. In all cases, plasmid DNA was extracted from individual bacterial colonies (QIAwave Plasmid Miniprep Kit, Qiagen), and the sequences of generated plasmids were verified by Sanger sequencing (Eurofins Genomics).

pCDNA5_FRT_TO_IgKchL_HA_HaloTag7_myc_PDGFRtmb (Addgene, #175534), pCDNA5_FRT_TO_LifeAct_HaloTag7_T2A_EGFP (Addgene, #135445), pET51b_His_TEV_HaloTag7 (Addgene #167266) and pET51b_StrepTagII_Enterokinase_SNAP-tag_PreScission_HaloTag7 vectors were kindly provided by the Johnsson group (Max Planck Institute for Medical Research). pET51b_StrepTagII_Enterokinase_SNAP-tag_Prescission_HaloTag7 plasmid has a StrepTagII and an Enterokinase cleavage site before SNAP-tag, followed by a PreScission cleavage site (GRLEVLFQGPKAFLE amino acid sequence) before HaloTag7. The SNAP-tag sequence is the same one as in plasmid pET51b_His_TEV_SNAP-tag (Addgene #167269).

We generated the pCDNA5_FRT_TO_IgKchL_HA_SNAP-tag_HaloTag_myc_PDGFRtmb vector by inserting the amplified SNAP-tag-PreScission fragment (from the pET51b_StrepTagII_Enterokinase_SNAP-tag_Prescission_HaloTag7 vector) into the pCDNA5_FRT_TO_IgKchL_HA_HaloTag_myc_PDGFRtmb vector (Addgene #175534). For the EGFP co-expression in mammalian cells, we generated the pCDNA5_FRT_TO_IgKchL_HA_SNAP-tag_HaloTag_myc_PDGFRtmb_T2A_EGFP vector by inserting the amplified T2A_EGFP fragment (from pCDNA5_FRT_TO_LifeAct_HaloTag7_T2A_EGFP, Addgene, #135445) into the pCDNA5_FRT_TO_IgKchL_HA_SNAP-tag_HaloTag_myc_PDGFRtmb vector.

In addition, we generated the pET51b_His_TEV_cp178HaloTag vector by Gibson assembly. Here, the HaloTag7 was replaced by the circularly permuted HaloTag 177/180,[2,3] bearing a (GGTGGS)x3 linker in between and 2 extra amino acids at the end of the sequence (SG), in the pET51b_His_TEV_HaloTag7 (Addgene #167266) vector.

**Protein Expression and Purification**

The pET51b vectors (pET51b_His_TEV_HaloTag7 and pET51b_StrepTagII_Enterokinase_SNAP-tag_Prescission_HaloTag7) were used in *E. coli* BL21(DE3) for protein expression (HaloTag and SNAP-tag-HaloTag, respectively). LB cultures containing 100 $\mu g \cdot mL^{-1}$ ampicillin were grown at 37°C to an optical density at 600 nm (OD600) of 0.4−0.6, induced by the addition of IPTG (1 mM final concentration) for protein expression and further grown at 18°C overnight (shaking 200 rpm). The cells were harvested by centrifugation (6000 rpm, 20 min, 4°C) and lysed by microfluidizer (Microfluidics Microfluidizer, M-110L FluidProcessor) in lysis buffer (20 mM Tris-HCl pH 7.4, 300 mM NaCl, 1 mM PMSF, ½ tablet of protease inhibitor, 0.01 $mg \cdot mL^{-1}$ DNase). The cell lysate was cleared by ultra-centrifugation (35000 rpm, 45 min, 4°C). Proteins were purified using affinity-tag Ni-NTA agarose (Qiagen) and the eluted fractions (20 mM Tris-HCl pH 7.4, 300 mM NaCl, 500 mM imidazole) were then pooled. Size exclusion chromatography (SEC) (HiLoad 16/600 Superdex 200 pg, Cytiva) was performed to obtain the monomeric fraction with elution using SEC buffer (20 mM Tris-HCl, 100 mM NaCl, pH 7.4). This final protein fraction was concentrated (Amicon Ultra Centrifugal Filter, 30 kDa MWCO, Merck) to 100-170 µM and stored in 100 µL aliquots at -70°C. These proteins were also further characterized by NanoDSF (Prometheus NT.Flex, Table S7), mass photometry (Refeyn Two$^{MP}$ from Refeyn), and SDS-PAGE gel (Bio-Rad, Mini-PROTEAN® TGX™ stain-free gels).[2]

**Structure Prediction**

ColabFold is an accessible tool for predicting protein structures that combines the fast homology search capabilities of MMseqs2 with AlphaFold2 or RoseTTAFold algorithms.[7] ColabFold also gives important confidence metrics for evaluation of the predicted protein structure: pLDDT (predicted Local Distance Difference Test) and PAE (Predicted Aligned Error). For 3D protein structure prediction, protein sequences were inputted into ColabFold version 1.5.2 (https://github.com/sokrypton/ColabFold) installed on the local HPC cluster through EasyBuild environment module (https://docs.easybuild.io/version-specific/supported-software/c/ColabFold/) with the help of Federico Marotta (Bork Group, EMBL Heidelberg). For each sequence, five models were generated and ranked by their pLDDT scores. The predicted structures were visualized and analyzed using PyMOL.

#### Cell Culture

U2OS cells (ATCC) were cultured in Dulbecco's modified Eagle medium (DMEM, high glucose 4.5 g·L$^{-1}$) with phenol red (Gibco), supplemented with 10% v/v fetal bovine serum (ThermoFisher Scientific), penicillin (100 units·mL$^{-1}$), streptomycin (100 µg·mL$^{-1}$, Gibco), 2 mM·L$^{-1}$ Glutamine (Gibco) and 1 mM sodium pyruvate (Gibco), and maintained at 37°C in a humidified 5% v/v $CO_2$ environment.

Transient transfections were performed using FUGENE6 (Promega) transfection reagent according to the manufacturer´s recommendations ratio (1.5 µg plasmid DNA, 4.5 µL FUGENE6) in OptiMEM (Gibco) and added to wells (Nunclon delta surface 6 well plates, Nunc) with 70−80% cell confluency in imaging buffer (DMEM without phenol red (Gibco), supplemented with glucose (4.5 g·L$^{-1}$), 10% v/v fetal bovine serum (ThermoFisher Scientific), 2 mM L-Glutamine (Gibco) and 1 mM sodium pyruvate (Gibco)) for 24 h. Transfected cells were then trypsinized, counted, and seeded into imaging plates (cellview cell culture slide, PS, 75/25 mm, glass bottom, 10 compartments, TC, sterile, single packed (Greiner, #543078); 6 channel µ-Slide VI 0.5 Glass Bottom, Glass coverslips, No. 1.5H, selected quality, 170 µm±5 µm (Ibidi, #80607); Imaging Plate 96 CG, TC-Surface, 145 µm Cover Glass Bottom (Zellkontakt, #5241-20). The labeling and imaging were conducted the following day.

##### Labeling Efficiency Assay

U2OS cells co-expressing SNAP-tag-HaloTag on the cell surface and EGFP in the cytosol, or both SNAP-tag-HaloTag and EGFP in the cytosol, were incubated in the absence or presence of ligands **1-10** (2 µM) for 2 h at 37°C, and washed 3 times with prewarmed imaging buffer. $JF_{635}$-HaloTag ligand or $JF_{635}$-SNAP-tag ligand (1 µM) was then added and the cells were incubated for another 30 min, together with the negative control (transfected cells with neither ligand nor dye-ligand). After washing with prewarmed imaging buffer the cells were imaged on a Nikon Ti-E microscope (see below). Far-red fluorescence increase after the addition of $JF_{635}$ ligand increase indicates incomplete binding by the DIO-ligands.

##### MTS assay

U2OS cells transfected with pCDNA5_FRT_TO_IgKchL_HA_SNAP-tag_HaloTag_myc_PDGFRtmb vector (1·10$^4$ cells per well) were seeded in triplicates in 96-well plates (Nunclon Delta surface, Nunc) and incubated overnight to allow the cells to attach to the surface of the wells. The cells were then incubated with 1 µM ligand **1, 7** or **10** for 2 h, washed 3 times with prewarmed imaging buffer and illuminated for 5 min at 375 nm (700 µW·cm$^{-2}$). Cell viability was determined using an MTS assay (Abcam Inc.) according to the supplier's instructions. Briefly, 20 µL of MTS reagent was added (to each well) to both the non-illuminated (control plate) and the illuminated plate. After 1h 45 min, absorbance at 490 nm was measured on a microplate reader (TECAN Spark).

##### Cellular Labeling

Cells expressing either only SNAP-tag-HaloTag on the cell surface, or co-expressing SNAP-tag-HaloTag on the cell surface and EGFP in the cytosol, were labeled with respective ligands **1−10** (1 µM, 2 to 3 h at 37°C, in the dark) in imaging buffer and subsequently washed three to five times with prewarmed imaging buffer before fluorescence imaging.

#### Flow cytometry

Transfected U2OS cells co-expressing HaloTag-SNAP-tag fusion on the cell surface and EGFP in the cytosol were labeled with ligands **1−10** (1 µM, 2-3 h at 37°C, in the dark) in imaging buffer directly in a

6-well plate (Nunclon delta surface 6 well plates, Nunc) and subsequently washed four times with prewarmed imaging buffer. CF®647-TCO (2 μM) was then added and incubated at 37°C for 5 min. The plate was then illuminated at 375 nm (700 $\mu W \cdot cm^{-2}$) for 5 min, followed by washing with prewarmed imaging buffer 5 times. Ice-cold PBS (2 × 600 μL per sample) was added to the wells and the cells were gently detached from the well surface using cell scrapers (Sigma, Greiner cell scrapers, C5981-100EA). The cells were transferred to an Eppendorf vial and centrifuged (1500 rpm, 5 min), followed by resuspending the cell pellet in ice-cold PBS (300 μL per sample).
As controls, the following samples were additionally prepared: (i) non-transfected cells, to determine the gates for relative size and complexity (FSC-A SSC-A); (ii) unlabeled transfected cells to determine the positive gate for EGFP+ cells and within those cells also was determined the background signal on the 670/30 bandpass filter; (iii) transfected cells labeled with $JF_{635}$-HaloTag ligand (1 μM) as positive control for fluorescence detected at 670/30.

The flow cytometry experiment was performed on a FACS Symphony A3 (BD, San Jose , CA) equipped with the Facs Diva 10 software. EGFP was excited with a 100 mW 488 nm laser and the fluorescence emitted was acquired on a PMT tube equipped with a 530/30 bandpass filter. CF®647 was excited with a 100 mW 640nm laser and the fluorescence emitted was acquired on a PMT tube equipped with a 670/30 bandpass filter. The data was analyzed using Flowjo Version 10.

#### Microscopy

Widefield imaging was performed at the Advanced Light Microscopy Facility (ALMF, EMBL Heidelberg), on a Nikon Ti-E microscope equipped with a Spectra X light engine (Lumencore) with a 40x objective (CFI P-Apo 40x Lambda/0.95/0.25−0.16) and imaged onto a Hamamatsu Orca Flash 4 V2 camera (pixel size: 6.5 $\mu$m/2048 x 2048 pixel). The microscope was equipped with a $CO_2$ and temperature-controllable incubator (home-built, 37°C). A quad bandpass filter cube was used to image EGFP (excitation 485/20, emission 525/50) and $JF_{635}$-HaloTag ligand/$JF_{635}$-SNAP-tag ligand/CF®647-TCO (excitation 650/13, emission 692/40).

Multicolor imaging was performed at the Advanced Light Microscopy Facility (ALMF, EMBL Heidelberg), on an Evident FV3000 Confocal laser scanning microscope with 4 GaAsP spectral detectors, lasers: 405, (445), 488, (514), 561, (594), 640 nm, hardware autofocus (ZDC), motorized IX3-SSU xy-stage and a UPLXAPO 40x/0.95 Corr air objective. Laser stimulation settings (“FRAP” settings) were used to scan a selected ROI with 405 nm at 0.5% power level with 4μs/pixel (41 $kW \cdot cm^{-2}$) for 20 seconds, during which the ROI was scanned several times.

#### Widefield and Confocal Image analysis

Statistical analysis and curve fitting were performed using GraphPad Prism. All images were processed with ImageJ/Fiji and macros written therein unless otherwise stated.[8,9]

For segmentation and analysis of the widefield images of labeling of entire field of view, Arif Khan (Bioimage Analysis Support Team at the EMBL Data Science Centre) re-adapted a Python script from our previous work.[2] The code is based on cell segmentation of the cytosolic EGFP signal using a model trained on top of Cellpose model ‘cyto2’.[10] To compare the basal brightness of the clicked state, the fluorescence intensity was divided by the cytosolic EGFP intensity. The resulting cell labels were matched across images. The fluorescence intensity for the Cy3 and Cy5 channel were determined for each cell. Dead cells were excluded from the analysis. To compare the basal brightness of the clicked state, the fluorescence intensity was divided by the cytosolic EGFP intensity. The code is available at https://git.embl.org/grp-cba/htlov-mitochondria-intensity-quantification/-/tree/main/code/python?ref_type=heads, and the instructions to run the code is available at https://git.embl.org/grp-cba/htlov-mitochondria-intensity-quantification/-/blob/main/analysis_workflow.md?ref_type=heads.

For analysis of the two- and three-color experiments, ROIs corresponding to single cells or line profiles were manually drawn in ImageJ/Fiji. Background was substracted, and the intensity in all channels was plotted using GraphPad Prism.

#### Microfluidics Setup

A custom microfluidics setup was kindly provided by Franziska Kundel (Ellenberg group, EMBL Heidelberg).[11] The setup consists of a commercial 3-axis GRBL-controlled CNC stage (30 × 18 cm) adapted by mounting a syringe needle in place of the drill head. This needle is connected via 1-mm inner diameter PEEK and silicone tubing (VWR) to a male Luer adapter interfacing with the sample microslide. The microslide's output tubing is connected either to a CPP1 peristaltic micropump (Jobst Technologies, Freiburg, Germany) running at full speed and delivering about 1 $mL \cdot min^{-1}$. Dye solutions and imaging buffer (for washing) are stored in falcon tubes. Automated control is implemented via custom Python software on a PC which communicates with the MP-X controller, the Jobst Pump Driver, and the GRBL CNC board. The channel was flushed with the respective dye for 40 s, followed by adding imaging buffer for 15 s. After imaging the channel was washed by flushing for 60 s with imaging buffer. Changes of buffer and fluorophore solutions were performed manually.

#### Two-Color Photoclick on Cell Surface

U2OS cells expressing a HaloTag-SNAP-tag fusion on the cell surface were labeled with ligand **7** (1 µM) for 2 h at 37°C in the dark. The cells were washed for three times with prewarmed imaging buffer. The respective chamber was connected to the microfluidics device and washed for 60 s with imaging buffer. ROIs were selected and an image before photoclick reaction was taken. The respective dye (CF®647-TCO or Cy3-TCO, 2 µM in imaging buffer) was added to the channel by flushing for 40 s, followed by 15 s for the dead volume. The respective ROIs were activated with the microscope (405 nm, 4µs/pixel, 0,5% power level (41kW $cm^{-2}$), 20 s). The channel was washed with imaging buffer by flushing for 60 s. The procedure was then repeated with another dye.

#### Three-Color Photoclick on Cell Surface

Co-labeling with $JF_{722}$-HaloTag ligand was used to identify transient transfected cells (as EGFP could not be used as it blocks use of the 488 channel). Cells expressing a HaloTag-SNAP-tag fusion on the cell surface were labeled with a mixture of $JF_{722}$-HaloTag ligand (100 nM) and ligand **7** (1 µM) in imaging buffer for 1.5 h at 37°C in the dark. Subsequently, the cells were washed three to five times with prewarmed imaging buffer before fluorescence imaging. The respective chamber was connected to the microfluidics device and washed for 60 s with imaging buffer. ROIs were selected by searching for dim labeling in the far-red channel and an image before photoclick reaction was taken. The dye (CF®647-TCO, CF®488ATCO or Cy3-TCO, 2 µM in imaging buffer) was then added to the channel by flushing for 40 s, followed by 15 s for the dead volume. The respective ROIs were activated with the microscope (405 nm, 4µs/pixel, 0,5% power level (41kW $cm^{-2}$), 20 s). The channel was washed with imaging buffer by flushing for 60 s. The procedure was then repeated with another dye.

### Synthesis

**Table S7:** Known precursors synthesized for this project with corresponding reference.

| Name | Structure | Reference |
|---|---|---|
| **1-oxo-2,3-diphenyl-1H-indene-6-carboxylate** | 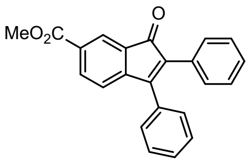 | 1 |
| **6-bromo-2,3-diphenyl -1H-inden-1-one** | 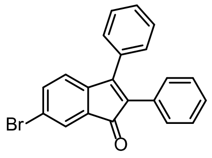 | 1 |
| **1,2-di(naphthalen-2-yl)ethyne** | 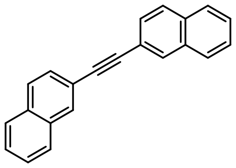 | 12 |
| **1,2-bis(4-(trifluoromethyl)phenyl)ethyne** | 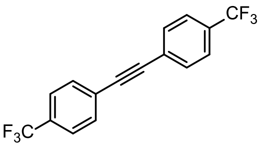 | 13 |
| **1,2-bis(3-(trifluoromethyl)phenyl)ethyne** | 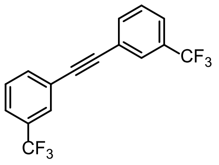 | 13 |
| **BCN-amine** | 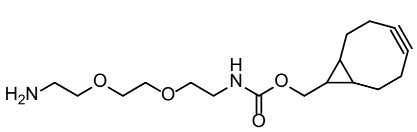 | 14 |
| **6-tert-butoxycarbonyl-$JF_{549}$** | 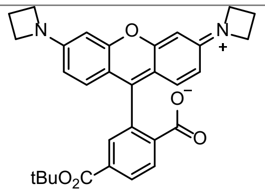 | 15 |
| **$JF_{549}$-NHS ester** | 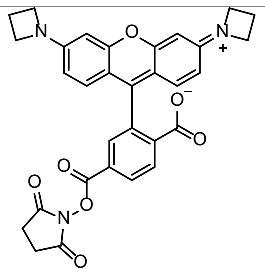 | 16 |
| **ethanamine, 2-[2-[2-[(6-chlorohexyl)oxy]ethoxy]ethoxy] (TFA salt)** | 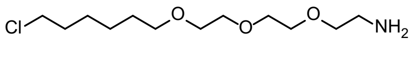 | 17 |

**Synthesis of precursors**

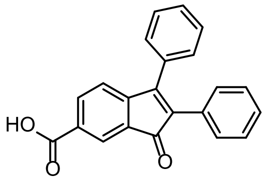

**1-Oxo-2,3-diphenyl-1H-indene-6-carboxylic acid (S1)**: 1-oxo-2,3-diphenyl-1H-indene-6-carboxylate[1] (100 mg, 294 µmol, 1.0 eq) was dissolved in a mixture of MeOH:THF (1:1, 20 mL). NaOH (1 M in $H_2O$, 1.2 mL, 1.2 mmol, 4.0 eq) was added and the reaction was stirred at room temperature overnight. The solvent was removed under reduced pressure and the crude material was dissolved in EtOAc. The organic phase was washed with water, dried over $Na_2SO_4$ and the solvent was removed under reduced pressure. The product was used without purification (80 mg, 0.24 mmol, 83%). **$^1$H NMR** ($(CD_3)_2SO$, 400 MHz) δ 8.03 – 7.97 (m, 2H), 7.52 – 7.15 (m, 10H), 7.07 (d, *J* = 7.9 Hz, 1H). **$^{13}$C NMR** ($(CD_3)_2SO$, 101 MHz) δ 195.9 (C), 168.1 (C), 155.1 (C), 144.7 (C), 142.3 (C), 134.6 (CH), 132.4 (C), 132.2 (C), 130.7 (C), 129.7 (CH), 129.5 (CH), 129.3 (C), 128.9 (CH), 128.3 (CH), 128.0 (CH), 127.7 (CH), 123.5 (CH), 120.5 (CH), 69.8 (C). **HRMS** (ESI) calcd for $C_{22}H_{14}O_3$ [M+H] 327.1016, found 327.1012.

**General procedure for preparation 2,3-diphenyl-1H-inden-1-one precursors *via* palladium catalyzed annulation of substituted 1,2-diphenylethyne derivatives (General Method M1):** The following procedure for **S2** is shown as representative. A pressure tube was charged with methyl 4-bromo-3-formylbenzoate (100 mg, 411 µmol, 1 eq), $Pd(OAc)_2$ (4.6 mg, 21 µmol, 0.05 eq ), NaOAc (135 mg, 1.65 mmol, 4 eq), n-$Bu_4NCl$ (114 mg, 411 µmol, 1 eq), and 1,2-di(naphthalen-2-yl)ethyne[12] (115 mg, 411 µmol, 1 eq). The tube was evacuated and backfilled with argon three times. Anhydrous DMF (4 mL) was added under an atmosphere of inert gas. The mixture was stirred at 100°C overnight. The reaction mixture was allowed to cool down to room temperature, diluted with water, and extracted with EtOAc (3x). The organic phase was washed with $NH_4Cl$ and dried over $Na_2SO_4$. The crude was purified by silica gel flash chromatography (0–5%, EtOAc/cyclohexane, linear gradient) affording **S2** as a light red solid (127 mg, 70%).

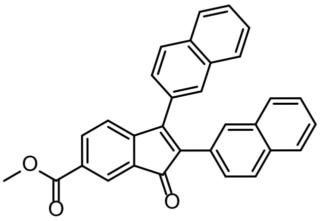

**Methyl 2,3-di(naphthalen-2-yl)-1-oxo-1H-indene-6-carboxylate (S2)**: **$^1$H NMR** ($CDCl_3$, 400 MHz) δ 8.27 (d, *J* = 1.6 Hz, 1H), 8.16 (dd, *J* = 7.6, 1.6 Hz, 1H), 8.04 (s, 2H), 7.90 – 7.78 (m, 4H), 7.77 – 7.72 (m, 1H), 7.63 – 7.53 (m, 3H), 7.50 – 7.41 (m, 2H), 7.39 – 7.33 (m, 2H), 7.20 (dd, *J* = 8.6, 1.7 Hz, 1H), 3.97 (s, 3H). **$^{13}$C NMR** ($CDCl_3$, 101 MHz) δ 195.4 (C), 166.3 (C), 154.4 (C), 149.8 (C), 135.8 (CH), 134.6 (C), 133.8 (C), 133.3 (C), 133.1 (C), 131.2 (C), 131.1 (C), 130.4 (CH), 130.0 (C), 128.9 (C), 128.8 (CH), 128.6 (CH), 128.3 (CH), 128.1 (CH), 127.9 (CH), 127.8 (CH), 127.7 (CH), 127.4 (CH), 127.3 (CH), 126.9 (CH), 126.8 (CH), 126.3 (CH), 126.2 (CH), 123.9 (CH), 121.2 (CH), 52.5 ($CH_3$). **HRMS** (ESI) calcd for $C_{31}H_{20}O_3$ [M+H] 441.1485, found 441.1480.

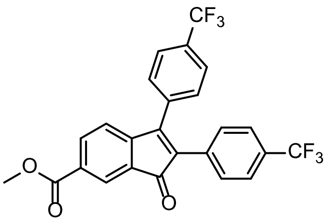

**Methyl 1-oxo-2,3-bis(4-(trifluoromethyl)phenyl)-1H-indene-6-carboxylate (S3):** The title compound **S3** was synthesized from methyl 4-bromo-3-formylbenzoate (100 mg, 411 µmol, 1 eq) and 1,2-bis(4-(trifluoromethyl)phenyl)ethyne[13] (130 mg, 2411 µmol, 1 eq) according to **general method M1**. Purification by silica chromatography (0–10% EtOAc/cyclohexane, linear gradient) afforded **S3** as yellow solid (88 mg, 45%). **$^{1}$H NMR** ($CDCl_3$, 400 MHz) δ 8.25 (d, *J* = 1.5 Hz, 1H), 8.16 (dd, *J* = 7.7, 1.6 Hz, 1H), 7.74 (d, *J* = 8.1 Hz, 2H), 7.53 (dd, *J* = 19.8, 8.1 Hz, 4H), 7.37 (d, *J* = 8.1 Hz, 2H), 7.20 (d, *J* = 7.7 Hz, 1H), 3.96 (s, 3H). **$^{13}$C NMR** ($CDCl_3$, 101 MHz) 194.1 (C), 165.9 (C), 154.3 (C), 148.6 (C), 136.1 (CH), 135.6 (C), 134.2 (C), 133.5 (C), 131.9 (C), 130.5 (C), 130.4 (CH), 129.0 (CH), 126.5 (t, *J* = 3.7 Hz, $CF_3$), 125.5 (t, *J* = 3.9 Hz, $CF_3$), 124.4 (CH), 121.4 (CH), 52.7 ($CH_3$). **HRMS** (ESI) calcd for $C_{25}H_{14}F_6O_3$ [M+H] 477.0916 found 477.0916.

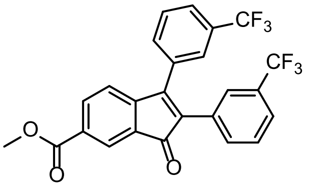

**Methyl 1-oxo-2,3-bis(2-(trifluoromethyl)phenyl)-1H-indene-6-carboxylate (S4):** The title compound **S4** was synthesized from methyl 4-bromo-3-formylbenzoate (100 mg, 411 µmol, 1 eq) and 1,2-bis(3-(trifluoromethyl)phenyl)ethyne[13] (130 mg, 411 µmol, 1 eq) according to **general method M1**. Purification by silica chromatography (0–5% EtOAc/cyclohexane, linear gradient) followed by a second column (20–80% $CH_2Cl_2$/cyclohexane, linear gradient) afforded **S4** as yellow solid (87 mg, 45%). **$^{1}$H NMR** ($CDCl_3$, 400 MHz) δ 8.26 (d, *J* = 1.6 Hz, 1H), 8.18 (dd, J = 7.6, 1.6 Hz, 1H), 7.74 (d, *J* = 7.6 Hz, 1H), 7.67 – 7.52 (m, 4H), 7.52 – 7.38 (m, 3H), 7.21 (d, *J* = 7.7 Hz, 1H), 3.96 (s, 3H). **$^{13}$C NMR** ($CDCl_3$, 101 MHz) δ 194.1 (C), 165.9 (C), 153.9 (C), 148.5 (C), 136.1 (CH), 134.1 (C), 133.3 (CH), 132.7 (CH), 132.2 (C), 131.9 (C), 131.2 (C), 130.9 (C), 130.6 (C), 130.5 (C), 130.1 (CH), 129.1 (CH), 126.8 (m, $CF_3$), 125.4 (m, $CF_3$), 124.4 (CH), 121.2 (CH), 52.6 ($CH_3$)z. **HRMS** (ESI) calcd for $C_{25}H_{14}F_6O_3$ [M+H] 477.0916, found 477.0916.

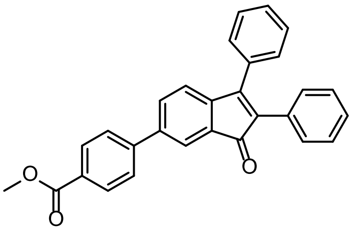

**Methyl 4-(1-oxo-2,3-diphenyl-1H-inden-6-yl)benzoate (S5):** A pressure tube was charged with 6-bromo-2,3-diphenyl-1H-inden-1-one[1] (100 mg, 277 µmol, 1 eq), (4-(methoxycarbonyl)phenyl)boronic acid (75 mg, 420 µmol, 1.5 eq), $Pd(PPh_3)_4$ (16 mg, 14 µmol, 0.05 eq ) and $K_2CO_3$ (77 mg, 0.55 mmol, 2 eq). The tube was evacuated and backfilled with argon three times. A mixture of dioxane:water (1:1, 10 mL) was added under an atmosphere of inert gas. The mixture was stirred at 70°C for 3 h. The reaction mixture was allowed to cool down to room temperature, diluted with $CH_2Cl_2$, and washed with water (2x)

and brine (1x). The organic phase was dried over $Na_2SO_4$. The crude was purified by silica gel flash chromatography (0–5% EtOAc/cyclohexane, linear gradient) affording the product **S5** as a light red solid (76 mg, 66%). **$^1$H NMR** ($CDCl_3$, 400 MHz) δ 8.17 – 8.09 (m, 2H), 7.88 (d, *J* = 1.7 Hz, 1H), 7.73 – 7.67 (m, 2H), 7.65 (dd, *J* = 7.6, 1.8 Hz, 1H), 7.48 – 7.38 (m, 5H), 7.27 (d, *J* = 10.5 Hz, 6H), 3.95 (s, 3H). **$^{13}$C NMR** ($CDCl_3$, 101 MHz) δ 175.8 (C), 167.0 (C), 144.4 (C), 141.1 (C), 132.8 (C), 132.1 (CH), 131.9 (C), 130.8 (C), 130.4 (CH), 130.1 (CH), 129.6 (CH), 129.0 (CH), 128.7 (CH), 128.3(CH), 128.1 (C), 126.9 (CH), 122.1 (CH), 121.9 (CH), 52.4 ($CH_3$). **HRMS** (ESI) calcd for $C_{29}H_{20}O_3$ [M+H] 417.1485, found 417.1476.

**General procedure for preparation 2,3-diarylepoxy indenones (General Method M2):** The following procedure for **S6** is shown as representative. The title compounds were prepared from **S1** according to the procedure of Xie *et al.*[1] **S1** (100 mg, 306 µmol, 1.0 eq) was dissolved in MeOH (5 mL). Hydrogen peroxide was added (17.5% in $H_2O$, 0.2 mL, 1 mmol) dropwise, followed by the slow addition of NaOH (1 M, 0.1 mL, 0.1 µmol). The reaction was stirred at room temperature until colorless. The solvent was removed under reduced pressure and the crude material was dissolved in EtOAc. The organic phase was washed with water, dried over $Na_2SO_4$ and the solvent was removed under reduced pressure. Purification by silica chromatography (0–10% EtOAc/cyclohexane, linear gradient) afforded **S6** as light orange, foamy solid (86 mg, 82%).

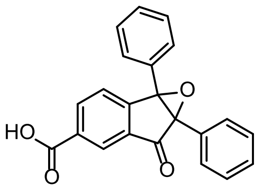

**6-Oxo-1a,6a-diphenyl-1a,6a-dihydro-6H-indeno[1,2-b]oxirene-4-carboxylic acid (S6): $^1$H NMR** ($CDCl_3$, 400 MHz) δ 8.57 (d, *J* = 1.6 Hz, 1H), 8.29 (dd, *J* = 7.8, 1.6 Hz, 1H), 7.46 (d, *J* = 7.8 Hz, 1H), 7.37 – 7.34 (m, 2H), 7.29 – 7.19 (m, 8H). **$^{13}$C NMR** ($CDCl_3$, 101 MHz) δ 195.2 (C), 170.6 (C), 153.9 (C), 136.3 (CH), 135.5 (C), 131.4 (C), 130.4 (CH), 129.2 (CH), 129.2 (C), 128.9 (CH), 128.7 (CH), 128.6 (CH), 128.3 (CH), 128.1 (CH), 127.9 (CH), 127.6 (C), 127.4 (CH), 125.8 (CH), 72.15 (C), 72.05 (C). **HRMS** (ESI) calcd for $C_{22}H_{14}F_6O_4$ [M+H] 343.0965, found 343.0960.

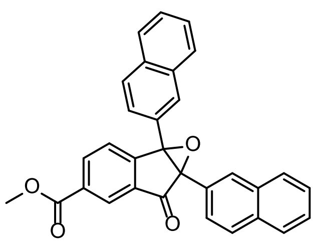

**Methyl 1a,6a-di(naphthalen-2-yl)-6-oxo-1a,6a-dihydro-6H-indeno[1,2-b]oxirene-4-carboxylate (S7):** The title compound **S7** was synthesized from **S2** (20 mg, 45 µmol, 1 eq) according to **<u>general method M2</u>**. Purification by silica chromatography (0–10% EtOAc/cyclohexane, linear gradient) afforded **S6** as a yellow foamy solid (15 mg, 72%). **$^1$H NMR** ($CDCl_3$, 400 MHz) δ 8.61 (dd, *J* = 1.6, 0.7 Hz, 1H), 8.31 (dd, *J* = 7.8, 1.6 Hz, 1H), 8.01 (d, *J* = 1.7 Hz, 1H), 7.92 (d, *J* = 1.7 Hz, 1H), 7.75 (s, 6H), 7.51 (dd, *J* = 7.9, 0.7 Hz, 1H), 7.49 – 7.39 (m, 6H), 3.99 (s, 3H). **$^{13}$C NMR** ($CDCl_3$, 101 MHz) δ 195.5 (C), 165.9 (C), 153.1 (C), 135.9 (CH), 135.4 (C), 133.5 (C), 133.4 (C), 133.0 (C), 132.9 (C), 132.3 (C), 128.5 (CH), 128.2 (CH), 128.2 (CH), 128.2 (CH), 128.1 (CH), 127.9 (CH), 127.8 (CH), 126.9 (CH), 126.8 (CH), 126.6 (CH), 126.4 (CH), 125.8 (CH), 125.2 (C), 124.9 (CH), 124.8 (CH), 72.7 (C), 72.5 (C), 52.8 ($CH_3$). **HRMS** (ESI) calcd for $C_{31}H_{20}O_4$ [M+H] 457.1434, found 457.1432.

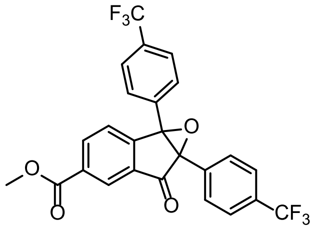

**Methyl 6-oxo-1a,6a-bis(4-(trifluoromethyl)phenyl)-1a,6a-dihydro-6H-indeno[1,2-b]oxirene-4-carboxylate (S8):** The title compound **S8** was synthesized from **S3** (20 mg, 45 µmol, 1 eq) according **general method M2**. Purification by silica chromatography (0–10% EtOAc/cyclohexane, linear gradient) afforded **S8** as yellow foamy solid (6 mg, 29%)· $^{1}$**H NMR** ($CDCl_3$, 400 MHz) δ 8.57 (d, *J* = 1.5 Hz, 1H), 8.32 (dd, *J* = 7.9, 1.6 Hz, 1H), 7.64 – 7.60 (m, 2H), 7.59 – 7.53 (m, 4H), 7.45 (t, *J* = 7.6 Hz, 3H), 3.99 (s, 3H). $^{13}$**C NMR** ($CDCl_3$, 101 MHz) δ 194.0 (C), 165.6 (C), 151.7 (C), 136.2 (CH), 135.1 (C), 133.0 (C), 132.8 (C), 131.9 (C), 131.5 (C), 131.5 (C), 131.3 (C), 128.5 (CH), 128.3 (CH), 127.2 (CH), 125.9 (t, *J* = 3.8 Hz, $CF_3$), 125.6 (CH), 125.5 (t, *J* = 3.7 Hz, $CF_3$), 122.6 (C), 122.4 (C), 72.1 (C), 71.6 (C), 52.9 ($CH_3$). **HRMS** (ESI) calcd for $C_{25}H_{14}F_6O_3$ [M+H] 493.0869, found 493.0864.

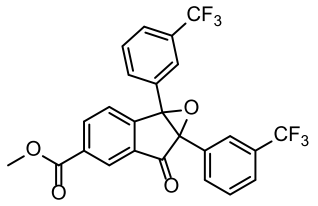

**Methyl 6-oxo-1a,6a-bis(3-(trifluoromethyl)phenyl)-1a,6a-dihydro-6H-indeno[1,2-b]oxirene-4-carboxylate (S9):** The title compound **S9** was synthesized from **S5** (30 mg, 63 µmol, 1 eq) according **general method M2**. Purification by silica chromatography (0–10% EtOAc/cyclohexane, linear gradient) afforded **S9** as white solid (16 mg, 52%).$^{1}$**H NMR** ($CDCl_3$, 400 MHz) δ 8.58 (s, 1H), 8.37 – 8.31 (m, 1H), 7.70 (s, 1H), 7.64 – 7.41 (m, 8H), 3.99 (s, 3H). $^{13}$**C NMR** ($CDCl_3$, 101 MHz) δ 193.9 (C), 165.6 (C), 151.4 (C), 136.3 (CH), 135.2 (C), 132.8 (C), 131.6 (C), 131.4 (CH), 131.3 (C), 131.2 (C), 131.1 (CH), 130.9 (C), 130.2 (C), 129.4 (CH), 129.0 (CH), 128.4 (C), 127.2 (CH), 126.2 (m, $CF_3$), 125.6 (CH), 125.2 (C), 125.0 (C), 124.8 (m, $CF_3$), 122.5 (C), 122.3 (C), 72.0 (C), 71.5 (C), 52.8 ($CH_3$). **HRMS** (ESI) calcd for $C_{25}H_{14}F_6O_3$ [M+H] 493.0869, found 493.0864.

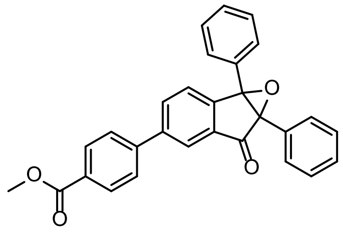

**Methyl 4-(6-oxo-1a,6a-diphenyl-1a,6a-dihydro-6H-indeno[1,2-b]oxiren-4-yl)benzoate (S10):** The title compound **S10** was synthesized from **S6** (20 mg, 48 µmol, 1 eq) according to **general method M2**. Purification by silica chromatography (0–10% EtOAc/cyclohexane, linear gradient) afforded **S10** as white solid (12 mg, 58%). $^{1}$**H NMR** ($CDCl_3$, 400 MHz) δ 8.19 – 8.12 (m, 3H), 7.84 (dd, *J* = 7.9, 1.8 Hz, 1H), 7.72 – 7.67 (m, 2H), 7.51 (d, *J* = 7.8 Hz, 1H), 7.45 (dd, *J* = 6.6, 3.1 Hz, 2H), 7.38 – 7.22 (m, 8H), 3.96 (s, 3H). $^{13}$**C NMR** ($CDCl_3$, 101 MHz) δ 196.3 (C), 166.9 (C), 148.5 (C), 144.0 (C), 142.2 (C), 135.9 (C), 133.5 (CH), 130.5 (CH), 130.0 (C), 129.8 (C), 129.0 (CH), 128.7 (CH), 128.5 (CH), 128.2 (CH),

128.1 (C), 128.1 (CH), 128.0 (CH), 127.4 (C), 127.3 (CH), 126.1 (CH), 124.3 (CH), 72.4 (C), 72.0 (C), 52.4 ($CH_3$). **HRMS** (ESI) calcd for $C_{29}H_{20}O_4$ [M+H] 433.1434, found 433.1429.

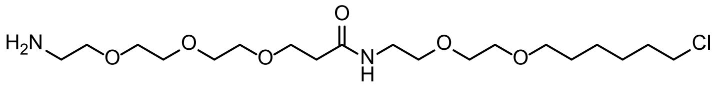

**3-(2-(2-(2-Aminoethoxy)ethoxy)ethoxy)-N-(2-(2-((6-chlorohexyl)oxy)ethoxy)ethyl)-propanamide (S9):** The procedure was adapted from Houštecká *et al.*[18] A reaction vial was charged with 2,2-dimethyl-4-oxo-3,8,11,14-tetraoxa-5-azaheptadecan-17-oic acid (500 mg, 1.56 mmol, 1 eq) and HBTU (590 mg, 1.56 mmol, 1 eq). The vial was evacuated and backfilled with argon (3x). Anhydrous DMF (10 mL) was added, followed by the dropwise addition of DIPEA (725 µL, 4.67 mmol, 3 eq). The reaction was stirred at room temperature for 30 min. A solution of 2-(2-((6-chlorohexyl)oxy)ethoxy)ethan-1-amine hydrochloride (403 mg, 1.56 mmol, 1 eq) in DMF (4 mL) was then added, and the solution was stirred overnight at room temperature. The reaction mixture was diluted with water, extracted with EtOAc (3x), and the combined organic layers were washed with brine (1x). After drying over $Na_2SO_4$ and removal of the solvent under reduced pressure, the crude material was purified by silica gel flash chromatography (0–10% MeOH/$CH_2Cl_2$, linear gradient). The resulting intermediate was dissolved in $CH_2Cl_2$ (5 mL), and TFA (1 mL) was carefully added dropwise. The reaction was stirred for 5 h and monitored by LCMS. Toluene (3 mL) was added, the reaction mixture was concentrated to dryness and then azeotroped with MeOH three times. The resulting yellowish oil **S9** (641 mg, 96% over 2 steps) was stored at -20°C and used without further purification. **$^1$H NMR** ($CDCl_3$, 400 MHz) δ 3.88 – 3.80 (m, 2H), 3.76 – 3.67 (m, 4H), 3.63 (dtd, $J$ = 10.0, 5.2, 2.1 Hz, 10H), 3.58 (dt, $J$ = 6.2, 1.8 Hz, 2H), 3.53 (t, $J$ = 6.6 Hz, 2H), 3.51 – 3.42 (m, 4H), 3.23 – 3.13 (m, 2H), 2.56 (t, $J$ = 5.4 Hz, 2H), 1.83 – 1.71 (m, 2H), 1.60 (p, $J$ = 7.0 Hz, 2H), 1.51 – 1.39 (m, 2H), 1.35 (dddd, $J$ = 14.3, 9.2, 6.4, 3.4 Hz, 2H). **$^{13}$C NMR** ($CDCl_3$, 101 MHz) δ 173.3 (C), 71.4 ($CH_2$), 70.3 ($CH_2$), 70.1 ($CH_2$), 70.0 ($CH_2$), 69.9 ($CH_2$), 69.9 ($CH_2$), 69.3 ($CH_2$), 67.1 ($CH_2$), 67.0 ($CH_2$), 45.1 ($CH_2$), 40.0 ($CH_2$), 39.4 ($CH_2$), 36.3 ($CH_2$), 32.6 ($CH_2$), 29.4 ($CH_2$), 26.8 ($CH_2$), 25.4 ($CH_2$). **HRMS** (ESI) calcd for $C_{19}H_{39}ClN_2O_6$ $[M+H]^+$ 427.2569, found 427.2567.

**Synthesis of DIO ligands**

**General procedure for the preparation of HaloTag- and SNAP-tag ligands *via* amide coupling (General Method M3):** The following procedure for **1** is shown as representative. **S5** (10 mg, 29 μmol, 1.0 eq), HaloTag ligand-amine hydrochloride[19] (15 mg, 58 μmol, 2.0 eq) and HATU (22 mg, 58 μmol, 2.0 eq) were dissolved in $CH_2Cl_2$ (1 mL), and DIPEA (20 μL, 0.12 mmol, 4.0 eq) was added. The reaction was stirred at room temperature for 2 h, and monitored by LCMS. After completion, the solvent was removed under reduced pressure. The crude material was purified by reverse phase HPLC (5–95% MeCN/$H_2O$, linear gradient) affording **1** as a white solid (7 mg, 56%).

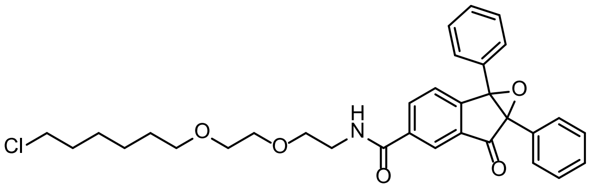

**1: $^1$H NMR** (($CD_3)_2SO$, 400 MHz) δ 8.85 (t, *J* = 5.5 Hz, 1H), 8.37 (d, *J* = 1.6 Hz, 1H), 8.20 (dd, *J* = 7.9, 1.7 Hz, 1H), 7.51 – 7.41 (m, 3H), 7.40 – 7.28 (m, 8H), 3.61 – 3.53 (m, 6H), 3.50 – 3.43 (m, 4H), 3.36 (t, *J* = 6.5 Hz, 2H), 1.73 – 1.61 (m, 2H), 1.46 (p, *J* = 6.8 Hz, 2H), 1.39 – 1.20 (m, 4H). **Analytical HPLC**: $t_R$ = 5.11 min, >99% purity (5–95% MeCN/$H_2O$, linear gradient, with constant 0.1% v/v formic acid additive, 9 min run, 0.6 mL·min$^{-1}$ flow, detection at 280 nm). **HRMS** (ESI) calcd for $C_{32}H_{34}ClNO_5$ [M+H]$^+$ 548.2198, found 548.2200.

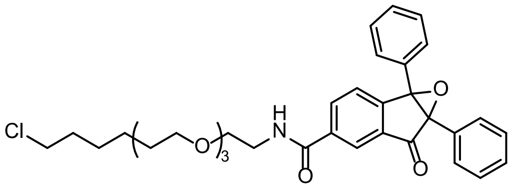

**2**: The title compound (**2**) (11 mg, 41%) was synthesized from **S5** (15 mg, 44 μmol, 1.0 eq) and ethanamine,2-[2-[2-[(6-chlorohexyl)oxy]ethoxy]ethoxy] TFA salt[17] (17 mg, 43 μmol, 1.0 eq) according to **general method M3.** **$^1$H NMR** (($CD_3)_2SO$, 400 MHz) δ 8.85 (s, 1H), 8.37 (d, *J* = 1.6 Hz, 1H), 8.21 (dd, *J* = 7.8, 1.7 Hz, 1H), 7.51 – 7.41 (m, 3H), 7.41 – 7.28 (m, 8H), 3.64 – 3.40 (m, 14H), 3.35 (d, *J* = 6.5 Hz, 2H), 1.74 – 1.62 (m, 2H), 1.47 (q, *J* = 7.0 Hz, 3H), 1.40 – 1.21 (m, 4H). **Analytical HPLC**: $t_R$ = 5.11 min, >99% purity (5–95% MeCN/$H_2O$, linear gradient, with constant 0.1% v/v formic acid additive, 9 min run, 0.6 mL·min$^{-1}$ flow, detection at 280 nm). **HRMS** (ESI) calcd for $C_{34}H_{38}ClNO_6$ [M+H]$^+$ 592.2460, found 592.2457.

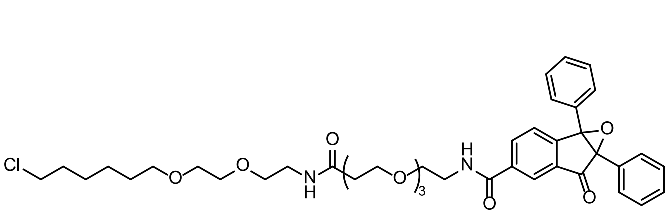

**3**: The title compound (**3**) (16 mg, 49%) was synthesized from **S5** (15 mg, 44 μmol, 1.0 eq) and **S9** (47 mg, 87 μmol, 2.0 eq) according to **general method M3**. **$^1$H NMR** (($CD_3)_3SO$, 400 MHz) δ 8.86 (t, *J* = 5.5 Hz, 1H), 8.37 (d, *J* = 1.6 Hz, 1H), 8.21 (dd, *J* = 7.9, 1.7 Hz, 1H), 7.88 (t, *J* = 5.7 Hz, 1H), 7.51 – 7.42 (m, 3H), 7.42 – 7.25 (m, 8H), 3.66 – 3.41 (m, 20H), 3.38 (dd, *J* = 11.6, 6.0 Hz, 4H), 3.18 (q, *J* = 5.8 Hz, 2H), 2.30 (t, *J* = 6.5 Hz, 2H), 1.75 – 1.63 (m, 2H), 1.47 (p, *J* = 6.8 Hz, 2H), 1.41 – 1.21 (m, 4H). **Analytical HPLC**: $t_R$ = 4.84 min, >99% purity (5–95% MeCN/$H_2O$, linear gradient, with constant 0.1%

v/v formic acid additive, 9 min run, 0.6 mL·min$^{-1}$ flow, detection at 254 nm). **HRMS** (ESI) calcd for $C_{41}H_{51}ClN_2O_9$ $[M+H]^+$ 751.3356, found 751.3346.

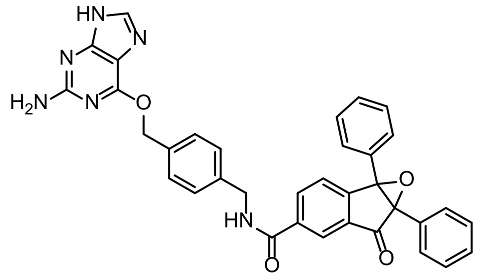

**4**: The title compound **4** (4 mg, 23%) was synthesized from **S5** (15 mg, 34 µmol, 1 eq) and 6-((4-(aminomethyl)benzyl)oxy)-9H-purin-2-amine (12 mg, 44 µmol, 1.5 eq) according to **general method M3.** **$^1$H NMR** ($(CD_3)_2SO$, 400 MHz) δ 9.36 (t, *J* = 5.8 Hz, 1H), 8.41 (d, *J* = 1.7 Hz, 1H), 8.24 (dd, *J* = 7.9, 1.7 Hz, 1H), 7.82 (s, 1H), 7.52 – 7.43 (m, 4H), 7.41 – 7.27 (m, 11H), 6.27 (s, 2H), 5.46 (s, 2H), 4.52 (d, *J* = 5.8 Hz, 2H). **Analytical HPLC**: $t_R$ = 4.24 min, >99% purity (5–95% MeCN/$H_2O$, linear gradient, with constant 0.1% v/v formic acid additive, 9 min run, 0.6 mL·min$^{-1}$ flow, detection at 280 nm). **HRMS** (ESI) calcd for $C_{35}H_{26}N_6O_4$ $[M+H]^+$ 595.2088, found 595.2086.

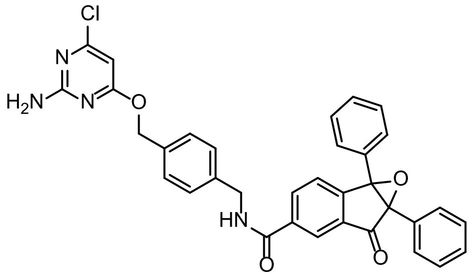

**5:** The title compound **5** (9 mg, 52%) was synthesized from **S5** (10 mg, 29 µmol, 1.0 eq) and 4-((4-(aminomethyl)benzyl)oxy)-6-chloropyrimidin-2-amine (12 mg, 44 µmol, 1.5 eq) according to **general method M3.** **$^1$H NMR** ($CDCl_3$, 400 MHz) δ 9.37 (t, *J* = 5.9 Hz, 1H), 8.41 (d, *J* = 1.7 Hz, 1H), 8.24 (dd, *J* = 7.9, 1.7 Hz, 1H), 7.52 – 7.25 (m, 15H), 7.10 (s, 2H), 6.13 (s, 1H), 5.30 (s, 2H), 4.51 (d, *J* = 5.8 Hz, 2H). **Analytical HPLC**: $t_R$ = 4.95 min, >99% purity (5–95% MeCN/$H_2O$, linear gradient, with constant 0.1% v/v formic acid additive, 9 min run, 0.6 mL·min$^{-1}$ flow, detection at 280 nm). **HRMS** (ESI) calcd for $C_{34}H_{25}ClN_4O_4$ $[M+H]^+$ 589.1637, found 589.1625.

**General procedure for preparation of HaloTag-ligands *via* amide coupling following removal of the methyl ester (General Method M4):** The following procedure for **6** is shown as representative. **S6** (15 mg, 33 µmol, 1 eq) was dissolved in a mixture of THF:$H_2O$:MeOH (1:1:1, 1.5 mL). LiOH·$H_2O$ (4 mg, 99 µmol, 3.0 eq) was added and the reaction was stirred at room temperature for 1 h. The solvent was removed under reduced pressure, diluted with $H_2O$ and the aqueous phase was extracted with EtOAc (3x). The solvent was removed under reduced pressure. The resulting crude material (15 mg, quant.), 2-(2-(6-chlorohexyloxy)ethoxy)ethanamine hydrochloride (17 mg, 66 µmol, 2.0 eq) and HATU (25 mg, 66 µmol, 2.0 eq) were dissolved in $CH_2Cl_2$ (1 mL) and DIPEA (23 µL, 0.13 mmol, 4.0 eq) was added. The reaction was stirred at room temperature for 2 h, and monitored by LCMS. After completion, the solvent was removed under reduced pressure. The crude was purified by reverse phase HPLC (5–95% MeCN/$H_2O$, linear gradient), affording the product **6** as a white solid (9 mg, 64%).

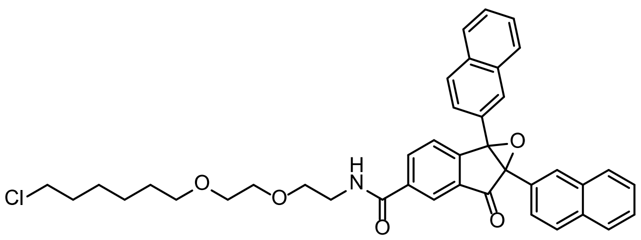

**6:** **$^{1}$H NMR** ($(CD_3)_2SO$, 400 MHz) δ 8.88 (t, *J* = 5.5 Hz, 1H), 8.43 (d, *J* = 1.7 Hz, 1H), 8.24 (dd, *J* = 7.8, 1.7 Hz, 1H), 8.15 (d, *J* = 1.7 Hz, 1H), 8.04 (d, *J* = 1.7 Hz, 1H), 7.96 – 7.91 (m, 1H), 7.90 – 7.78 (m, 5H), 7.63 – 7.52 (m, 3H), 7.53 – 7.43 (m, 4H), 3.63 – 3.52 (m, 6H), 3.49 (dt, *J* = 5.7, 2.7 Hz, 3H), 3.37 (t, *J* = 6.5 Hz, 2H), 1.72 – 1.63 (m, 2H), 1.49 – 1.43 (m, 3H), 1.39 – 1.28 (m, 4H). **Analytical HPLC**: $t_R$ = 5.59 min, >99% purity (5–95% MeCN/$H_2O$, linear gradient, with constant 0.1% v/v formic acid additive, 9 min run, 0.6 mL·min$^{-1}$ flow, detection at 280 nm). **HRMS** (ESI) calcd for $C_{40}H_{38}ClNO_5$ $[M+H]^+$ 648.2511, found 648.2504.

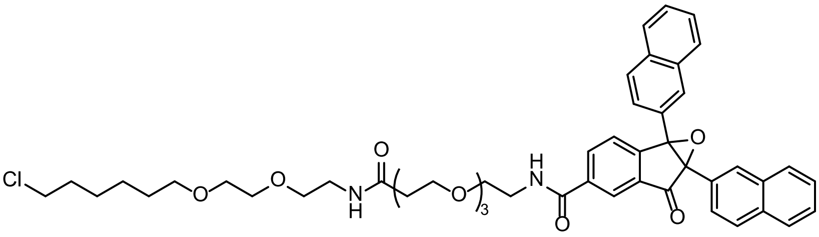

**7:** The title compound **7** (6 mg, 21%) was synthesized from **S6** (15 mg, 44 µmol, 1.0 eq) and **S9** (35 mg, 65 µmol, 2.0 eq) according to **general method M4**. **$^{1}$H NMR** ($(CD_3)_2SO$, 400 MHz) δ 8.89 (t, *J* = 5.5 Hz, 1H), 8.43 (d, *J* = 1.6 Hz, 1H), 8.24 (dd, *J* = 7.8, 1.7 Hz, 1H), 8.15 (d, *J* = 1.7 Hz, 1H), 8.04 (d, *J* = 1.7 Hz, 1H), 7.97 – 7.76 (m, 6H), 7.62 – 7.54 (m, 3H), 7.53 – 7.45 (m, 4H), 3.64 – 3.41 (m, 20H), 3.43 – 3.31 (m, 4H), 3.18 (q, *J* = 5.8 Hz, 2H), 2.30 (t, *J* = 6.5 Hz, 2H), 1.74 – 1.62 (m, 2H), 1.45 (q, *J* = 6.9 Hz, 2H), 1.40 – 1.25 (m, 4H). **Analytical HPLC**: $t_R$ = 5.32 min, >99% purity (5–95% MeCN/$H_2O$, linear gradient, with constant 0.1% v/v formic acid additive, 9 min run, 0.6 mL·min$^{-1}$ flow, detection at 280 nm). **HRMS** (ESI) calcd for $C_{49}H_{55}ClN_2O_9$ $[M+H]^+$ 851.3669, found 851.3652.

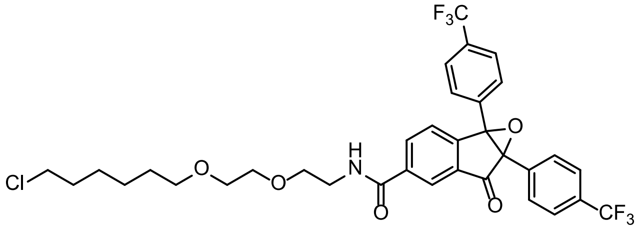

**8:** The title compound **8** (13 mg, 63%) was synthesized from **S7** (15 mg, 30 µmol, 1.0 eq) and HTL-amine hydrochloride[19] (16 mg, 61 µmol, 2.0 eq) according to **general method M4**. **$^{1}$H NMR** ($(CD_3)_2SO$, 400 MHz) δ 8.88 (t, *J* = 5.5 Hz, 1H), 8.41 (d, *J* = 1.8 Hz, 1H), 8.22 (dd, *J* = 7.9, 1.7 Hz, 1H), 7.81 – 7.65 (m, 8H), 7.53 (d, *J* = 7.8 Hz, 1H), 3.62 – 3.53 (m, 6H), 3.50 – 3.42 (m, 4H), 3.36 (t, *J* = 6.6 Hz, 2H), 1.73 – 1.62 (m, 2H), 1.46 (p, *J* = 6.8 Hz, 3H), 1.39 – 1.27 (m, 4H). **Analytical HPLC**: $t_R$ = 5.48 min, >99% purity (5–95% MeCN/$H_2O$, linear gradient, with constant 0.1% v/v formic acid additive, 9 min run, 0.6 mL·min$^{-1}$ flow, detection at 280 nm). **HRMS** (ESI) calcd for $C_{34}H_{32}ClF_6NO_5$ $[M+H]^+$ 684.1946, found 684.1931.

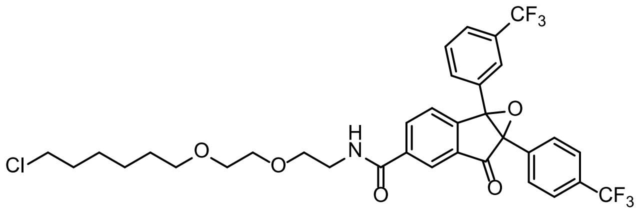

**9:** The title compound **9** (10 mg, 69%) was synthesized from **S9** (15 mg, 30 μmol, 1 eq) according to **general method M4**. Purification by silica chromatography (0–10% EtOAc/cyclohexane, linear gradient) afforded **9** as white solid. **$^1$H NMR** (($(CH_3)_3SO$, 400 MHz) δ 8.87 (t, J = 5.6 Hz, 1H), 8.40 (d, J = 1.7 Hz, 1H), 8.22 (dd, J = 7.8, 1.7 Hz, 1H), 7.81 (t, J = 7.2 Hz, 3H), 7.78 – 7.70 (m, 2H), 7.71 – 7.51 (m, 4H), 3.63 – 3.41 (m, 10H), 3.36 (s, 2H), 1.67 (p, J = 6.8 Hz, 2H), 1.46 (p, J = 6.8 Hz, 2H), 1.31 (dt, J = 33.4, 7.8 Hz, 4H). **Analytical HPLC**: $t_R$ = 5.48 min, >99% purity (5–95% MeCN/$H_2O$, linear gradient, with constant 0.1% v/v formic acid additive, 9 min run, 0.6 mL·min$^{-1}$ flow, detection at 280 nm). **HRMS** (ESI) calcd for $C_{34}H_{32}ClF_6NO_5$ $[M+H]^+$ 684.1946, found 684.1931.

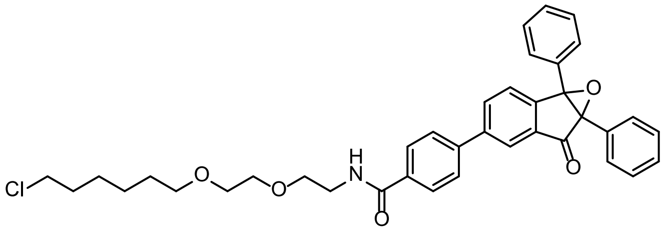

**10:** The title compound **10** (6 mg, 40%) was synthesized from **S10** (10 mg, 24 μmol, 1 eq) according to **general method M4**. Purification by silica chromatography (0–10% EtOAc/cyclohexane, linear gradient) afforded **10** as white solid. **$^1$H NMR** ($(CD_3)_2SO$, 400 MHz) δ 8.62 (t, *J* = 5.6 Hz, 1H), 8.19 (d, *J* = 1.8 Hz, 1H), 8.09 (dd, *J* = 7.9, 1.8 Hz, 1H), 8.02 – 7.97 (m, 2H), 7.92 – 7.85 (m, 2H), 7.49 (dd, *J* = 7.5, 2.2 Hz, 3H), 7.42 – 7.28 (m, 8H), 3.62 – 3.51 (m, 6H), 3.52 – 3.40 (m, 4H), 3.40 – 3.28 (m, 2H), 1.72 – 1.61 (m, 2H), 1.46 (p, *J* = 6.8 Hz, 2H), 1.39 – 1.20 (m, 4H). **Analytical HPLC**: $t_R$ = 5.42 min, >99% purity (5–95% MeCN/$H_2O$, linear gradient, with constant 0.1% v/v formic acid additive, 9 min run, 0.6 mL·min$^{-1}$ flow, detection at 280 nm). **HRMS** (ESI) calcd for $C_{38}H_{38}ClNO_5$ $[M+H]^+$ 624.2511, found 624.2501.

#### Synthesis of clickable fluorophore

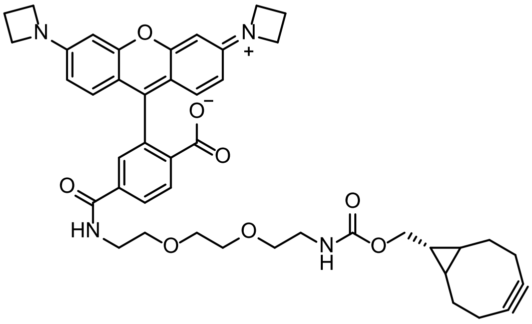

**JF$_{549}$-BCN (11):** A vial was charged with JF$_{549}$-NHS ester (20 mg, 36 μmol, 1.0 eq) and BCN-amine (12 mg, 36 μmol, 1.0 eq). DMF (1 mL) was added and the reaction was stirred at room temperature for 3 h. The reaction was quenched with $NH_4Cl$, extracted with EtOAc (3x), and the combined organic layers were washed with $NH_4Cl$ (1x) and brine (1x). Purification by reverse phase HPLC (10–80% MeCN/$H_2O$, linear gradient, with constant 0.1% v/v TFA additive) afforded **11** as a pink solid (20 mg, 63%, TFA salt). **$^1$H NMR** ($CD_3OD$, 400 MHz) δ 8.14 (d, *J* = 8.1 Hz, 1H), 8.09 (dd, *J* = 8.1, 1.8 Hz, 1H), 7.70 (d, *J* = 1.8

Hz, 1H), 7.16 (d, *J* = 9.2 Hz, 2H), 6.57 (dd, *J* = 9.2, 2.2 Hz, 2H), 6.48 (d, *J* = 2.2 Hz, 2H), 4.27 (t, *J* = 7.6 Hz, 8H), 4.06 (d, *J* = 8.1 Hz, 2H), 3.71 – 3.53 (m, 10H), 3.47 (t, *J* = 5.5 Hz, 2H), 3.17 (t, *J* = 5.5 Hz, 2H), 2.66 (s, 2H), 2.61 – 2.47 (m, 4H), 2.18 (td, *J* = 22.3, 12.4 Hz, 1H), 1.53 (d, *J* = 10.1 Hz, 2H), 1.31 (d, *J* = 14.8 Hz, 4H), 0.94 – 0.84 (m, 1H). **$^{13}$C NMR** ($CD_3OD$, 101 MHz) δ 172.1 (C), 168.7 (C), 161.6 (C), 158.8 (C), 158.0 (C), 143.9 (C), 136.5 (C), 134.3 (C), 132.9 (CH), 131.1 (CH), 129.6 (CH), 129.4 (CH), 115.0 (C), 113.1 (CH), 99.5 (C), 95.1 (CH), 73.7 (C), 71.6 ($CH_2$), 71.3 ($CH_2$), 71.3 ($CH_2$), 71.1 ($CH_2$), 70.5 ($CH_2$), 63.6 ($CH_2$), 52.8 ($CH_2$), 49.5 (C), 41.6 ($CH_2$), 41.0 ($CH_2$), 40.4 (C), 30.8 (C), 30.2 ($CH_2$), 21.9 ($CH_2$), 21.4 (CH), 19.0 (CH), 16.9 ($CH_2$). **HRMS** (ESI) calcd for $C_{44}H_{48}N_4O_8$ $[M+H]^+$ 761.3545, found 761.3531.

### NMR Spectra and HPLC Traces

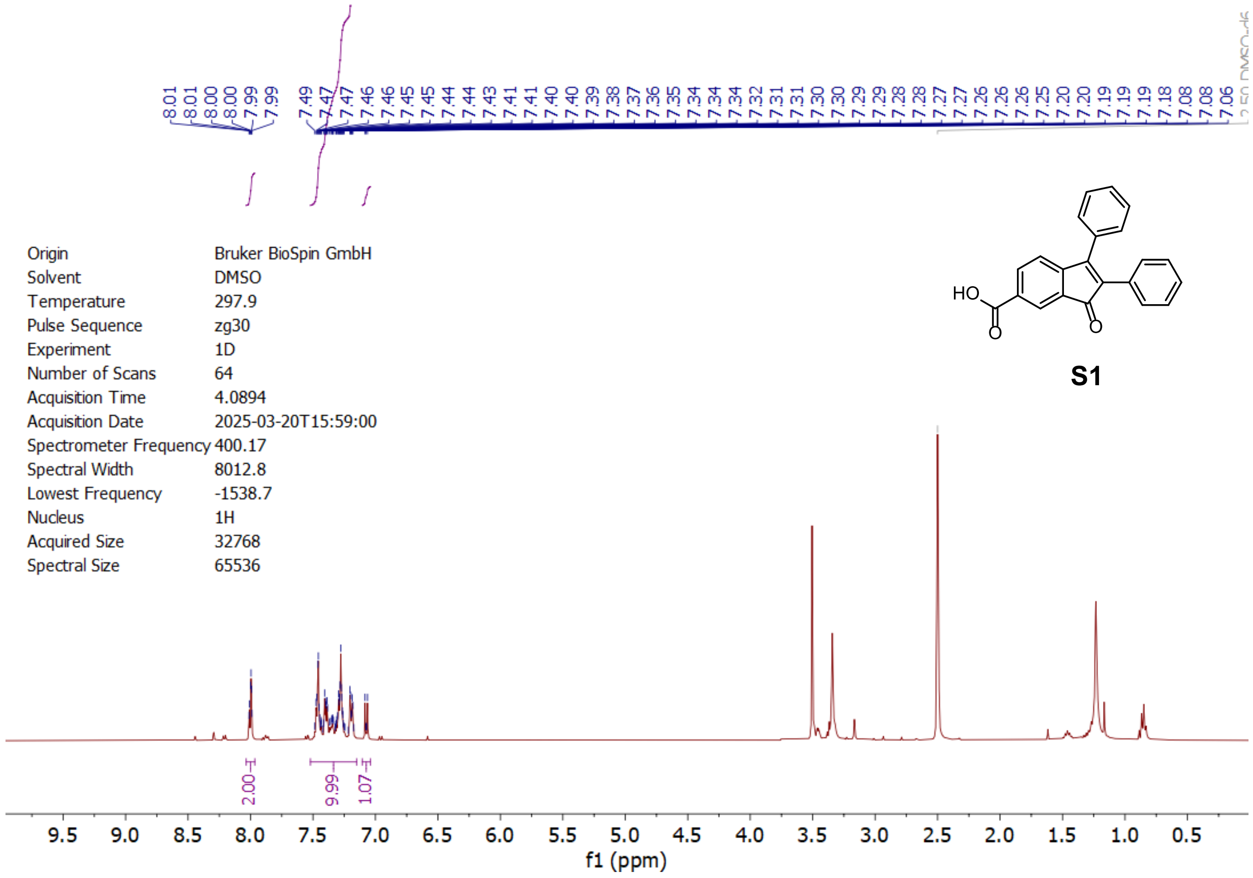

**S1**

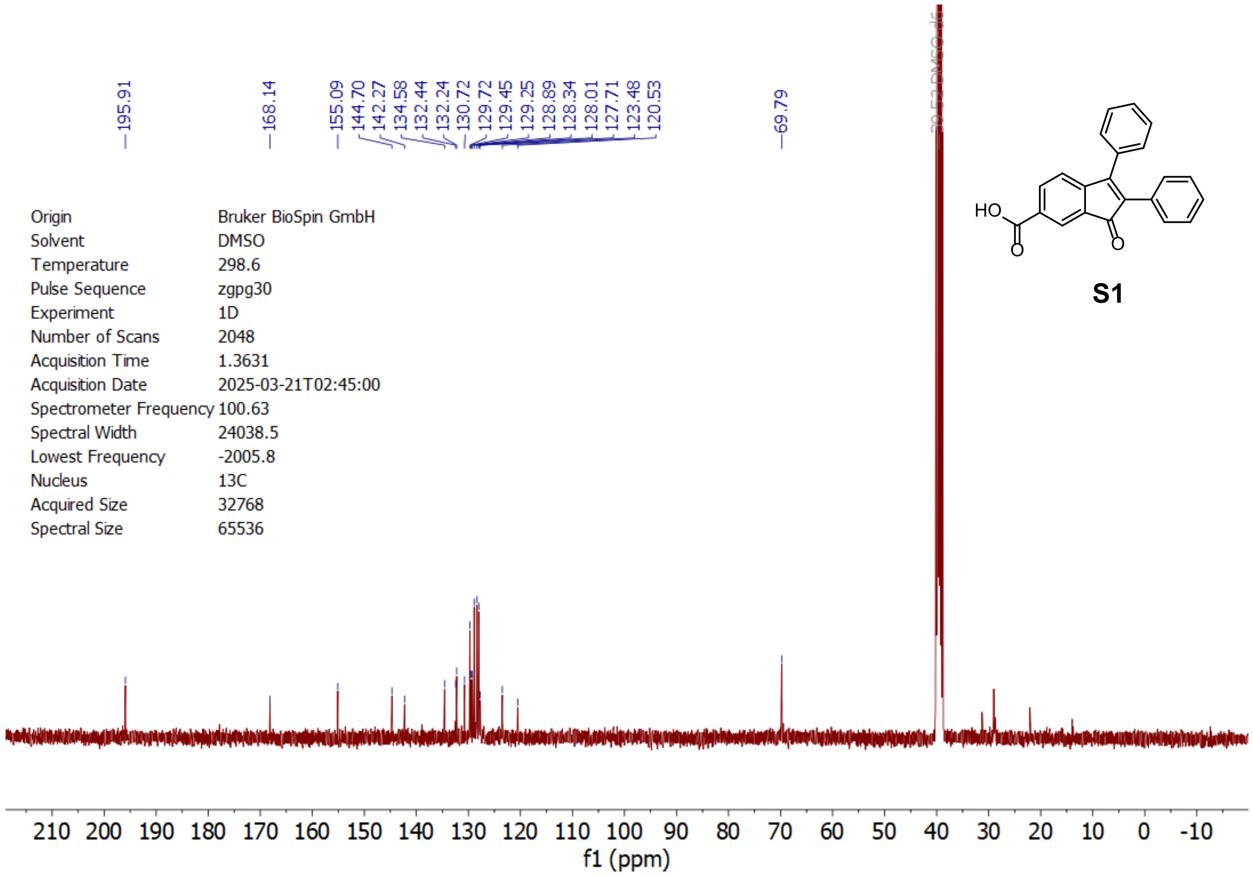

**S1**

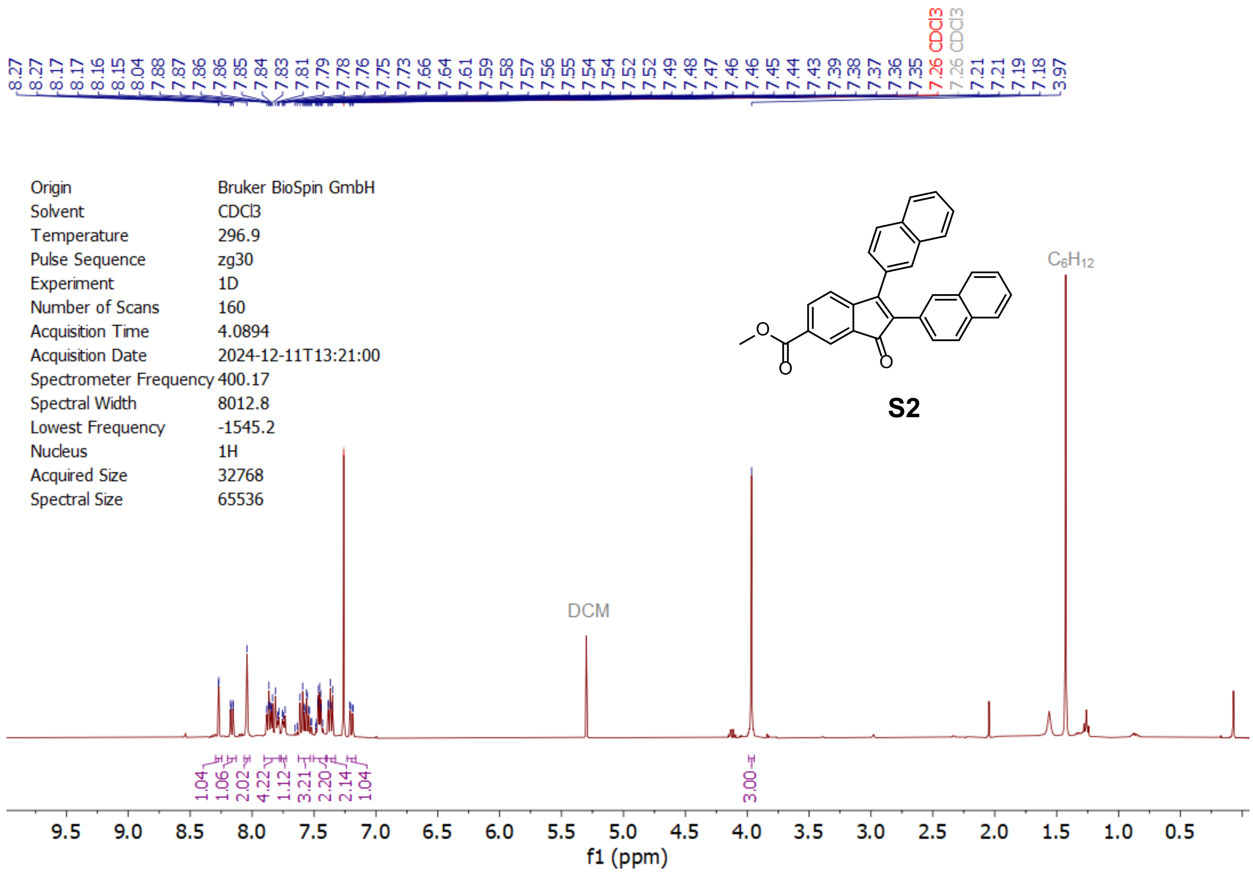
8.27 8.27 8.17 8.17 8.16 8.15 8.04 7.88 7.87 7.86 7.86 7.85 7.84 7.83 7.81 7.79 7.78 7.76 7.75 7.73 7.66 7.64 7.61 7.59 7.58 7.57 7.56 7.55 7.54 7.54 7.52 7.52 7.49 7.48 7.47 7.46 7.46 7.45 7.44 7.43 7.39 7.38 7.37 7.36 7.35 7.26 CDCl3 7.26 CDCl3 7.21 7.21 7.19 7.18 3.97
Origin Bruker BioSpin GmbH
Solvent CDCl3
Temperature 296.9
Pulse Sequence zg30
Experiment 1D
Number of Scans 160
Acquisition Time 4.0894
Acquisition Date 2024-12-11T13:21:00
Spectrometer Frequency 400.17
Spectral Width 8012.8
Lowest Frequency -1545.2
Nucleus 1H
Acquired Size 32768
Spectral Size 65536
S2
C6H12
DCM
1.04 1.06 2.02 4.22 1.12 3.21 2.20 2.14 1.04 3.00
9.5 9.0 8.5 8.0 7.5 7.0 6.5 6.0 5.5 5.0 4.5 4.0 3.5 3.0 2.5 2.0 1.5 1.0 0.5
f1 (ppm)

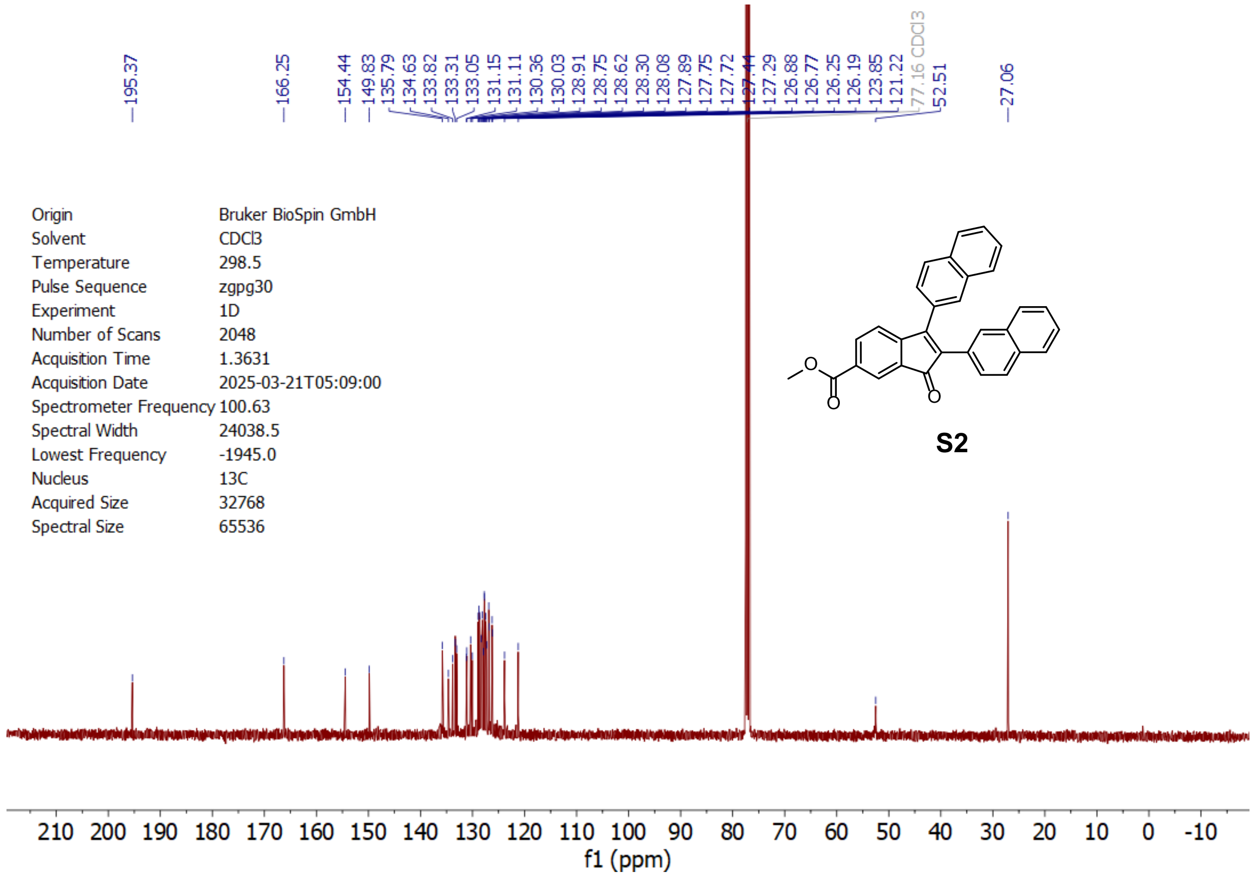
195.37 166.25 154.44 149.83 135.79 134.63 133.82 133.31 133.05 131.15 131.11 130.36 130.03 128.91 128.75 128.62 128.30 128.08 127.89 127.75 127.72 127.44 127.29 126.88 126.77 126.25 126.19 123.85 121.22 77.16 CDCl3 52.51 27.06
Origin Bruker BioSpin GmbH
Solvent CDCl3
Temperature 298.5
Pulse Sequence zgpg30
Experiment 1D
Number of Scans 2048
Acquisition Time 1.3631
Acquisition Date 2025-03-21T05:09:00
Spectrometer Frequency 100.63
Spectral Width 24038.5
Lowest Frequency -1945.0
Nucleus 13C
Acquired Size 32768
Spectral Size 65536
S2
210 200 190 180 170 160 150 140 130 120 110 100 90 80 70 60 50 40 30 20 10 0 -10
f1 (ppm)

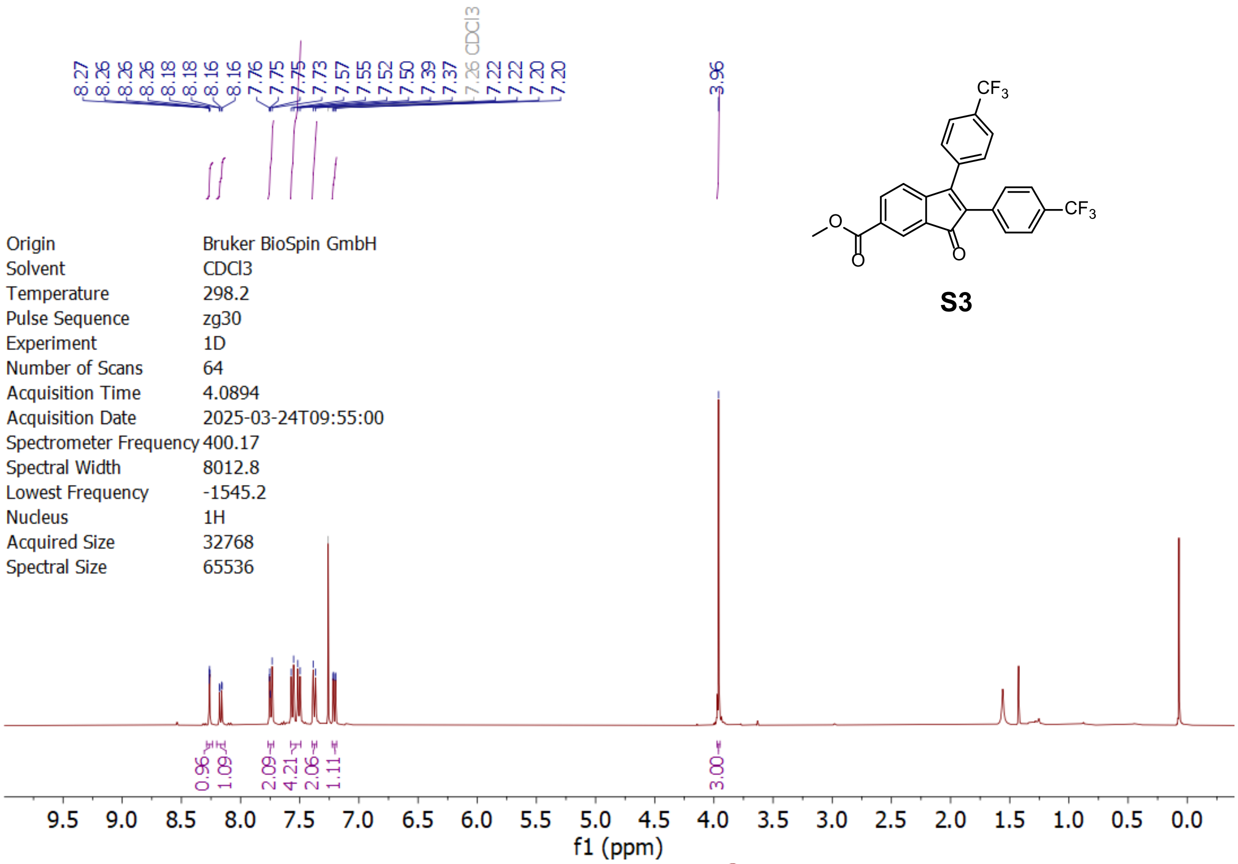
8.27 8.26 8.26 8.26 8.18 8.18 8.16 8.16 7.76 7.75 7.75 7.73 7.57 7.55 7.52 7.50 7.39 7.37 7.26 CDCl3 7.22 7.22 7.20 7.20
3.96
CF3
CF3
O
O
O
S3
Origin Bruker BioSpin GmbH
Solvent CDCl3
Temperature 298.2
Pulse Sequence zg30
Experiment 1D
Number of Scans 64
Acquisition Time 4.0894
Acquisition Date 2025-03-24T09:55:00
Spectrometer Frequency 400.17
Spectral Width 8012.8
Lowest Frequency -1545.2
Nucleus 1H
Acquired Size 32768
Spectral Size 65536
0.96 1.09 2.09 4.21 2.06 1.11
3.00
9.5 9.0 8.5 8.0 7.5 7.0 6.5 6.0 5.5 5.0 4.5 4.0 3.5 3.0 2.5 2.0 1.5 1.0 0.5 0.0
f1 (ppm)

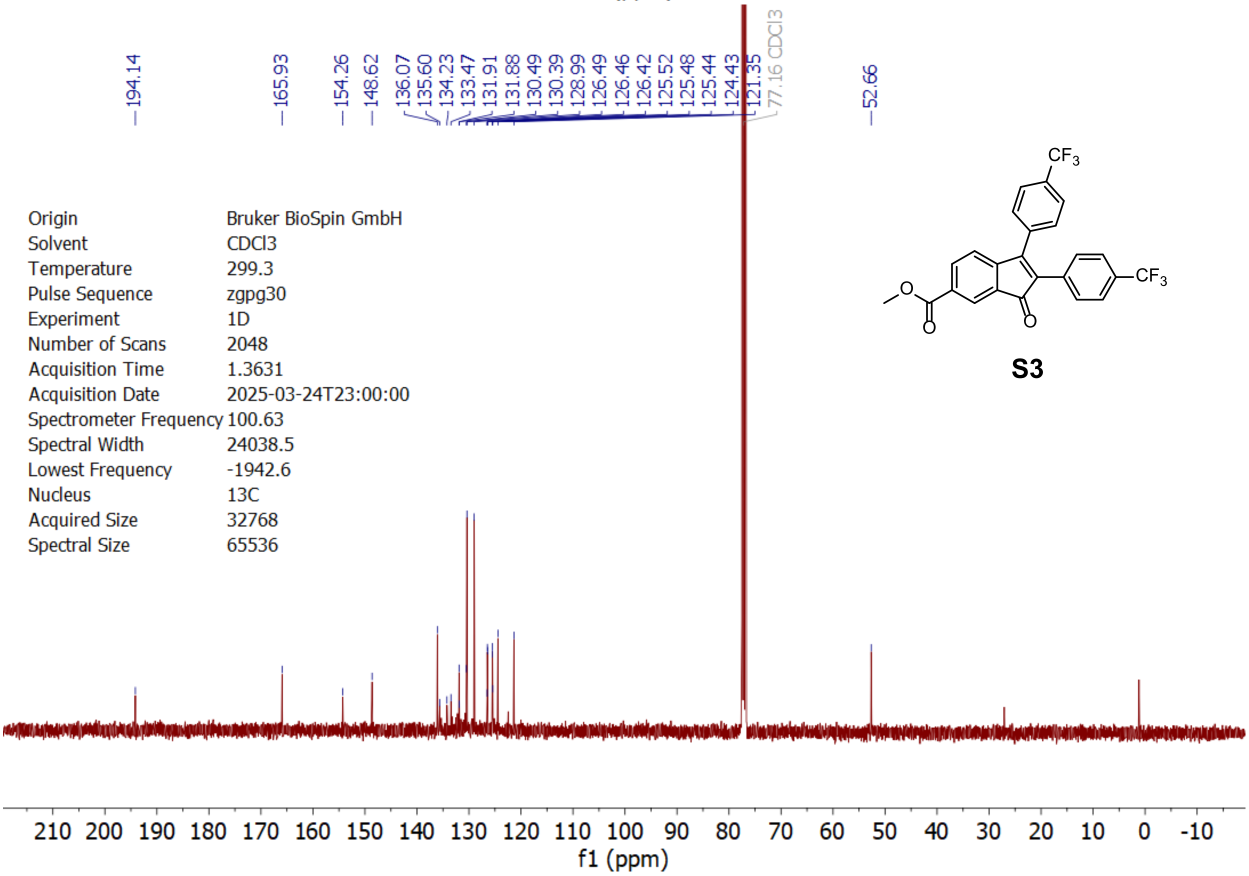
194.14
165.93
154.26
148.62
136.07 135.60 134.23 133.47 131.91 131.88 130.49 130.39 128.99 126.49 126.46 126.42 125.52 125.48 125.44 124.43 121.35
77.16 CDCl3
52.66
CF3
CF3
O
O
O
S3
Origin Bruker BioSpin GmbH
Solvent CDCl3
Temperature 299.3
Pulse Sequence zgpg30
Experiment 1D
Number of Scans 2048
Acquisition Time 1.3631
Acquisition Date 2025-03-24T23:00:00
Spectrometer Frequency 100.63
Spectral Width 24038.5
Lowest Frequency -1942.6
Nucleus 13C
Acquired Size 32768
Spectral Size 65536
210 200 190 180 170 160 150 140 130 120 110 100 90 80 70 60 50 40 30 20 10 0 -10
f1 (ppm)

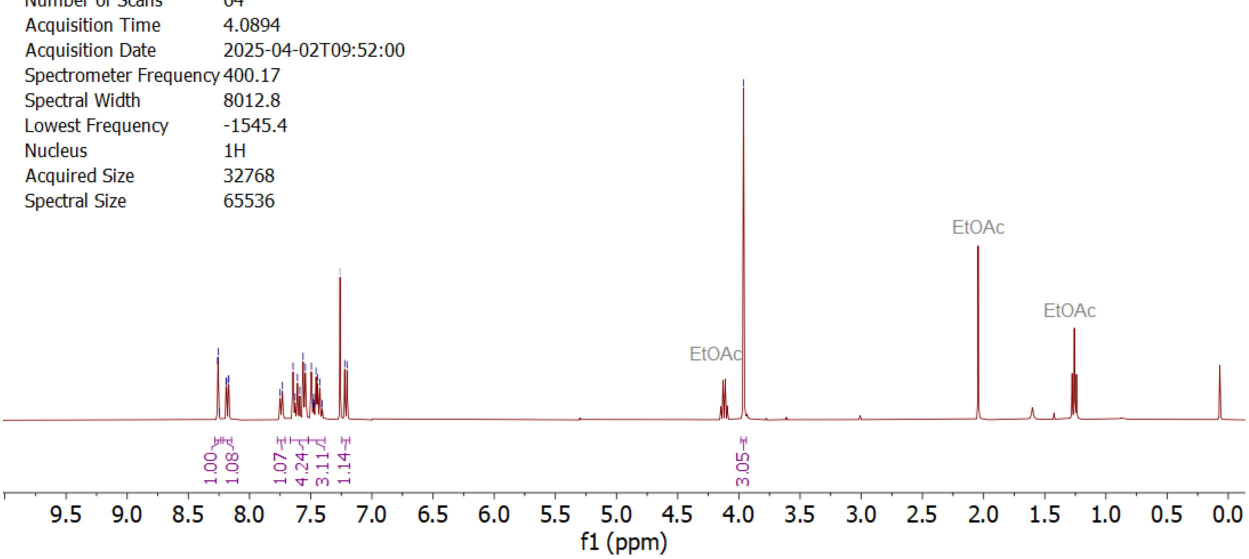
8.26
8.26
8.25
8.19
8.19
8.17
8.17
7.75
7.73
7.64
7.63
7.61
7.59
7.56
7.54
7.49
7.48
7.48
7.47
7.46
7.44
7.43
7.41
7.26 CDCl3
7.22
7.20
3.96
Origin Bruker BioSpin GmbH
Solvent CDCl3
Temperature 297.0
Pulse Sequence zg30
Experiment 1D
Number of Scans 64
Acquisition Time 4.0894
Acquisition Date 2025-04-02T09:52:00
Spectrometer Frequency 400.17
Spectral Width 8012.8
Lowest Frequency -1545.4
Nucleus 1H
Acquired Size 32768
Spectral Size 65536
EtOAc
EtOAc
EtOAc
1.00
1.08
1.07
4.24
3.11
1.14
3.05
9.5 9.0 8.5 8.0 7.5 7.0 6.5 6.0 5.5 5.0 4.5 4.0 3.5 3.0 2.5 2.0 1.5 1.0 0.5 0.0
f1 (ppm)

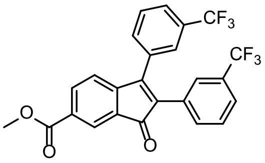
CF3
CF3

**S4**

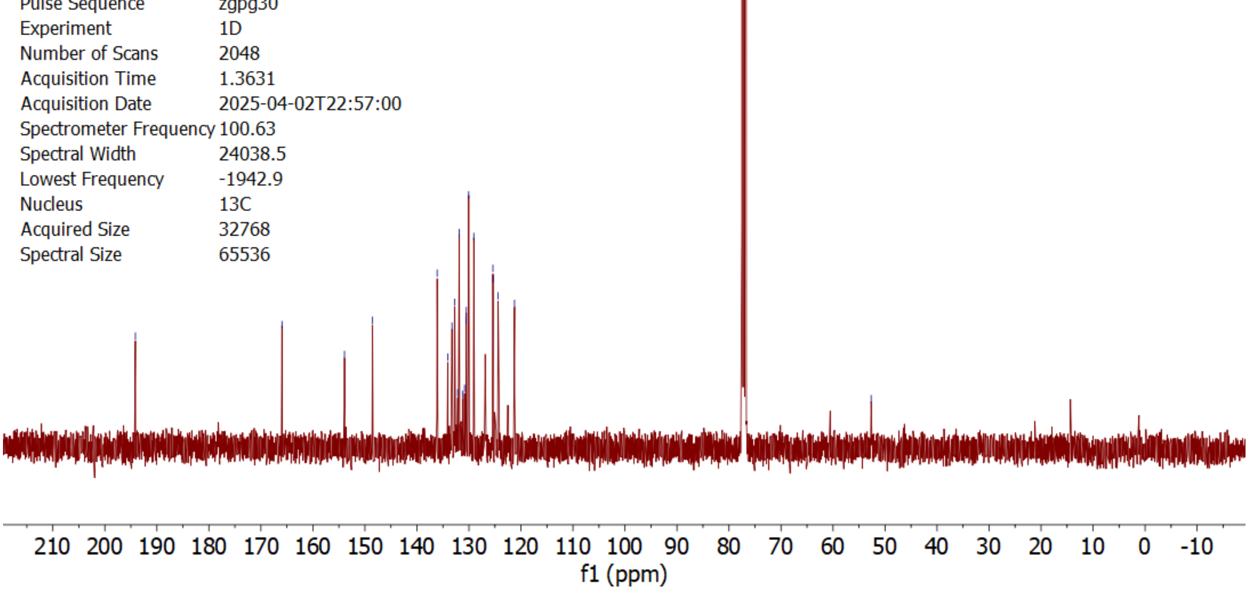
194.13
165.94
153.93
148.54
136.11
134.08
133.25
132.72
132.16
131.87
131.17
130.85
130.58
130.50
130.05
129.06
125.39
125.35
124.39
121.23
77.16 CDCl3
52.67
Origin Bruker BioSpin GmbH
Solvent CDCl3
Temperature 298.6
Pulse Sequence zgpg30
Experiment 1D
Number of Scans 2048
Acquisition Time 1.3631
Acquisition Date 2025-04-02T22:57:00
Spectrometer Frequency 100.63
Spectral Width 24038.5
Lowest Frequency -1942.9
Nucleus 13C
Acquired Size 32768
Spectral Size 65536
210 200 190 180 170 160 150 140 130 120 110 100 90 80 70 60 50 40 30 20 10 0 -10
f1 (ppm)

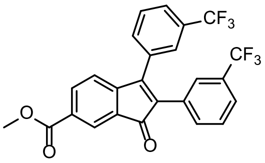
CF3
CF3

**S4**

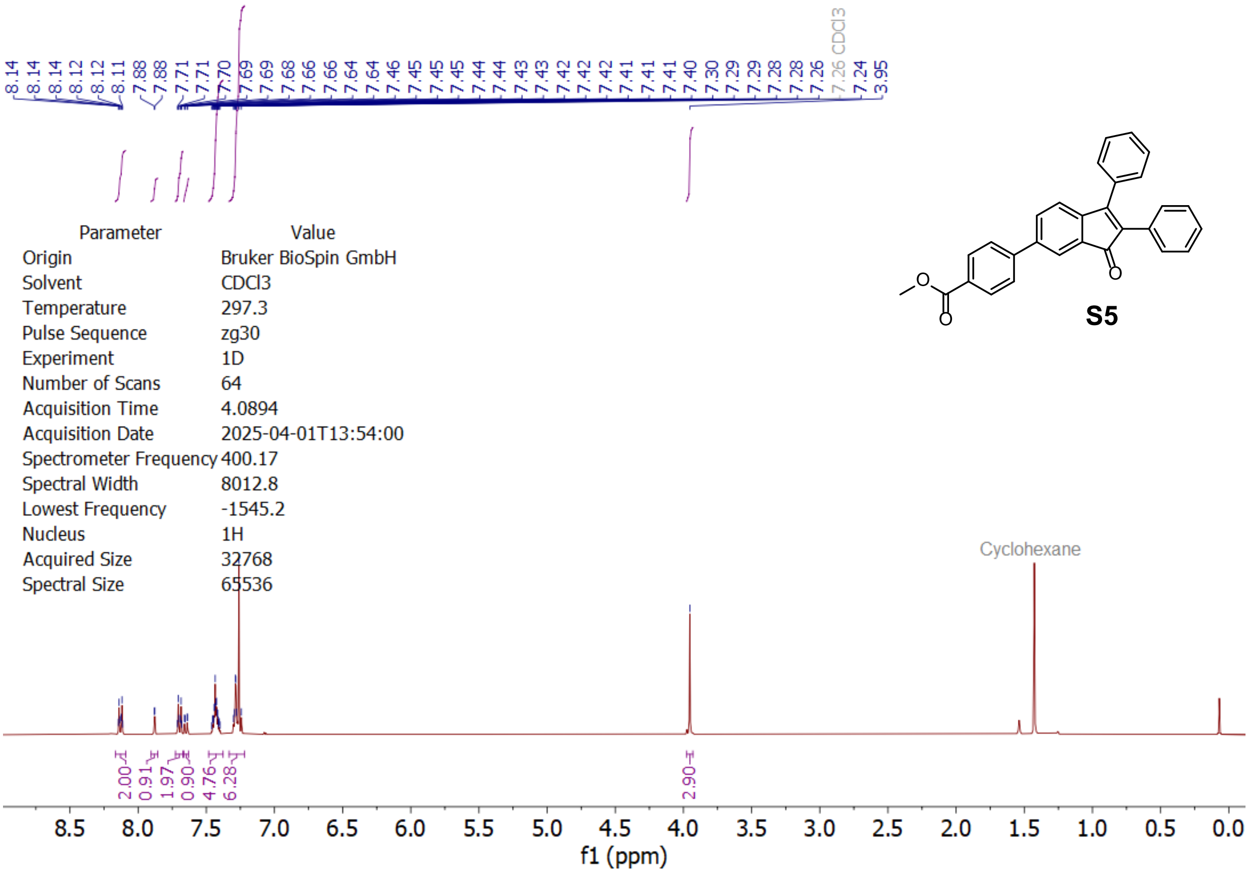

| Parameter | Value |
|---|---|
| Origin | Bruker BioSpin GmbH |
| Solvent | CDCl3 |
| Temperature | 297.3 |
| Pulse Sequence | zg30 |
| Experiment | 1D |
| Number of Scans | 64 |
| Acquisition Time | 4.0894 |
| Acquisition Date | 2025-04-01T13:54:00 |
| Spectrometer Frequency | 400.17 |
| Spectral Width | 8012.8 |
| Lowest Frequency | -1545.2 |
| Nucleus | 1H |
| Acquired Size | 32768 |
| Spectral Size | 65536 |

**S5**

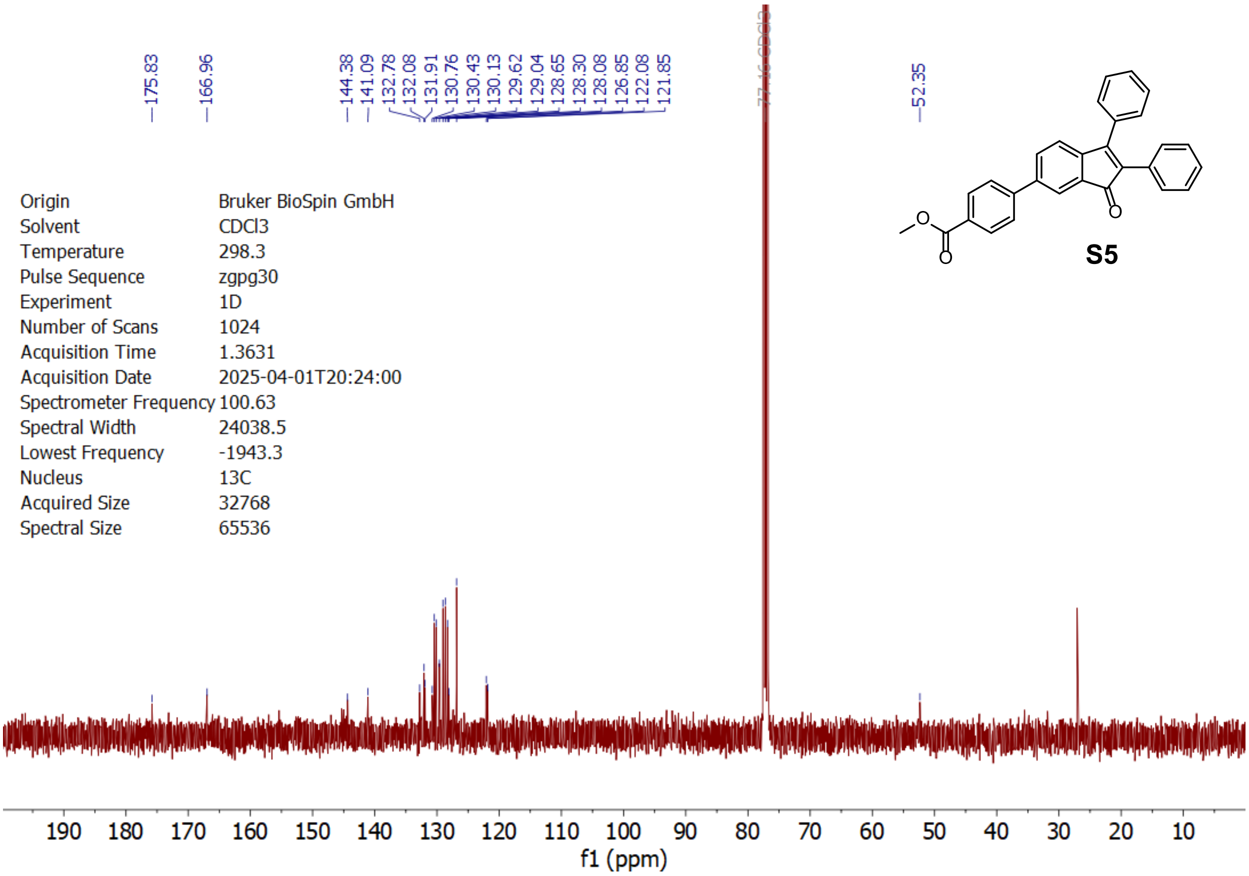

| | |
|---|---|
| Origin | Bruker BioSpin GmbH |
| Solvent | CDCl3 |
| Temperature | 298.3 |
| Pulse Sequence | zgpg30 |
| Experiment | 1D |
| Number of Scans | 1024 |
| Acquisition Time | 1.3631 |
| Acquisition Date | 2025-04-01T20:24:00 |
| Spectrometer Frequency | 100.63 |
| Spectral Width | 24038.5 |
| Lowest Frequency | -1943.3 |
| Nucleus | 13C |
| Acquired Size | 32768 |
| Spectral Size | 65536 |

**S5**

| | |
|---|---|
| Origin | Bruker BioSpin GmbH |
| Solvent | CDCl3 |
| Temperature | 295.3 |
| Pulse Sequence | zg30 |
| Experiment | 1D |
| Number of Scans | 160 |
| Acquisition Time | 4.0894 |
| Acquisition Date | 2024-01-05T15:57:00 |
| Spectrometer Frequency | 400.17 |
| Spectral Width | 8012.8 |
| Lowest Frequency | -1571.2 |
| Nucleus | 1H |
| Acquired Size | 32768 |
| Spectral Size | 65536 |

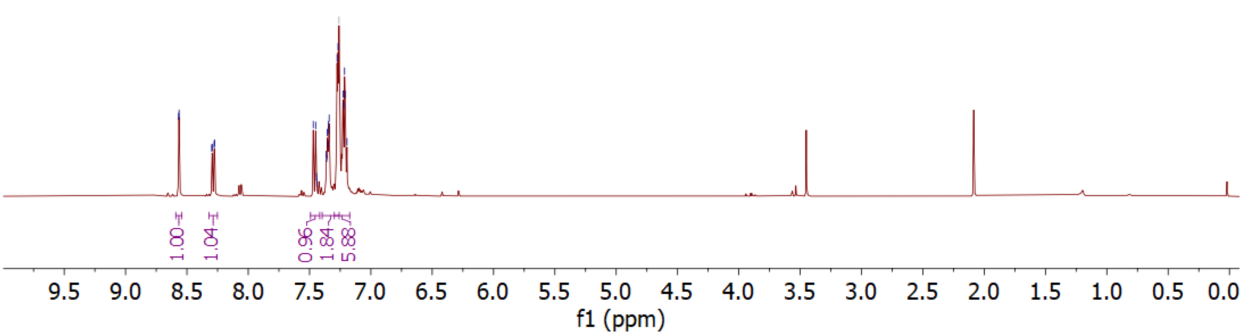

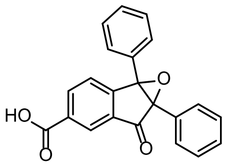

**S6**

| | |
|---|---|
| Origin | Bruker BioSpin GmbH |
| Solvent | CDCl3 |
| Temperature | 298.7 |
| Pulse Sequence | zgpg30 |
| Experiment | 1D |
| Number of Scans | 1024 |
| Acquisition Time | 1.3631 |
| Acquisition Date | 2024-01-05T23:07:00 |
| Spectrometer Frequency | 100.63 |
| Spectral Width | 24038.5 |
| Lowest Frequency | -1946.5 |
| Nucleus | 13C |
| Acquired Size | 32768 |
| Spectral Size | 65536 |

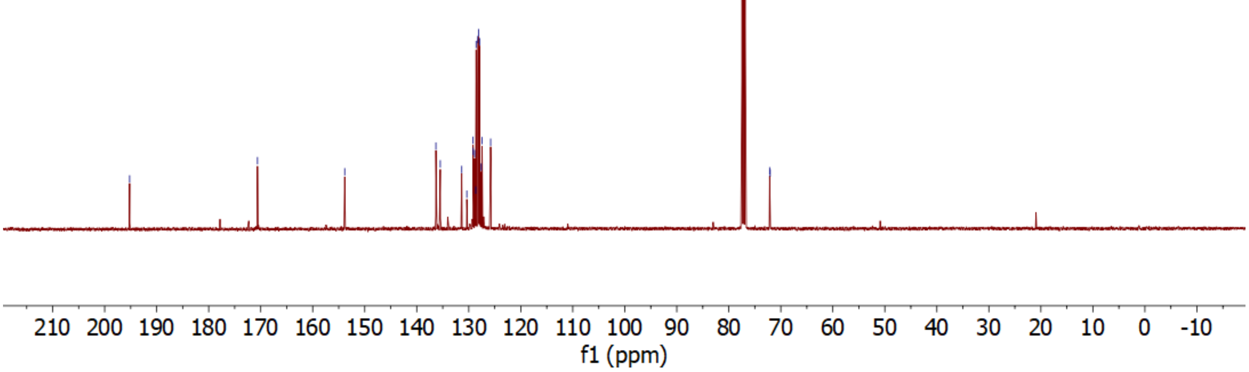

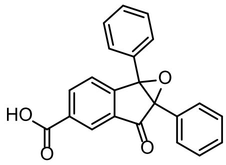

**S6**

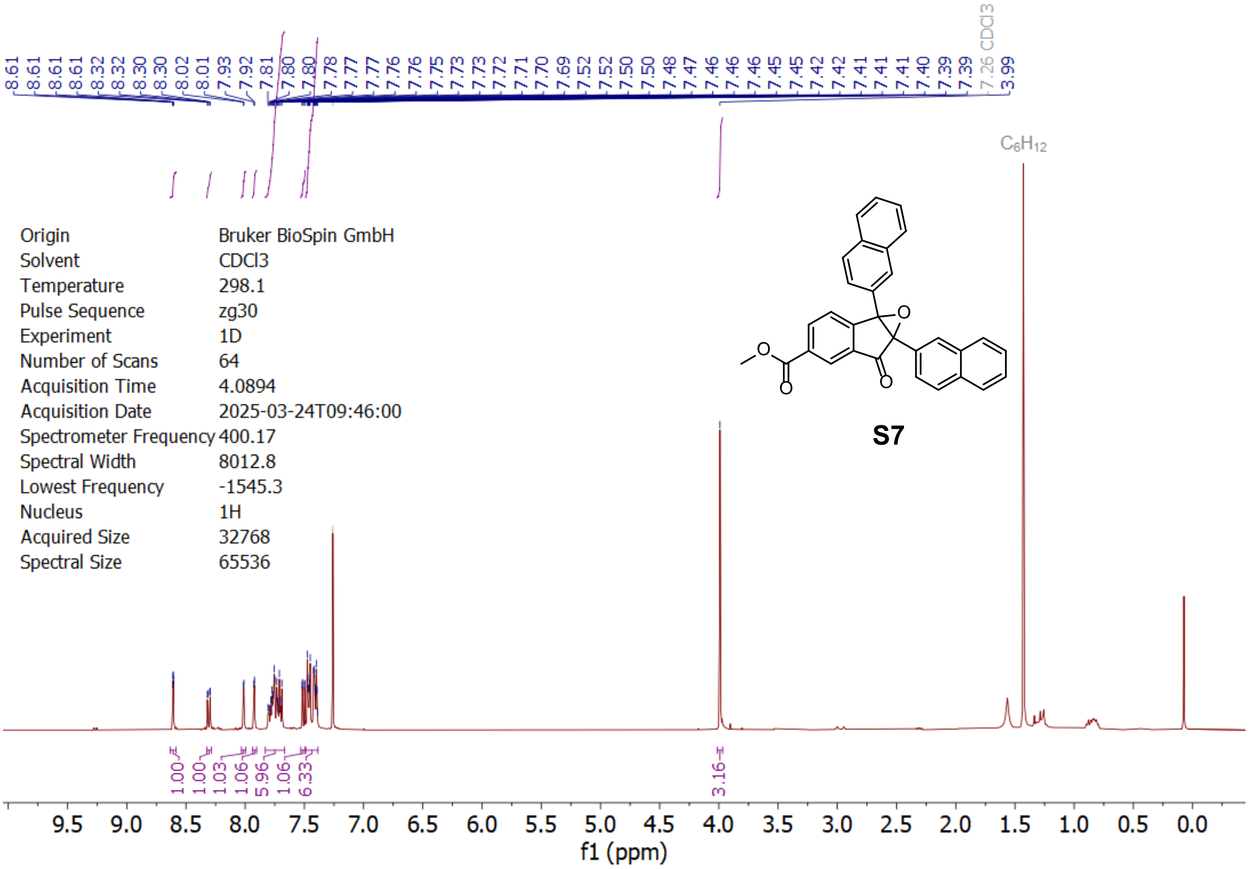
8.61 8.61 8.61 8.61 8.32 8.32 8.30 8.30 8.02 8.01 7.93 7.92 7.81 7.80 7.80 7.78 7.77 7.77 7.76 7.76 7.75 7.73 7.73 7.72 7.71 7.70 7.69 7.52 7.52 7.50 7.50 7.48 7.47 7.46 7.46 7.46 7.45 7.45 7.42 7.42 7.41 7.41 7.41 7.40 7.39 7.39 7.26 CDCl3 3.99
$C_6H_{12}$
Origin Bruker BioSpin GmbH
Solvent CDCl3
Temperature 298.1
Pulse Sequence zg30
Experiment 1D
Number of Scans 64
Acquisition Time 4.0894
Acquisition Date 2025-03-24T09:46:00
Spectrometer Frequency 400.17
Spectral Width 8012.8
Lowest Frequency -1545.3
Nucleus 1H
Acquired Size 32768
Spectral Size 65536
S7
1.00 1.00 1.03 1.06 5.96 1.06 6.33 3.16
9.5 9.0 8.5 8.0 7.5 7.0 6.5 6.0 5.5 5.0 4.5 4.0 3.5 3.0 2.5 2.0 1.5 1.0 0.5 0.0
f1 (ppm)

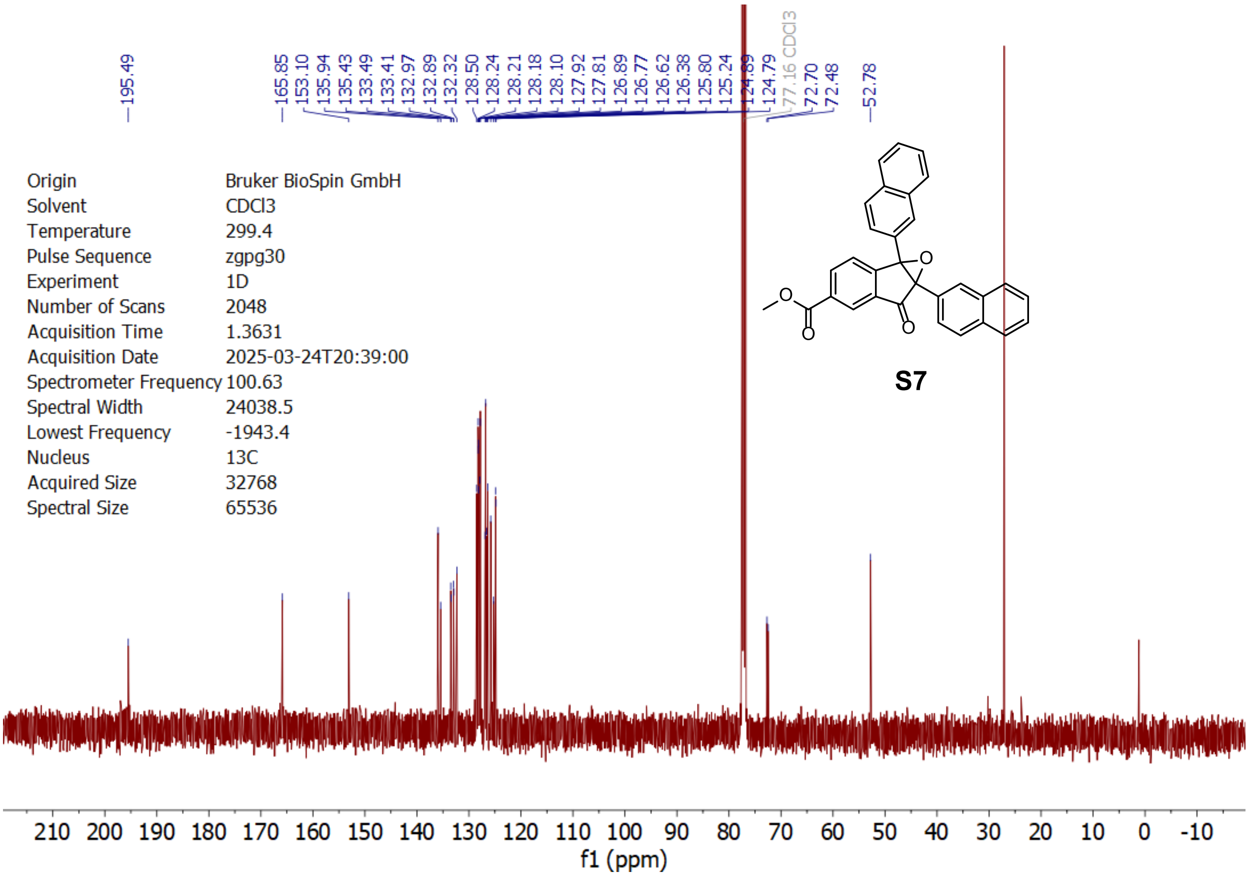
195.49 165.85 153.10 135.94 135.43 133.49 133.41 132.97 132.89 132.32 128.50 128.24 128.21 128.18 128.10 127.92 127.81 126.89 126.77 126.62 126.38 125.80 125.24 124.89 124.79 77.16 CDCl3 72.70 72.48 52.78
Origin Bruker BioSpin GmbH
Solvent CDCl3
Temperature 299.4
Pulse Sequence zgpg30
Experiment 1D
Number of Scans 2048
Acquisition Time 1.3631
Acquisition Date 2025-03-24T20:39:00
Spectrometer Frequency 100.63
Spectral Width 24038.5
Lowest Frequency -1943.4
Nucleus 13C
Acquired Size 32768
Spectral Size 65536
S7
210 200 190 180 170 160 150 140 130 120 110 100 90 80 70 60 50 40 30 20 10 0 -10
f1 (ppm)

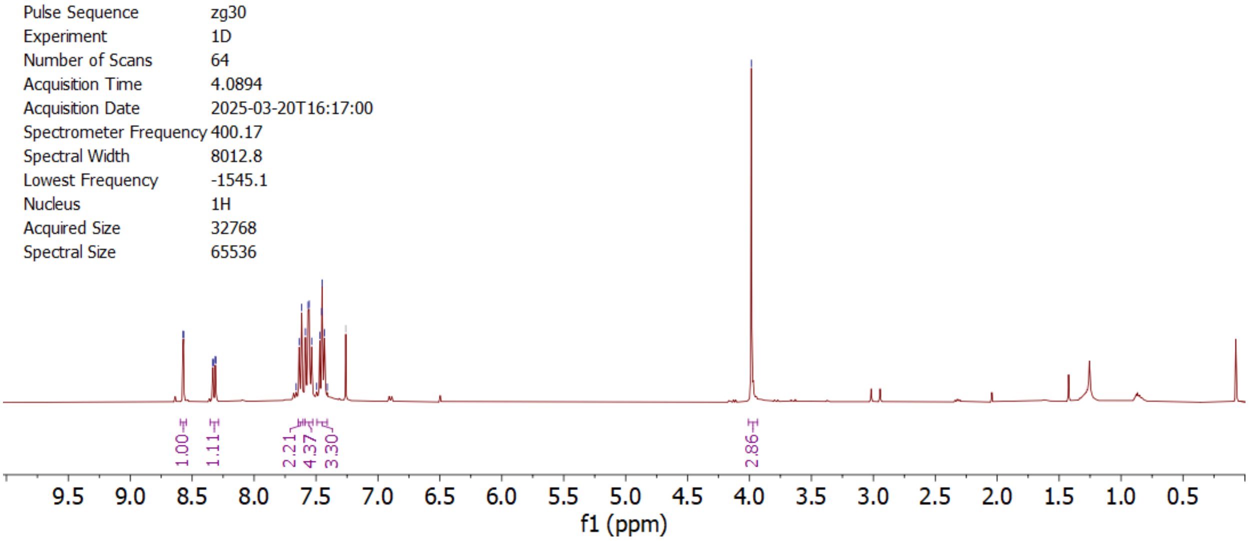

| | |
|---|---|
| Origin | Bruker BioSpin GmbH |
| Solvent | CDCl3 |
| Temperature | 297.9 |
| Pulse Sequence | zg30 |
| Experiment | 1D |
| Number of Scans | 64 |
| Acquisition Time | 4.0894 |
| Acquisition Date | 2025-03-20T16:17:00 |
| Spectrometer Frequency | 400.17 |
| Spectral Width | 8012.8 |
| Lowest Frequency | -1545.1 |
| Nucleus | 1H |
| Acquired Size | 32768 |
| Spectral Size | 65536 |

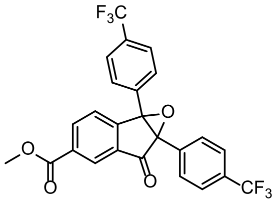

**S8**

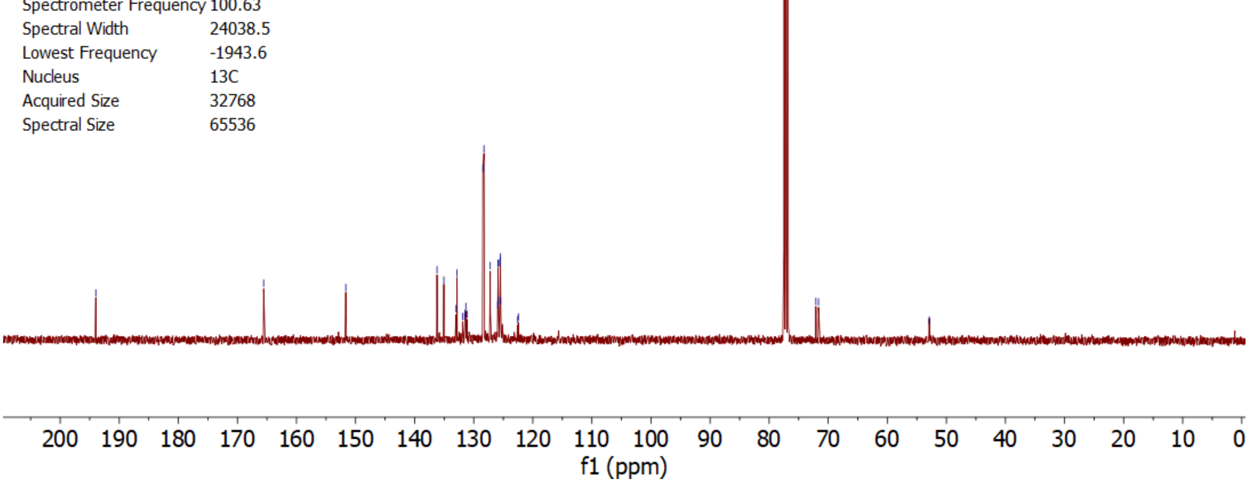

| | |
|---|---|
| Origin | Bruker BioSpin GmbH |
| Solvent | CDCl3 |
| Temperature | 298.7 |
| Pulse Sequence | zgpg30 |
| Experiment | 1D |
| Number of Scans | 1024 |
| Acquisition Time | 1.3631 |
| Acquisition Date | 2025-03-21T06:34:00 |
| Spectrometer Frequency | 100.63 |
| Spectral Width | 24038.5 |
| Lowest Frequency | -1943.6 |
| Nucleus | 13C |
| Acquired Size | 32768 |
| Spectral Size | 65536 |

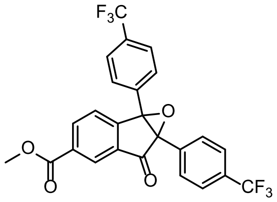

**S8**

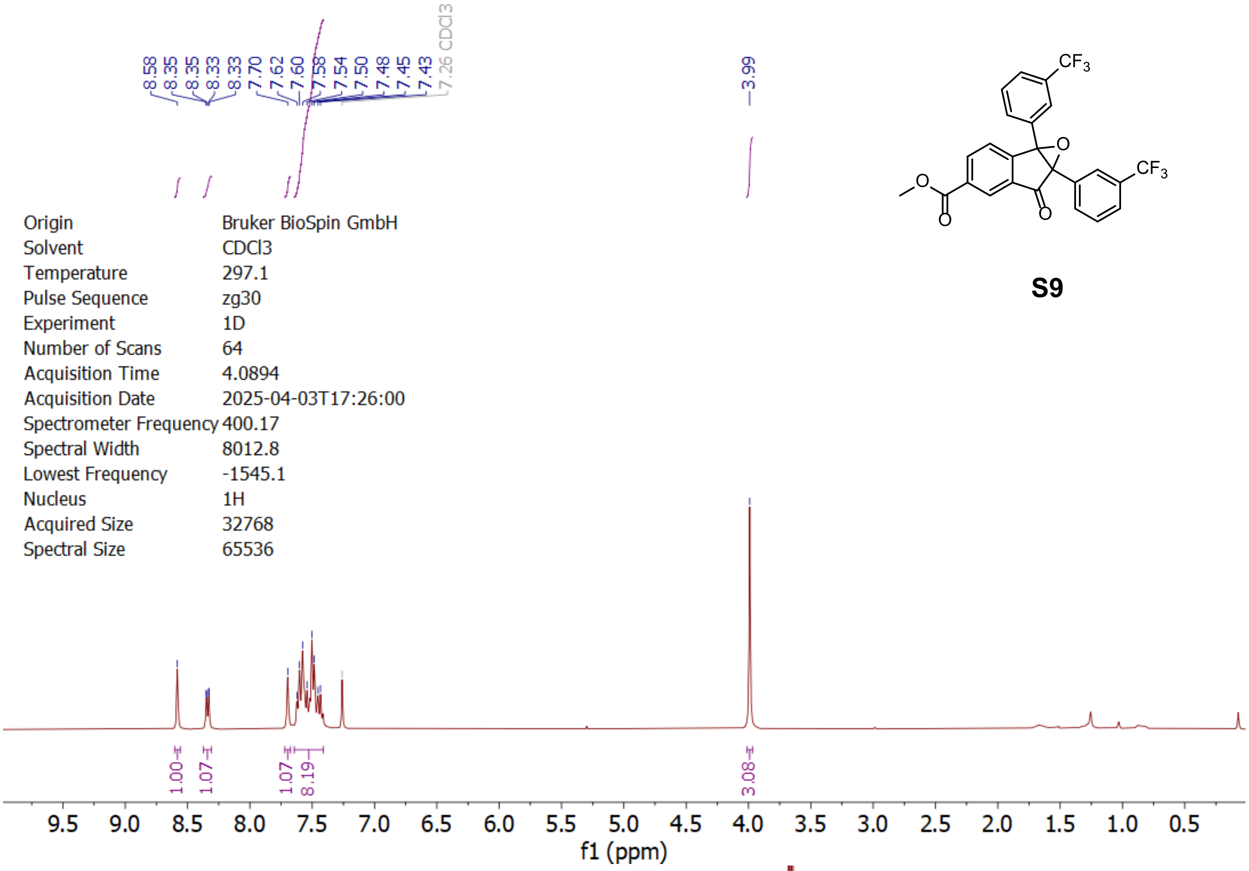
8.58
8.35
8.35
8.33
8.33
7.70
7.62
7.60
7.58
7.54
7.50
7.48
7.45
7.43
7.26 CDCl3
3.99
Origin Bruker BioSpin GmbH
Solvent CDCl3
Temperature 297.1
Pulse Sequence zg30
Experiment 1D
Number of Scans 64
Acquisition Time 4.0894
Acquisition Date 2025-04-03T17:26:00
Spectrometer Frequency 400.17
Spectral Width 8012.8
Lowest Frequency -1545.1
Nucleus 1H
Acquired Size 32768
Spectral Size 65536
1.00
1.07
1.07
8.19
3.08
f1 (ppm)
CF3
O
O
O
O
CF3

**S9**

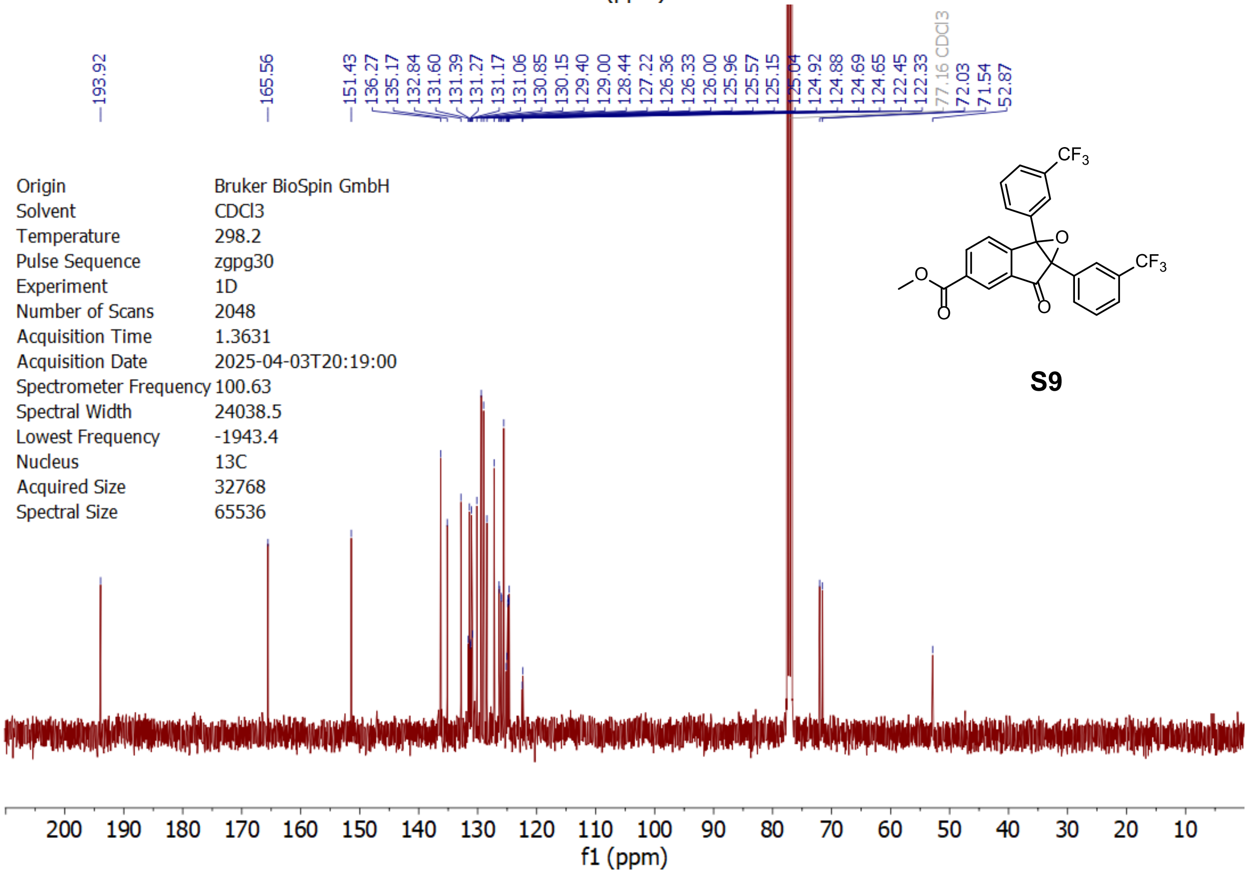
193.92
165.56
151.43
136.27
135.17
132.84
131.60
131.39
131.27
131.17
131.06
130.85
130.15
129.40
129.00
128.44
127.22
126.36
126.33
126.00
125.96
125.57
125.15
125.04
124.92
124.88
124.69
124.65
122.45
122.33
77.16 CDCl3
72.03
71.54
52.87
Origin Bruker BioSpin GmbH
Solvent CDCl3
Temperature 298.2
Pulse Sequence zgpg30
Experiment 1D
Number of Scans 2048
Acquisition Time 1.3631
Acquisition Date 2025-04-03T20:19:00
Spectrometer Frequency 100.63
Spectral Width 24038.5
Lowest Frequency -1943.4
Nucleus 13C
Acquired Size 32768
Spectral Size 65536
f1 (ppm)
CF3
O
O
O
O
CF3

**S9**

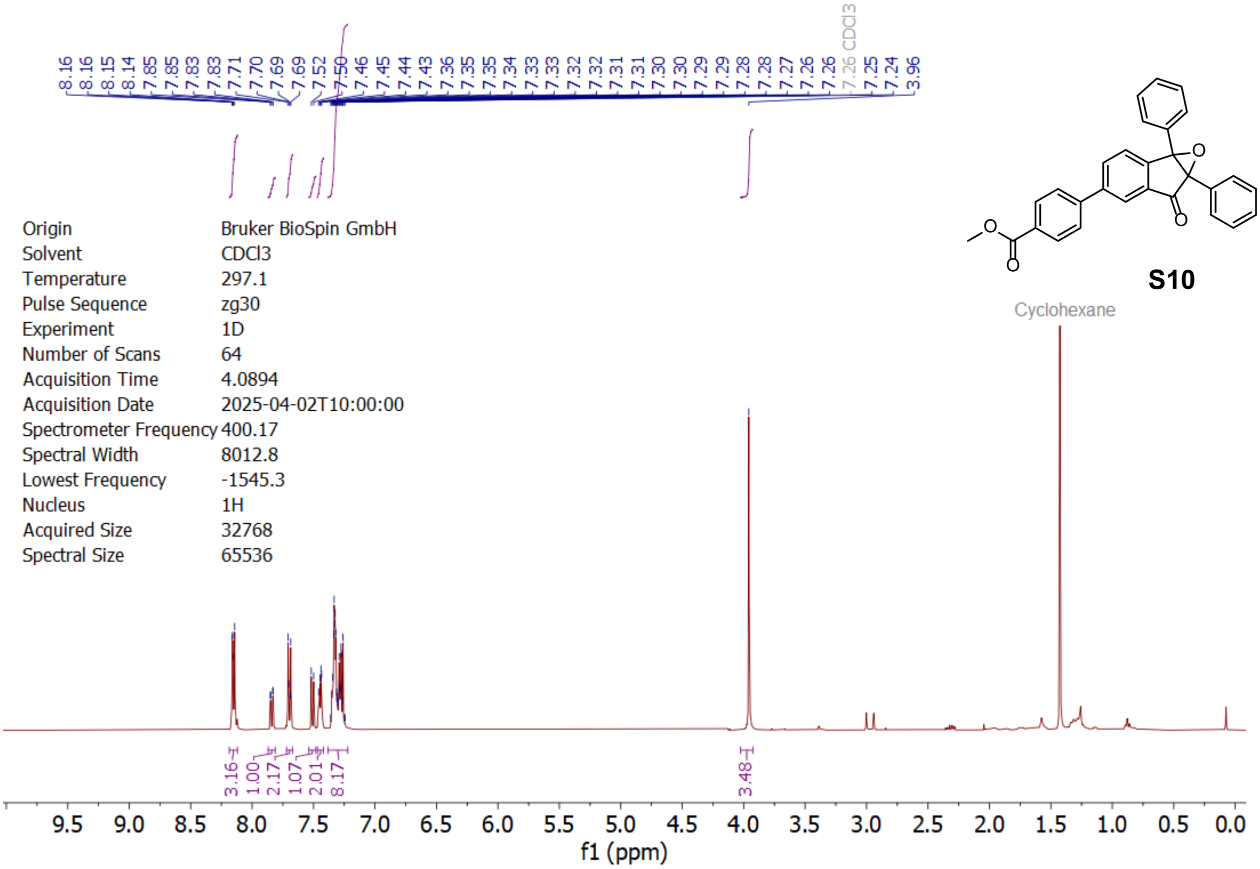
8.16 8.16 8.15 8.14 7.85 7.85 7.83 7.83 7.71 7.70 7.69 7.69 7.52 7.50 7.46 7.45 7.44 7.43 7.36 7.35 7.35 7.34 7.33 7.33 7.32 7.32 7.31 7.31 7.30 7.30 7.29 7.29 7.28 7.28 7.27 7.26 7.26 7.26 CDCl3 7.25 7.24 3.96
S10
Cyclohexane
Origin Bruker BioSpin GmbH
Solvent CDCl3
Temperature 297.1
Pulse Sequence zg30
Experiment 1D
Number of Scans 64
Acquisition Time 4.0894
Acquisition Date 2025-04-02T10:00:00
Spectrometer Frequency 400.17
Spectral Width 8012.8
Lowest Frequency -1545.3
Nucleus 1H
Acquired Size 32768
Spectral Size 65536
3.16 1.00 2.17 1.07 2.01 8.17 3.48
9.5 9.0 8.5 8.0 7.5 7.0 6.5 6.0 5.5 5.0 4.5 4.0 3.5 3.0 2.5 2.0 1.5 1.0 0.5 0.0
f1 (ppm)

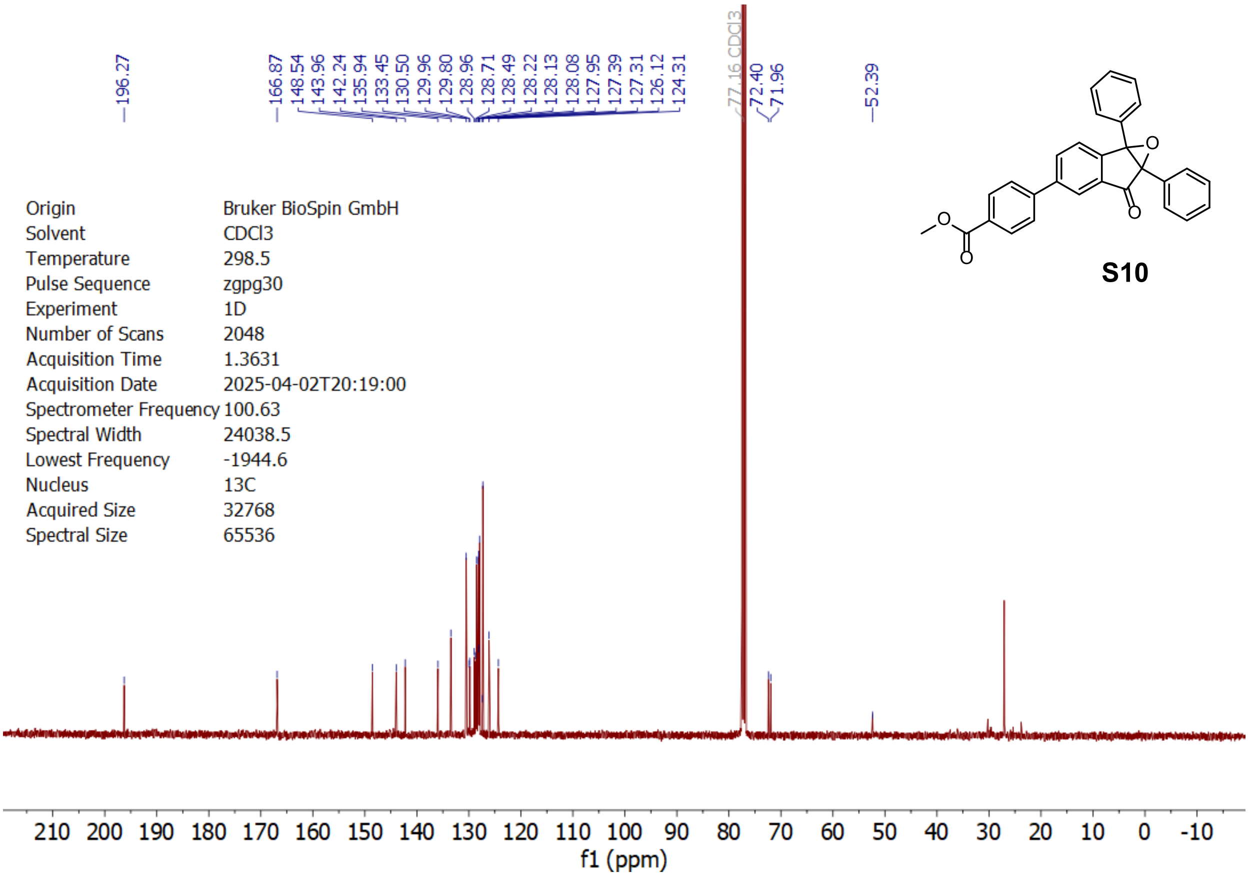
196.27 166.87 148.54 143.96 142.24 135.94 133.45 130.50 129.96 129.80 128.96 128.71 128.49 128.22 128.13 128.08 127.95 127.39 127.31 126.12 124.31 77.16 CDCl3 72.40 71.96 52.39
S10
Origin Bruker BioSpin GmbH
Solvent CDCl3
Temperature 298.5
Pulse Sequence zgpg30
Experiment 1D
Number of Scans 2048
Acquisition Time 1.3631
Acquisition Date 2025-04-02T20:19:00
Spectrometer Frequency 100.63
Spectral Width 24038.5
Lowest Frequency -1944.6
Nucleus 13C
Acquired Size 32768
Spectral Size 65536
210 200 190 180 170 160 150 140 130 120 110 100 90 80 70 60 50 40 30 20 10 0 -10
f1 (ppm)

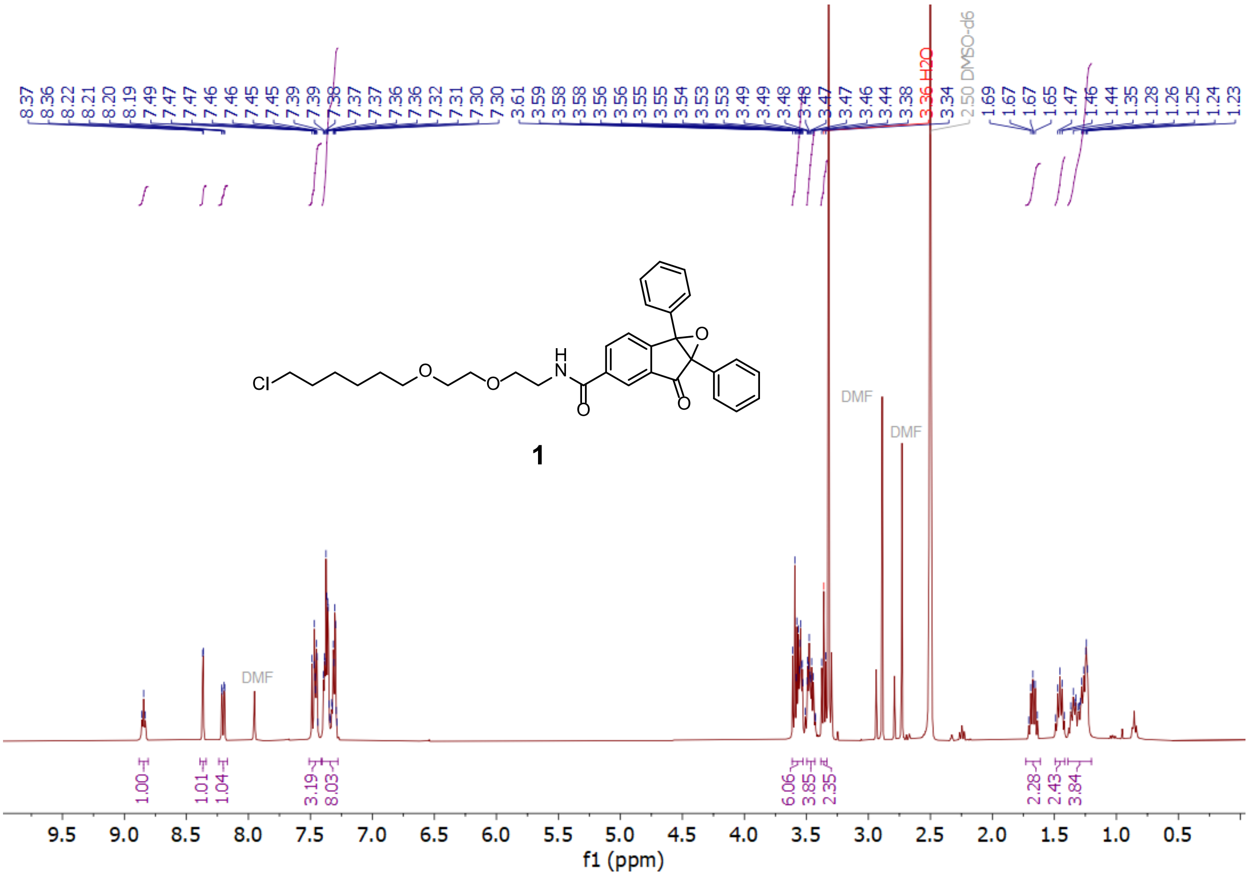
8.37 8.36 8.22 8.21 8.20 8.19 7.49 7.47 7.47 7.46 7.46 7.45 7.45 7.39 7.39 7.38 7.37 7.37 7.36 7.36 7.32 7.31 7.30 7.30 3.61 3.59 3.58 3.58 3.56 3.56 3.55 3.55 3.54 3.53 3.53 3.49 3.49 3.48 3.48 3.47 3.47 3.46 3.44 3.38 3.36 H2O 3.34 2.50 DMSO-d6 1.69 1.67 1.67 1.65 1.47 1.46 1.44 1.35 1.28 1.26 1.25 1.24 1.23
Cl
O
O
H
N
O
O
O
1
DMF
DMF
DMF
1.00
1.01
1.04
3.19
8.03
6.06
3.85
2.35
2.28
2.43
3.84
9.5 9.0 8.5 8.0 7.5 7.0 6.5 6.0 5.5 5.0 4.5 4.0 3.5 3.0 2.5 2.0 1.5 1.0 0.5
f1 (ppm)

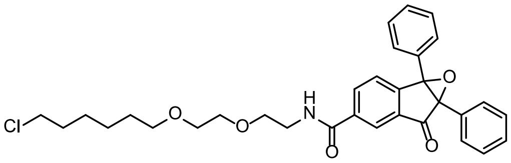
Cl
O
O
H
N
O
O
O

**1**

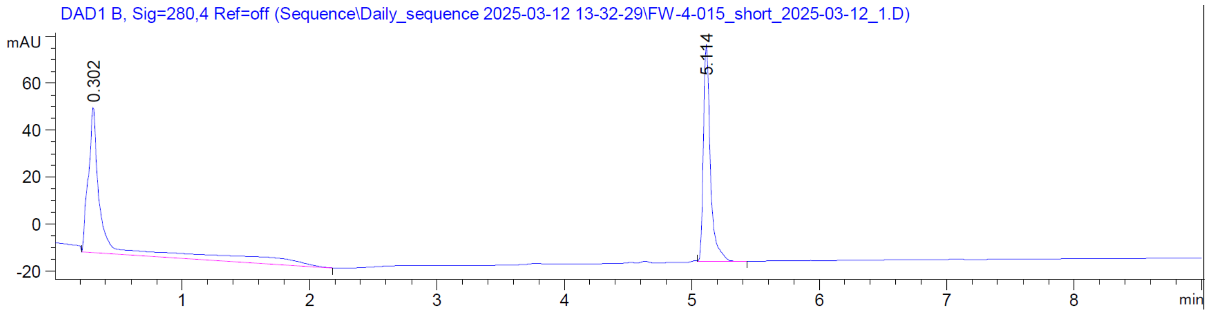
DAD1 B, Sig=280,4 Ref=off (Sequence\Daily_sequence 2025-03-12 13-32-29\FW-4-015_short_2025-03-12_1.D)
mAU
60
40
20
0
-20
0.302
5.114
1 2 3 4 5 6 7 8
min

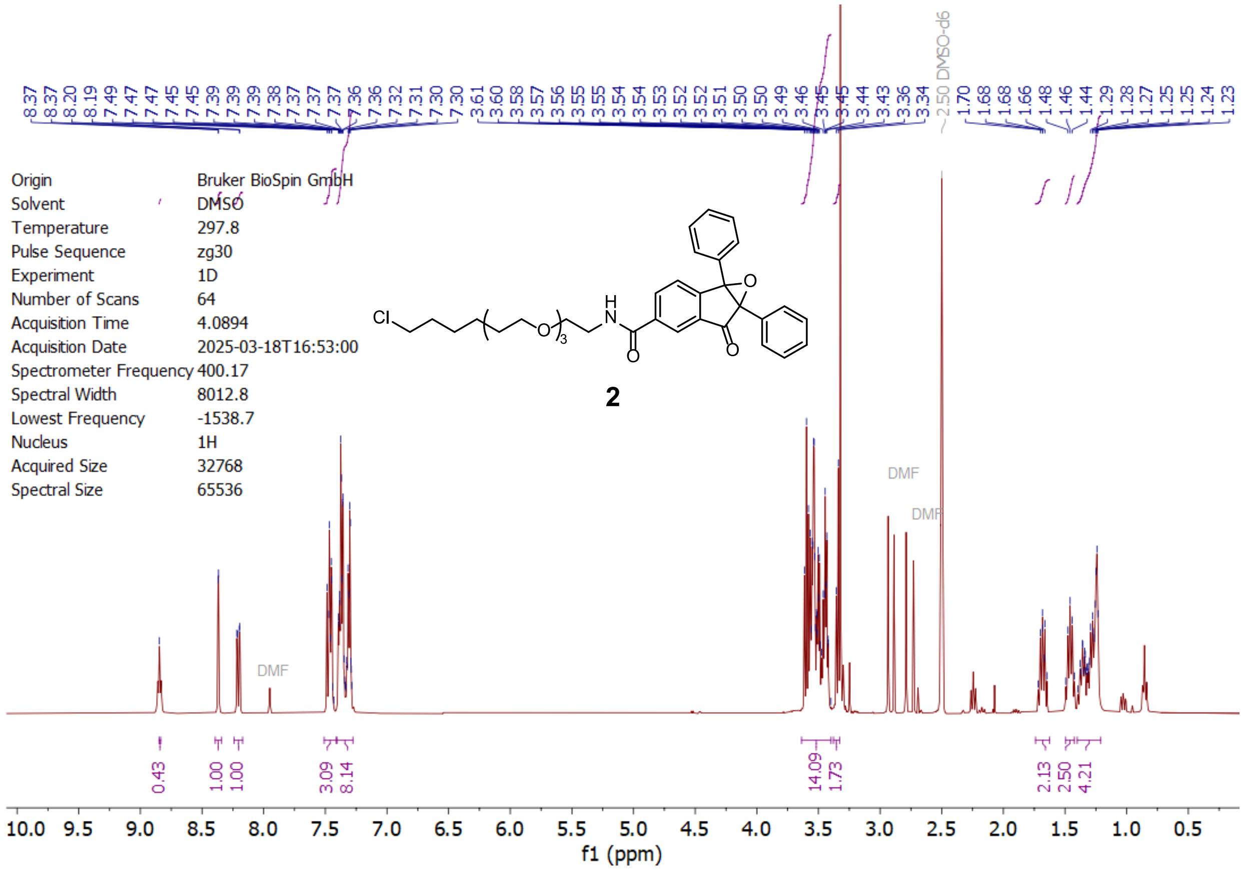

8.37 8.37 8.20 8.19 7.49 7.47 7.47 7.45 7.45 7.39 7.39 7.39 7.38 7.37 7.37 7.37 7.36 7.36 7.32 7.31 7.30 7.30 3.61 3.60 3.58 3.57 3.56 3.55 3.55 3.54 3.54 3.53 3.52 3.52 3.51 3.50 3.50 3.49 3.46 3.45 3.45 3.44 3.43 3.36 3.34 2.50 DMSO-d6 1.70 1.68 1.68 1.66 1.48 1.46 1.44 1.29 1.28 1.27 1.25 1.25 1.24 1.23
Origin Bruker BioSpin GmbH
Solvent DMSO
Temperature 297.8
Pulse Sequence zg30
Experiment 1D
Number of Scans 64
Acquisition Time 4.0894
Acquisition Date 2025-03-18T16:53:00
Spectrometer Frequency 400.17
Spectral Width 8012.8
Lowest Frequency -1538.7
Nucleus 1H
Acquired Size 32768
Spectral Size 65536
Cl
H
N
O
O
O
3
2
DMF
DMF
DMF
0.43 1.00 1.00 3.09 8.14 14.09 1.73 2.13 2.50 4.21
10.0 9.5 9.0 8.5 8.0 7.5 7.0 6.5 6.0 5.5 5.0 4.5 4.0 3.5 3.0 2.5 2.0 1.5 1.0 0.5
f1 (ppm)

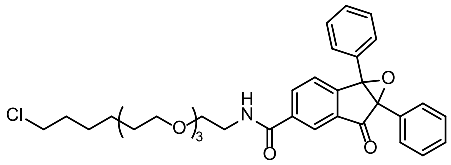

Cl
H
N
O
O
O
3

**2**

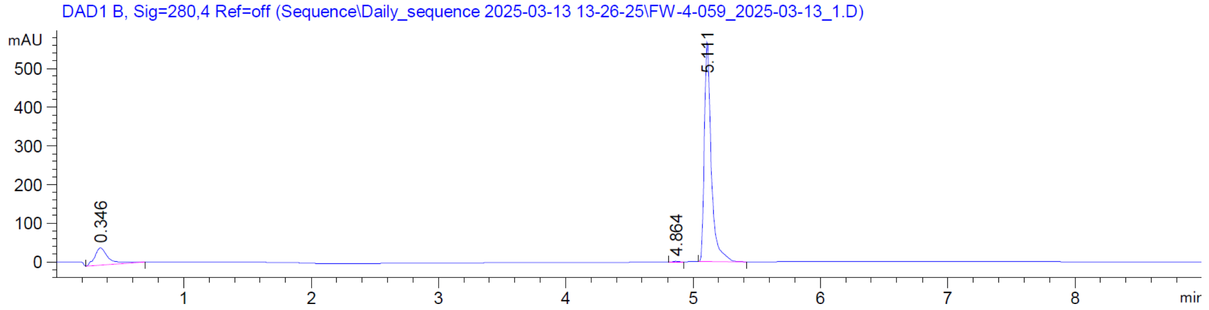

DAD1 B, Sig=280,4 Ref=off (Sequence\Daily_sequence 2025-03-13 13-26-25\FW-4-059_2025-03-13_1.D)
mAU
500
400
300
200
100
0
0.346
4.864
5.111
1 2 3 4 5 6 7 8
min

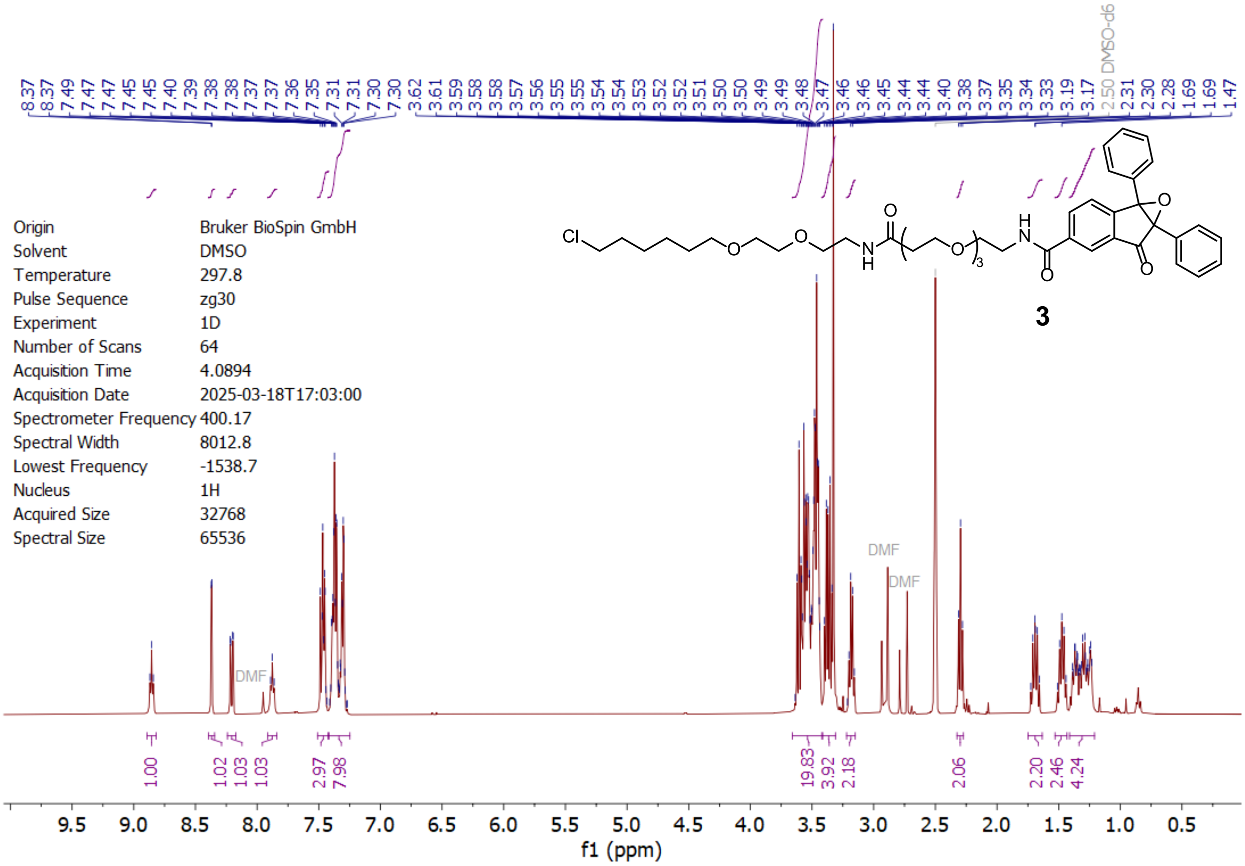

Origin Bruker BioSpin GmbH
Solvent DMSO
Temperature 297.8
Pulse Sequence zg30
Experiment 1D
Number of Scans 64
Acquisition Time 4.0894
Acquisition Date 2025-03-18T17:03:00
Spectrometer Frequency 400.17
Spectral Width 8012.8
Lowest Frequency -1538.7
Nucleus 1H
Acquired Size 32768
Spectral Size 65536
3
DMF
2.50 DMSO-d6
1.00
1.02
1.03
1.03
2.97
7.98
19.83
3.92
2.18
2.06
2.20
2.46
4.24
f1 (ppm)

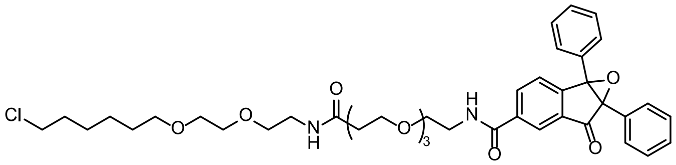

**3**

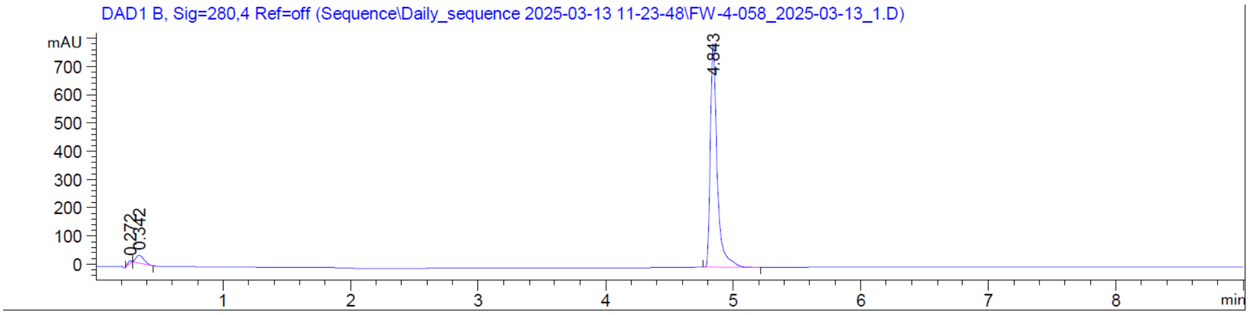

DAD1 B, Sig=280,4 Ref=off (Sequence\Daily_sequence 2025-03-13 11-23-48\FW-4-058_2025-03-13_1.D)
mAU
4.843
0.272
0.342
min

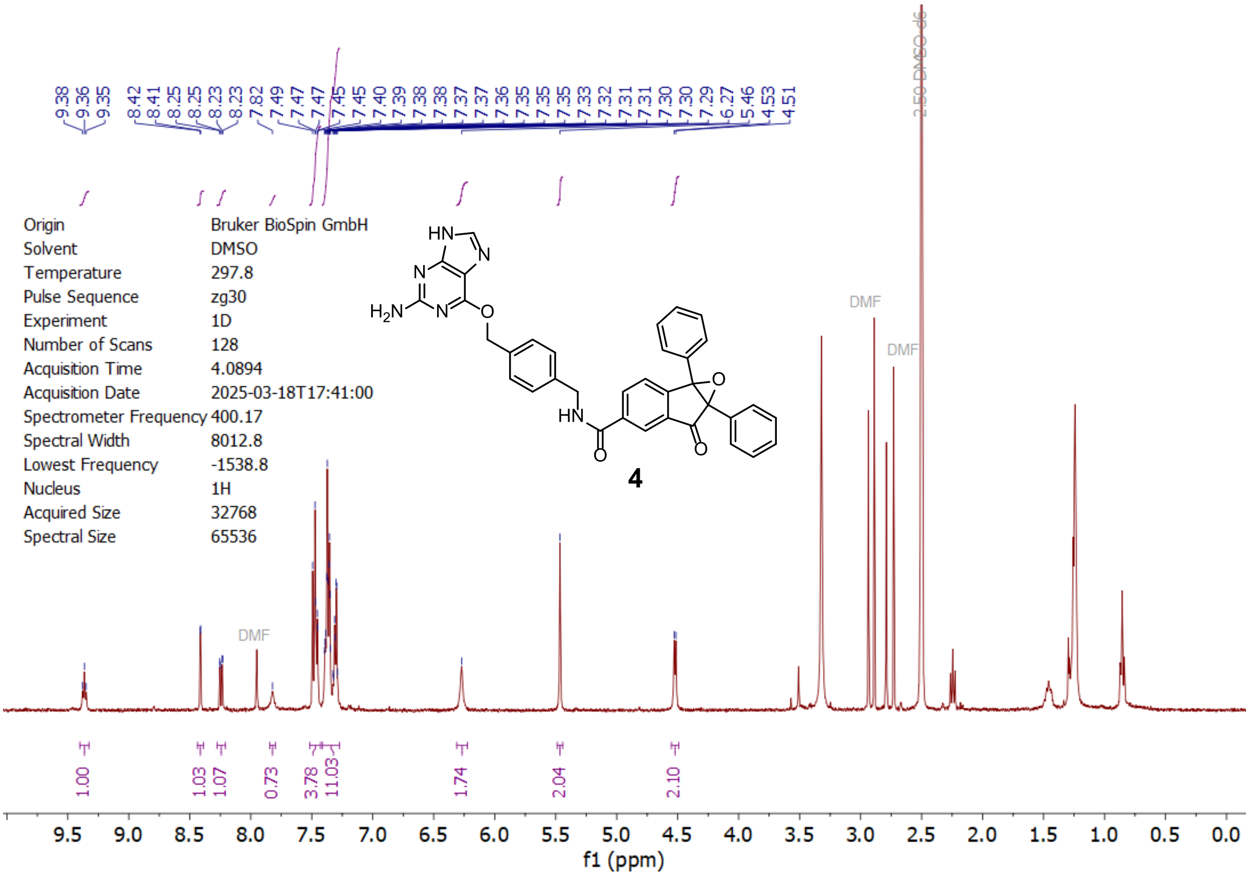
Origin Bruker BioSpin GmbH
Solvent DMSO
Temperature 297.8
Pulse Sequence zg30
Experiment 1D
Number of Scans 128
Acquisition Time 4.0894
Acquisition Date 2025-03-18T17:41:00
Spectrometer Frequency 400.17
Spectral Width 8012.8
Lowest Frequency -1538.8
Nucleus 1H
Acquired Size 32768
Spectral Size 65536
HN
N
N
H2N
N
O
HN
O
O
O
4
DMF
DMF
DMF
2.50 DMSO-d6
1.00
1.03
1.07
0.73
3.78
11.03
1.74
2.04
2.10
f1 (ppm)

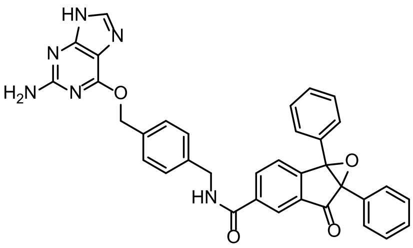
HN
N
N
H2N
N
O
HN
O
O
O

**4**

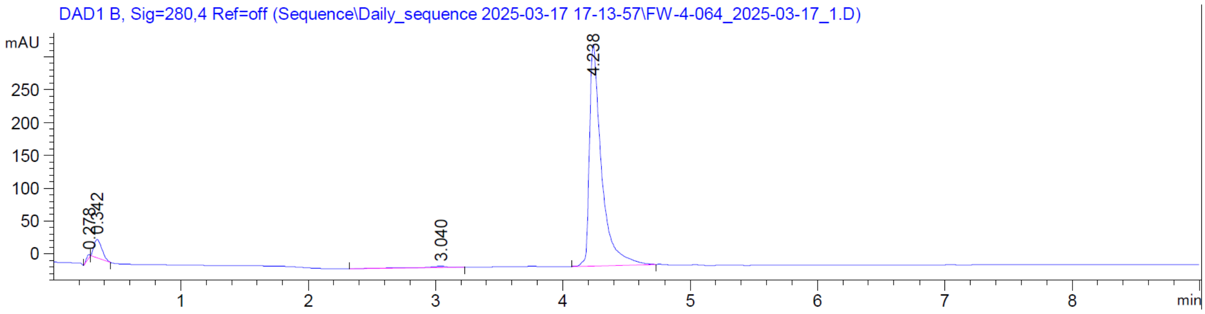
DAD1 B, Sig=280,4 Ref=off (Sequence\Daily_sequence 2025-03-17 17-13-57\FW-4-064_2025-03-17_1.D)
mAU
0.278
0.342
3.040
4.238
min

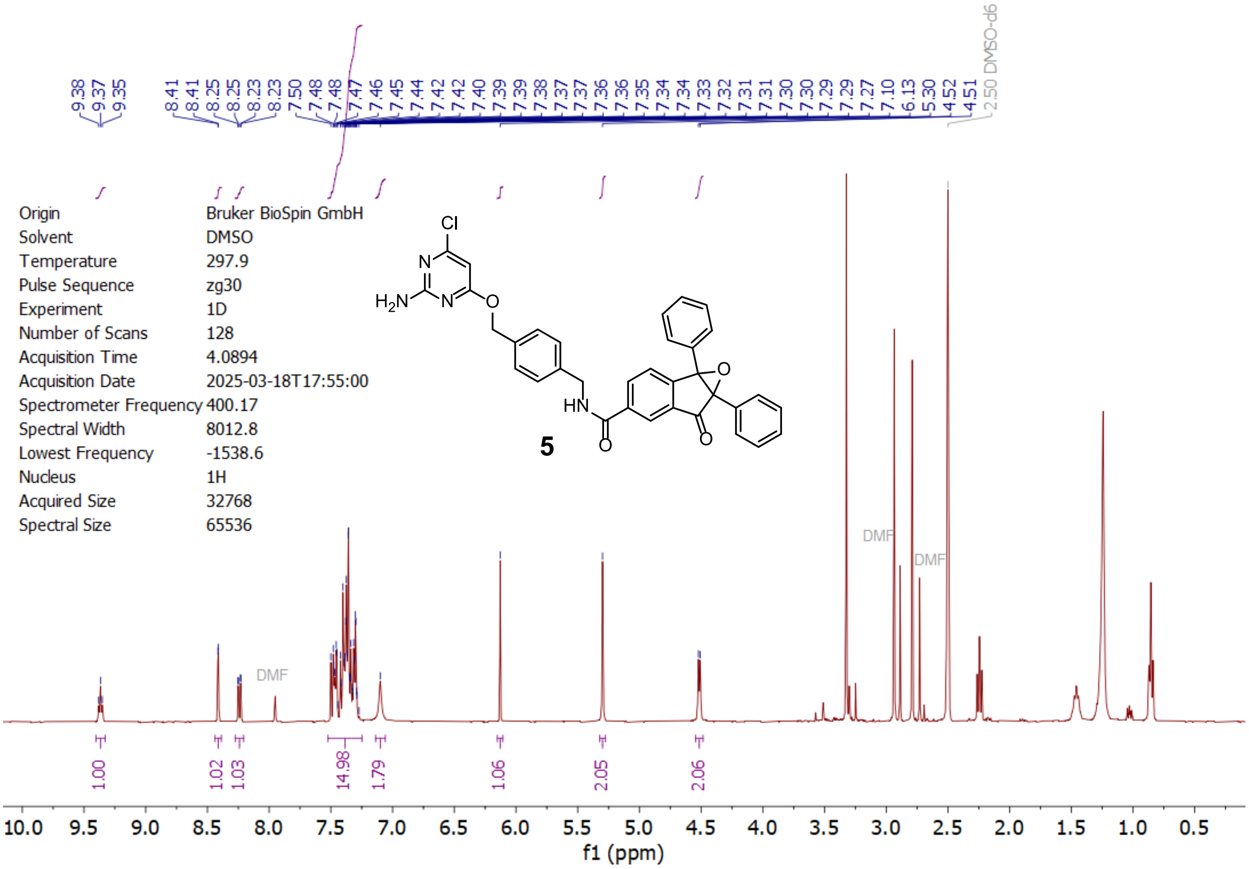

Origin Bruker BioSpin GmbH
Solvent DMSO
Temperature 297.9
Pulse Sequence zg30
Experiment 1D
Number of Scans 128
Acquisition Time 4.0894
Acquisition Date 2025-03-18T17:55:00
Spectrometer Frequency 400.17
Spectral Width 8012.8
Lowest Frequency -1538.6
Nucleus 1H
Acquired Size 32768
Spectral Size 65536
9.38 9.37 9.35 8.41 8.41 8.25 8.25 8.23 8.23 7.50 7.48 7.48 7.47 7.46 7.45 7.44 7.42 7.42 7.40 7.39 7.39 7.38 7.37 7.37 7.36 7.36 7.35 7.34 7.34 7.33 7.32 7.31 7.31 7.30 7.30 7.29 7.29 7.27 7.10 6.13 5.30 4.52 4.51 2.50 DMSO-d6
DMF
DMF
DMF
1.00 1.02 1.03 14.98 1.79 1.06 2.05 2.06
f1 (ppm)
5

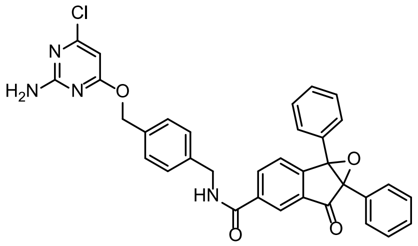

Cl
N
H2N
N
O
HN
O
O
O

**5**

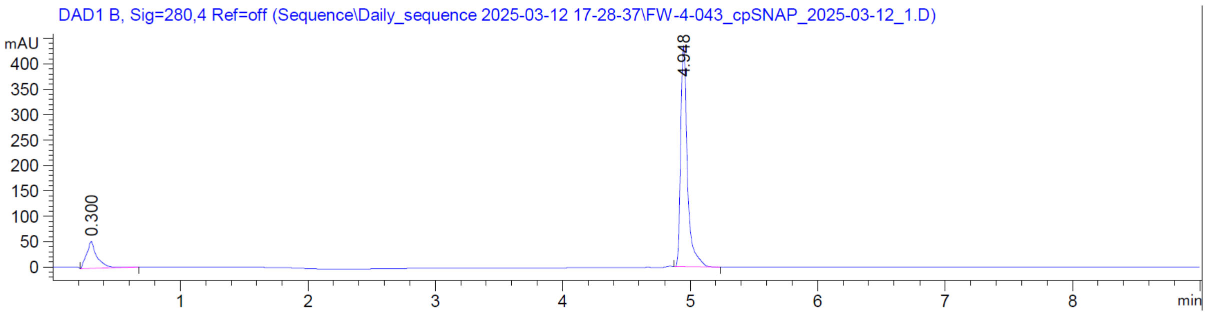

DAD1 B, Sig=280,4 Ref=off (Sequence\Daily_sequence 2025-03-12 17-28-37\FW-4-043_cpSNAP_2025-03-12_1.D)
mAU
0.300
4.948
min

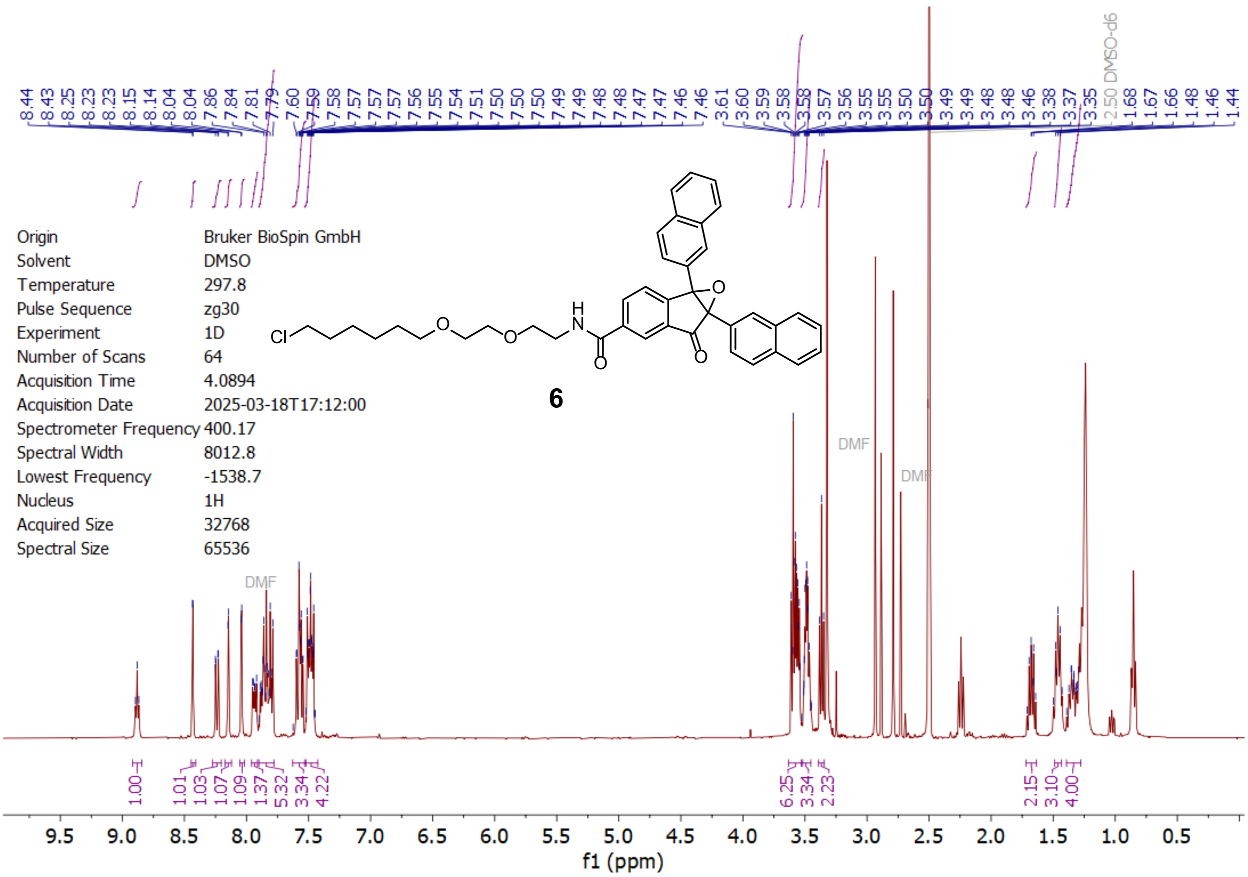

Origin Bruker BioSpin GmbH
Solvent DMSO
Temperature 297.8
Pulse Sequence zg30
Experiment 1D
Number of Scans 64
Acquisition Time 4.0894
Acquisition Date 2025-03-18T17:12:00
Spectrometer Frequency 400.17
Spectral Width 8012.8
Lowest Frequency -1538.7
Nucleus 1H
Acquired Size 32768
Spectral Size 65536
6
DMF
DMSO-d6
f1 (ppm)

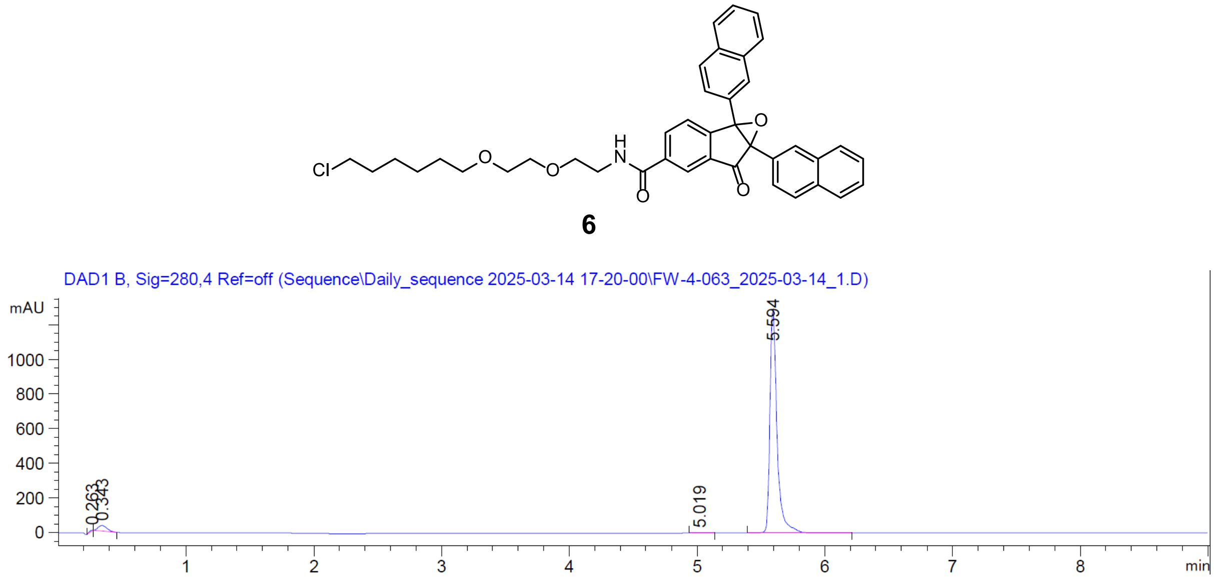

6
DAD1 B, Sig=280,4 Ref=off (Sequence\Daily_sequence 2025-03-14 17-20-00\FW-4-063_2025-03-14_1.D)
mAU
5.594
5.019
min

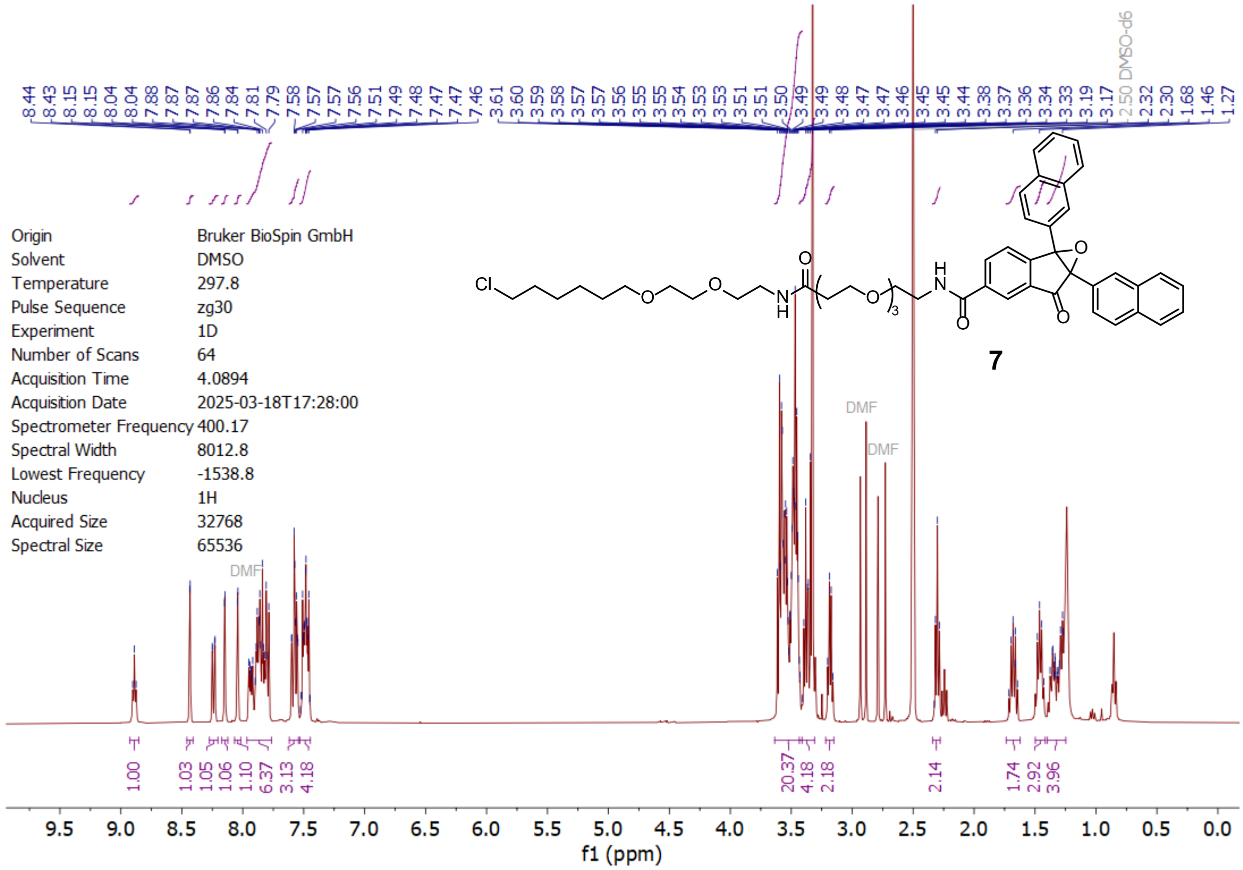

Origin Bruker BioSpin GmbH
Solvent DMSO
Temperature 297.8
Pulse Sequence zg30
Experiment 1D
Number of Scans 64
Acquisition Time 4.0894
Acquisition Date 2025-03-18T17:28:00
Spectrometer Frequency 400.17
Spectral Width 8012.8
Lowest Frequency -1538.8
Nucleus 1H
Acquired Size 32768
Spectral Size 65536
8.44 8.43 8.15 8.15 8.04 8.04 7.88 7.87 7.87 7.86 7.84 7.81 7.79 7.58 7.57 7.57 7.56 7.51 7.49 7.48 7.47 7.47 7.46 3.61 3.60 3.59 3.58 3.57 3.57 3.56 3.55 3.55 3.54 3.53 3.53 3.51 3.50 3.49 3.49 3.48 3.47 3.47 3.46 3.45 3.45 3.44 3.38 3.37 3.36 3.34 3.33 3.19 3.17 2.50 DMSO-d6 2.32 2.30 1.68 1.46 1.27
DMF
DMF
DMF
1.00 1.03 1.05 1.06 1.10 6.37 3.13 4.18 20.37 4.18 2.18 2.14 1.74 2.92 3.96
f1 (ppm)
Cl
O
O
N
H
O
O
3
H
N
O
O
O
7

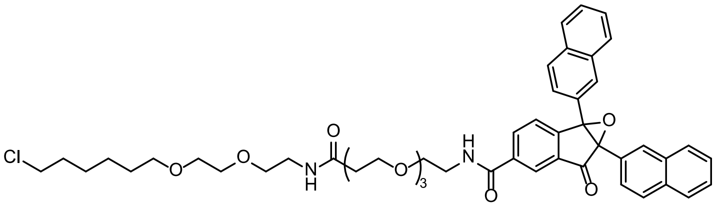

**7**

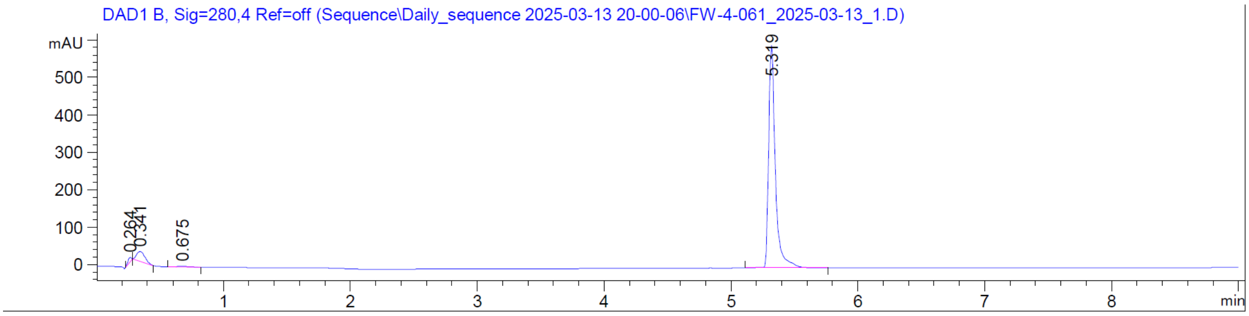

DAD1 B, Sig=280,4 Ref=off (Sequence\Daily_sequence 2025-03-13 20-00-06\FW-4-061_2025-03-13_1.D)
mAU
5.319
0.264
0.341
0.675
min

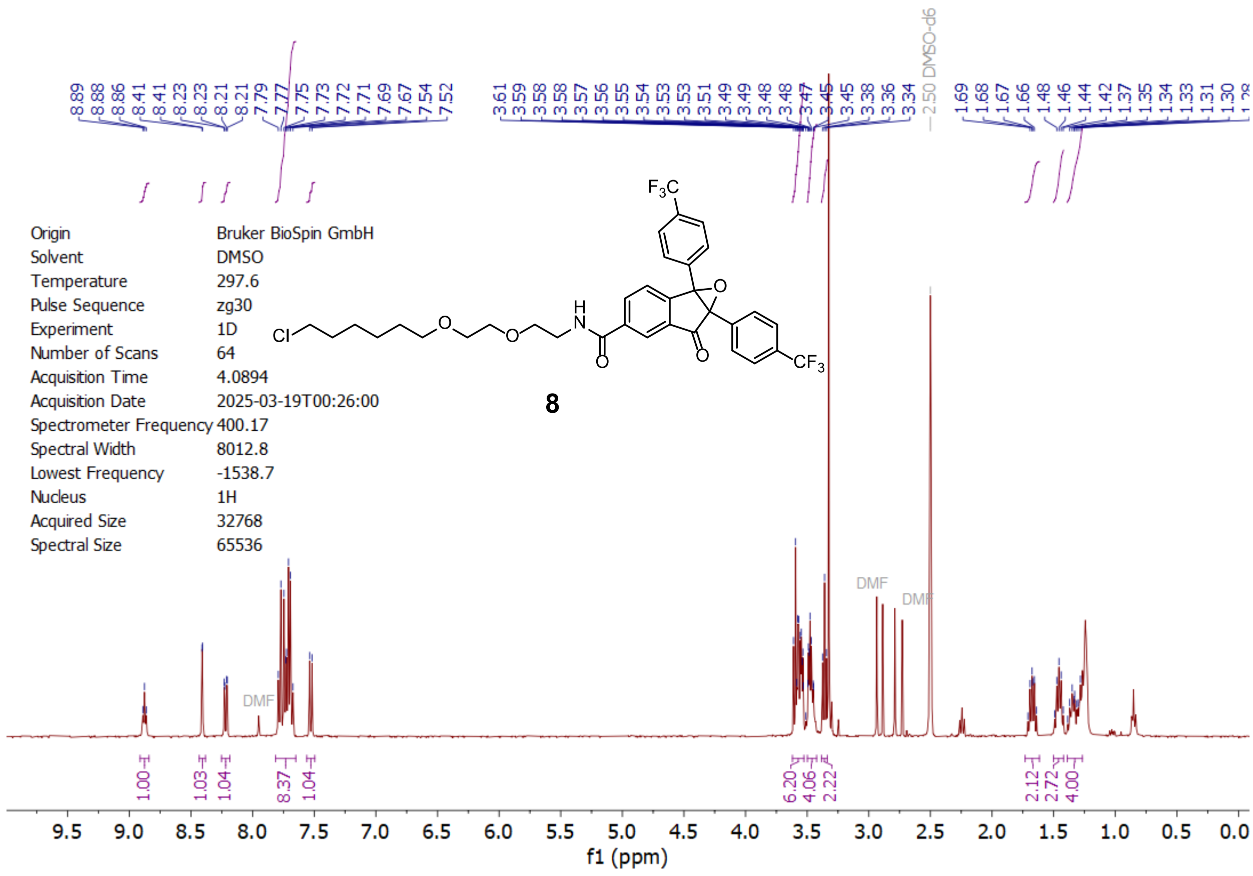

Origin Bruker BioSpin GmbH
Solvent DMSO
Temperature 297.6
Pulse Sequence zg30
Experiment 1D
Number of Scans 64
Acquisition Time 4.0894
Acquisition Date 2025-03-19T00:26:00
Spectrometer Frequency 400.17
Spectral Width 8012.8
Lowest Frequency -1538.7
Nucleus 1H
Acquired Size 32768
Spectral Size 65536
8
DMF
DMSO-d6
f1 (ppm)

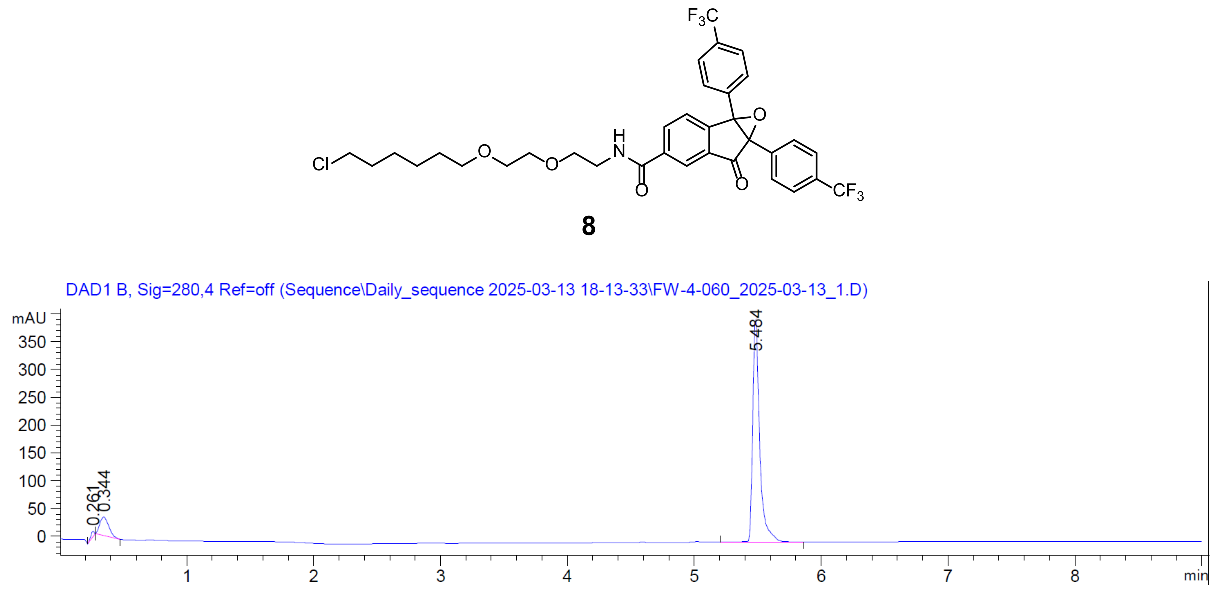

8
DAD1 B, Sig=280,4 Ref=off (Sequence\Daily_sequence 2025-03-13 18-13-33\FW-4-060_2025-03-13_1.D)
mAU
5.484
0.261
0.344
min

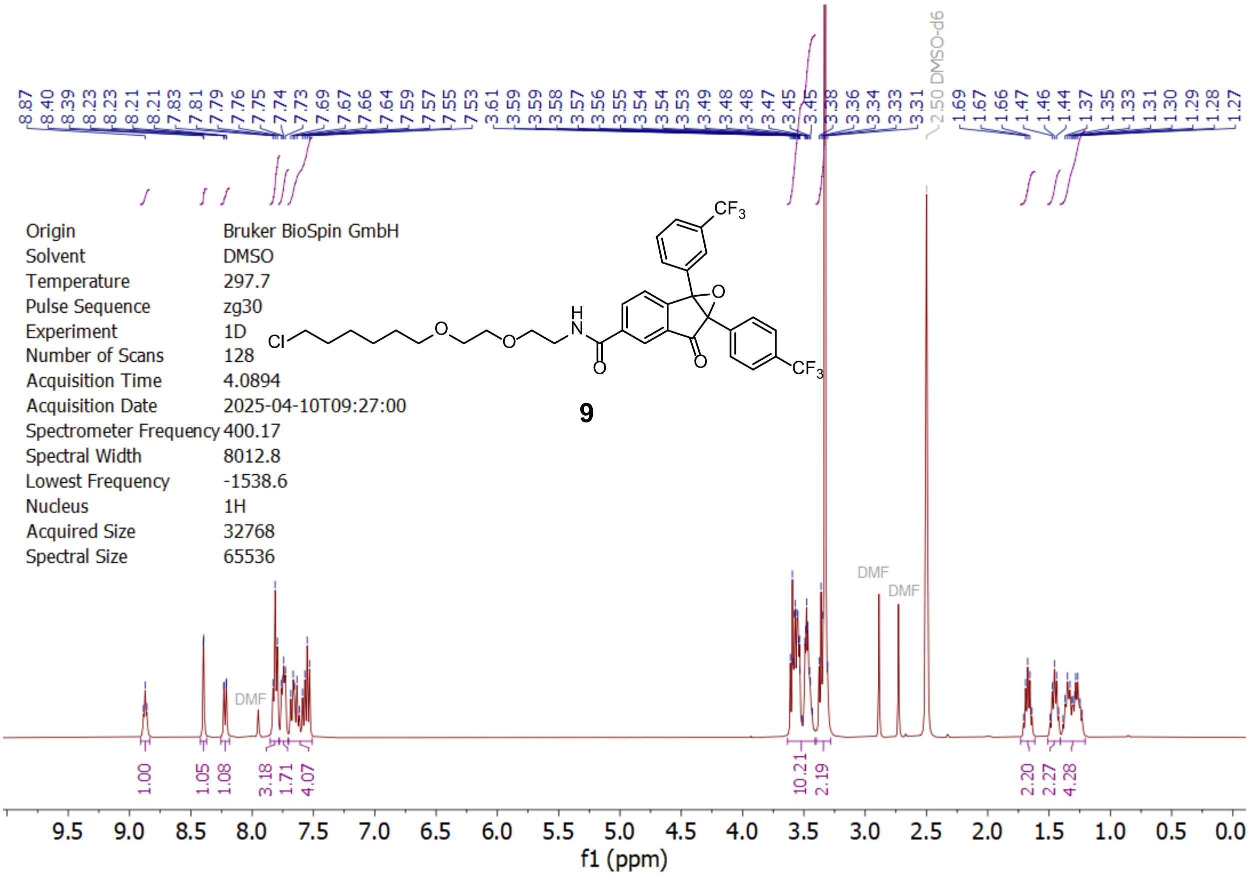

8.87 8.40 8.39 8.23 8.23 8.21 8.21 7.83 7.81 7.79 7.76 7.75 7.74 7.73 7.69 7.67 7.66 7.64 7.59 7.57 7.55 7.53 3.61 3.59 3.59 3.58 3.57 3.56 3.55 3.54 3.54 3.53 3.49 3.48 3.48 3.47 3.45 3.45 3.38 3.36 3.34 3.33 3.31 2.50 DMSO-d6 1.69 1.67 1.66 1.47 1.46 1.44 1.37 1.35 1.33 1.31 1.30 1.29 1.28 1.27
Origin Bruker BioSpin GmbH
Solvent DMSO
Temperature 297.7
Pulse Sequence zg30
Experiment 1D
Number of Scans 128
Acquisition Time 4.0894
Acquisition Date 2025-04-10T09:27:00
Spectrometer Frequency 400.17
Spectral Width 8012.8
Lowest Frequency -1538.6
Nucleus 1H
Acquired Size 32768
Spectral Size 65536
Cl
O
O
H
N
O
O
O
CF3
CF3
9
DMF
DMF
DMF
1.00 1.05 1.08 3.18 1.71 4.07 10.21 2.19 2.20 2.27 4.28
9.5 9.0 8.5 8.0 7.5 7.0 6.5 6.0 5.5 5.0 4.5 4.0 3.5 3.0 2.5 2.0 1.5 1.0 0.5 0.0
f1 (ppm)

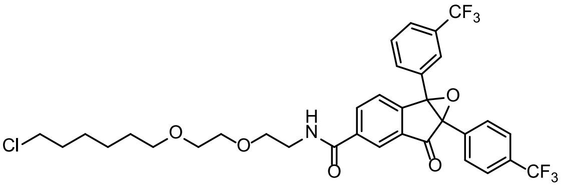

Cl
O
O
H
N
O
O
O
CF3
CF3

**9**

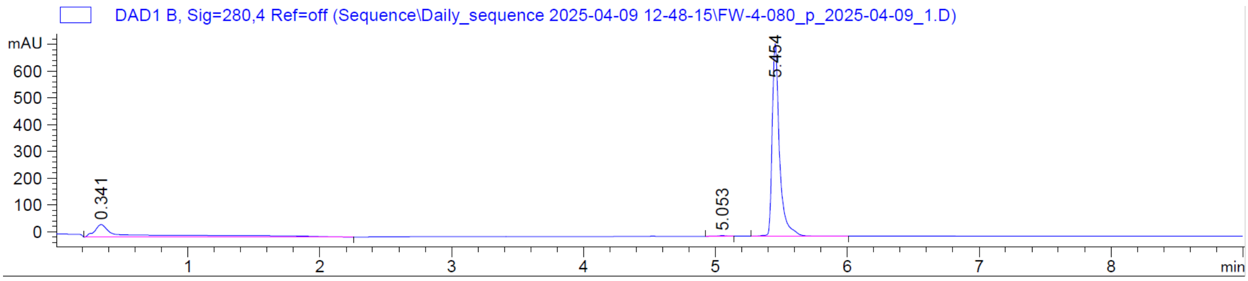

DAD1 B, Sig=280,4 Ref=off (Sequence\Daily_sequence 2025-04-09 12-48-15\FW-4-080_p_2025-04-09_1.D)
mAU
600
500
400
300
200
100
0
0.341
5.053
5.454
1 2 3 4 5 6 7 8
min

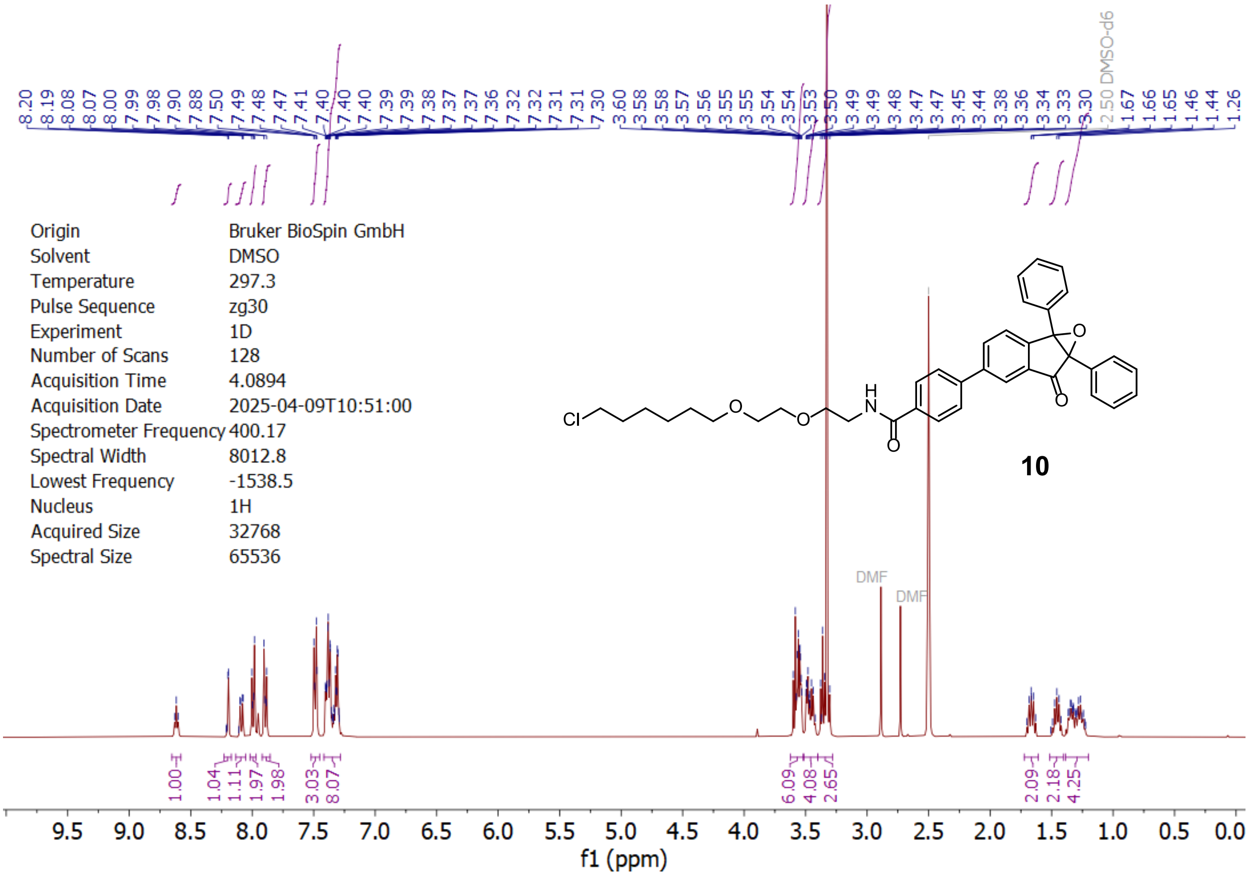
8.20 8.19 8.08 8.07 8.00 7.99 7.98 7.90 7.88 7.50 7.49 7.48 7.47 7.41 7.40 7.40 7.40 7.39 7.39 7.38 7.37 7.37 7.36 7.32 7.32 7.31 7.31 7.30 3.60 3.58 3.58 3.57 3.56 3.55 3.55 3.54 3.54 3.53 3.50 3.49 3.49 3.48 3.47 3.47 3.45 3.44 3.38 3.36 3.34 3.33 3.30 2.50 DMSO-d6 1.67 1.66 1.65 1.46 1.44 1.26
Origin Bruker BioSpin GmbH
Solvent DMSO
Temperature 297.3
Pulse Sequence zg30
Experiment 1D
Number of Scans 128
Acquisition Time 4.0894
Acquisition Date 2025-04-09T10:51:00
Spectrometer Frequency 400.17
Spectral Width 8012.8
Lowest Frequency -1538.5
Nucleus 1H
Acquired Size 32768
Spectral Size 65536
Cl
O
O
H
N
O
O
O
10
DMF
DMF
1.00 1.04 1.11 1.97 1.98 3.03 8.07 6.09 4.08 2.65 2.09 2.18 4.25
9.5 9.0 8.5 8.0 7.5 7.0 6.5 6.0 5.5 5.0 4.5 4.0 3.5 3.0 2.5 2.0 1.5 1.0 0.5 0.0
f1 (ppm)

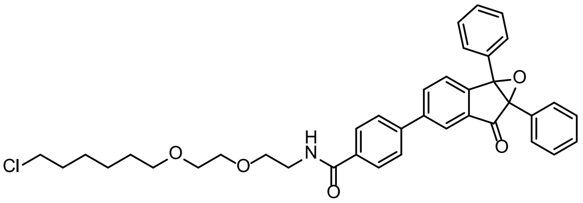
Cl
O
O
H
N
O
O
O

**10**

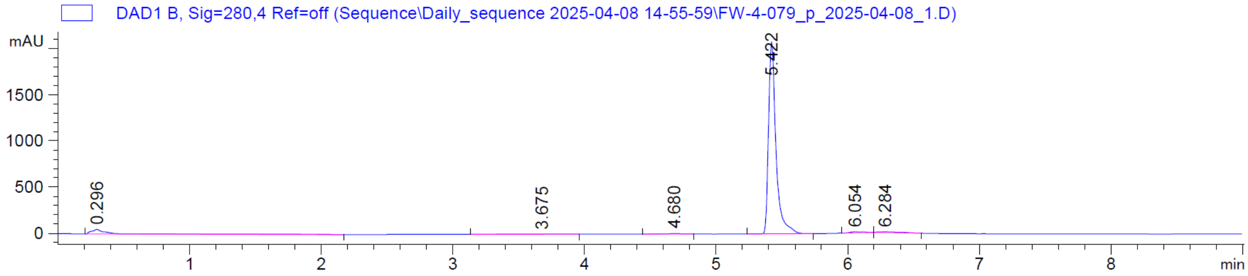
DAD1 B, Sig=280,4 Ref=off (Sequence\Daily_sequence 2025-04-08 14-55-59\FW-4-079_p_2025-04-08_1.D)
mAU
1500
1000
500
0
0.296
3.675
4.680
5.422
6.054
6.284
1 2 3 4 5 6 7 8
min

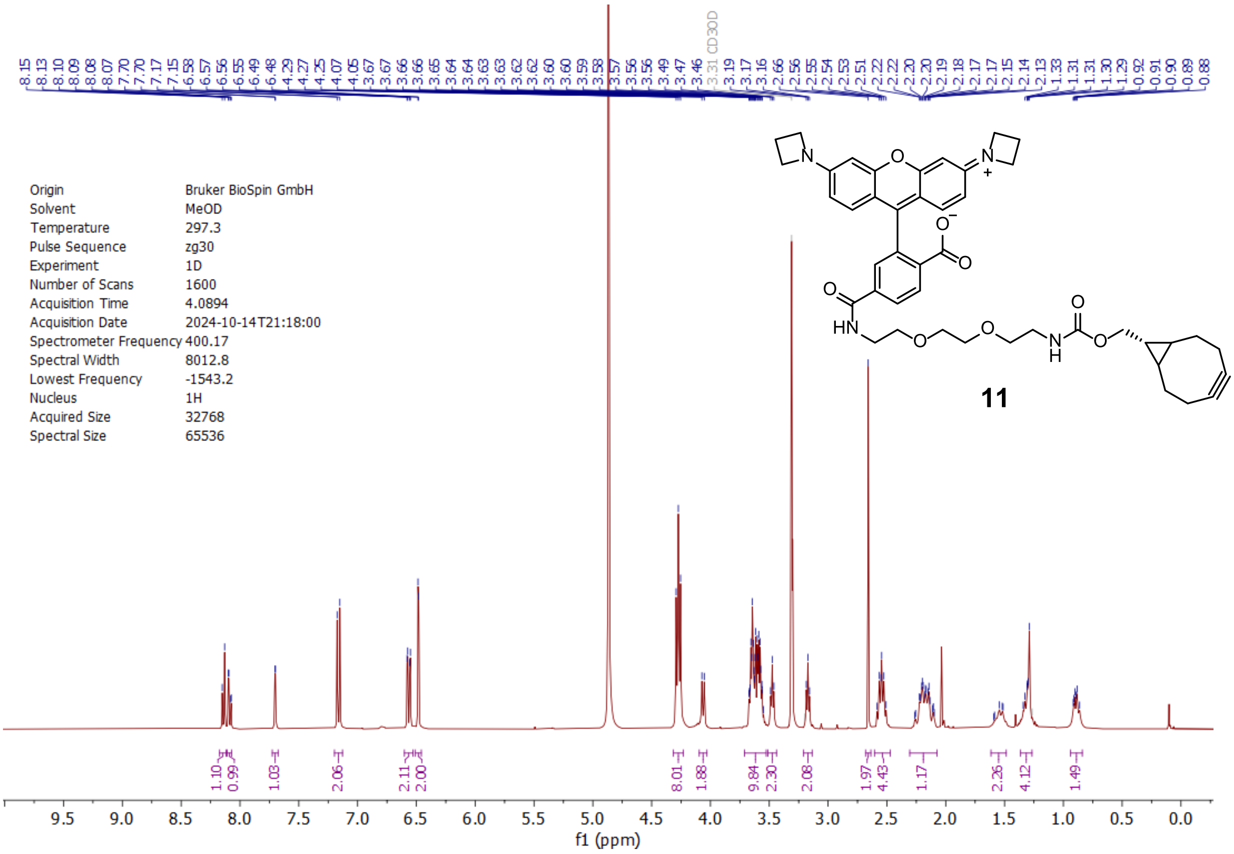

8.15 8.13 8.10 8.09 8.08 8.07 7.70 7.70 7.17 7.15 6.58 6.57 6.56 6.55 6.49 6.48 4.29 4.27 4.25 4.07 4.05 3.67 3.67 3.66 3.65 3.64 3.64 3.63 3.63 3.62 3.62 3.60 3.59 3.58 3.57 3.56 3.56 3.49 3.47 3.46 3.31 CD3OD 3.19 3.17 3.16 2.66 2.56 2.55 2.54 2.53 2.51 2.22 2.22 2.20 2.20 2.19 2.18 2.17 2.17 2.15 2.14 2.13 1.33 1.31 1.31 1.30 1.29 0.92 0.91 0.90 0.89 0.88
Origin Bruker BioSpin GmbH
Solvent MeOD
Temperature 297.3
Pulse Sequence zg30
Experiment 1D
Number of Scans 1600
Acquisition Time 4.0894
Acquisition Date 2024-10-14T21:18:00
Spectrometer Frequency 400.17
Spectral Width 8012.8
Lowest Frequency -1543.2
Nucleus 1H
Acquired Size 32768
Spectral Size 65536
11
1.10 0.99 1.03 2.06 2.11 2.00 8.01 1.88 9.84 2.30 2.08 1.97 4.43 1.17 2.26 4.12 1.49
9.5 9.0 8.5 8.0 7.5 7.0 6.5 6.0 5.5 5.0 4.5 4.0 3.5 3.0 2.5 2.0 1.5 1.0 0.5 0.0
f1 (ppm)

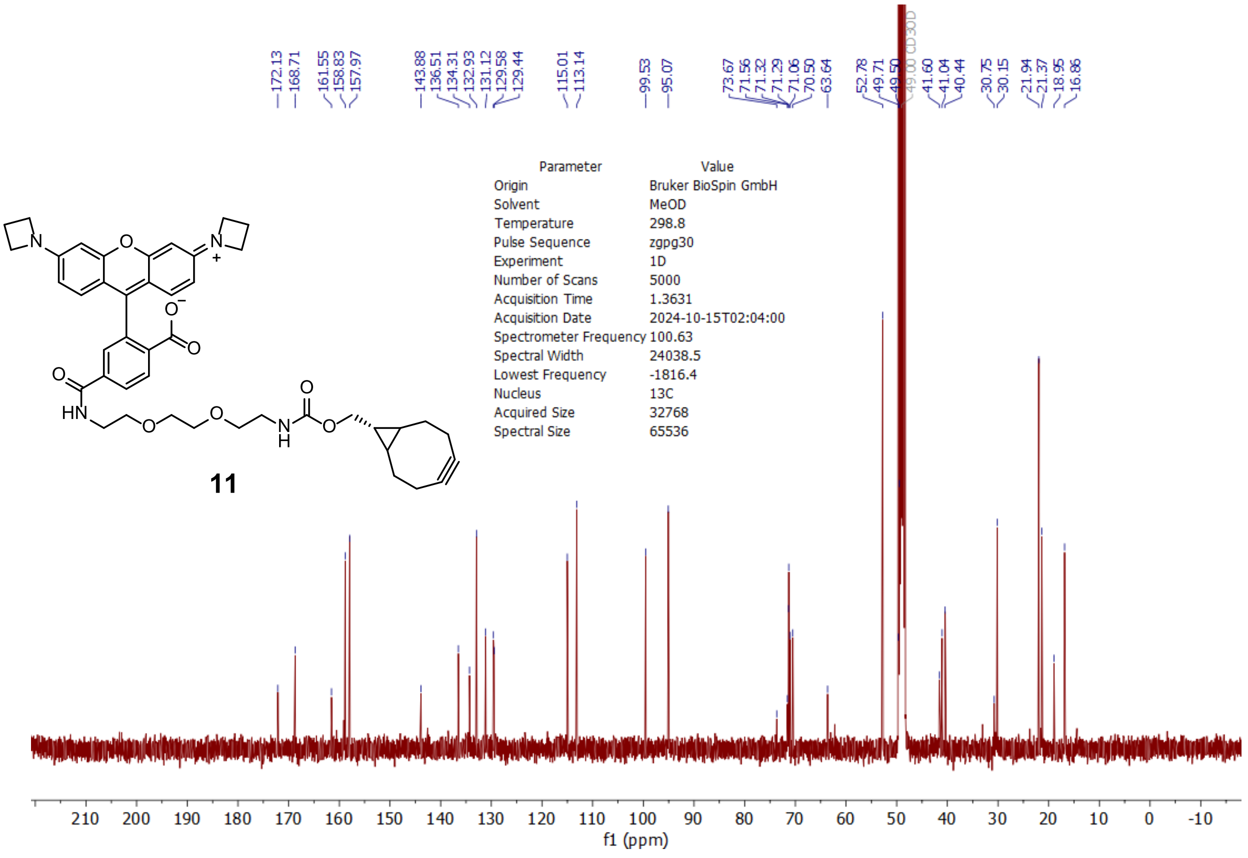

172.13 168.71 161.55 158.83 157.97 143.88 136.51 134.31 132.93 131.12 129.58 129.44 115.01 113.14 99.53 95.07 73.67 71.56 71.32 71.29 71.06 70.50 63.64 52.78 49.71 49.50 49.00 CD3OD 41.60 41.04 40.44 30.75 30.15 21.94 21.37 18.95 16.86
Parameter Value
Origin Bruker BioSpin GmbH
Solvent MeOD
Temperature 298.8
Pulse Sequence zgpg30
Experiment 1D
Number of Scans 5000
Acquisition Time 1.3631
Acquisition Date 2024-10-15T02:04:00
Spectrometer Frequency 100.63
Spectral Width 24038.5
Lowest Frequency -1816.4
Nucleus 13C
Acquired Size 32768
Spectral Size 65536
11
210 200 190 180 170 160 150 140 130 120 110 100 90 80 70 60 50 40 30 20 10 0 -10
f1 (ppm)
